# PHARP: A pig haplotype reference panel for genotype imputation

**DOI:** 10.1101/2021.06.03.446888

**Authors:** Zhen Wang, Zhenyang Zhang, Zitao Chen, Jiabao Sun, Caiyun Cao, Fen Wu, Zhong Xu, Wei Zhao, Hao Sun, Longyu Guo, Zhe Zhang, Qishan Wang, Yuchun Pan

## Abstract

Pigs not only function as a major meat source worldwide but also are commonly used as an animal model for studying human complex traits. A large haplotype reference panel has been used to facilitate efficient phasing and imputation of relatively sparse genome-wide microarray chips and low-coverage sequencing data. Using the imputed genotypes in the downstream analysis, such as GWASs, TWASs, eQTL mapping and genomic prediction (GS), is beneficial for obtaining novel findings. However, currently, there is still a lack of publicly available and high-quality pig reference panels with large sample sizes and high diversity, which greatly limits the application of genotype imputation in pigs. In response, we built the pig Haplotype Reference Panel (PHARP) database. PHARP provides a reference panel of 2,012 pig haplotypes at 34 million SNPs constructed using whole-genome sequence data from more than 49 studies of 71 pig breeds. It also provides Web-based analytical tools that allow researchers to carry out phasing and imputation consistently and efficiently. PHARP is freely accessible at http://alphaindex.zju.edu.cn/PHARP/index.php. We demonstrate its applicability for pig commercial 50K SNP arrays, by accurately imputing 2.6 billion genotypes at a concordance rate value of 0.971 in 81 Large White pigs (~ 17× sequencing coverage). We also applied our reference panel to impute the low-density SNP chip into the high-density data for three GWASs and found novel significantly associated SNPs that might be casual variants.

## INTRODUCTION

Over the last decade, because of the rapid development of high-throughput genotyping technologies, e.g., single nucleotide polymorphism (SNP) arrays (1), reduced-representation sequencing (RRS) (2,3) and whole-genome sequencing (WGS) (4), genome-wide association studies (GWASs) have detected thousands of loci associated with complex traits in animal (5) and human genomes. To date, considering the high genotyping cost of whole-genome sequencing for thousands of animals or more, the majority of GWASs still use low-density genotyping technologies (at tens of thousands of sites) such as SNP arrays or RRS. The GWASs based on the low-density SNP panels have been successful in terms of finding thousands of loci that have been statistically associated with risks for diseases and traits, and a large number of these loci are well replicated, indicating that they are true associations (6). However, because there are often many co-inherited variants in strong linkage disequilibrium (LD) with the most significant trait-associated variant (lead-SNP), the association of a locus with a disease/trait does not specify which variant at that locus is actually causing the association (i.e., the “causal variant”). As a consequence, a higher-resolution view of a genetic region obtained by adding more variants might be needed to determine which of the linked variants are functional. Thus, a high-density (1 ~ tens of millions of sites) genotypes are essential for GWASs, TWASs or eQTL mapping to provide deeper insights into disease/trait biology.

Genotype imputation is a more cost-efficient way to obtain a high-density genotype. Several imputation methods – e.g., BEAGLE (7), IMPUT2 (8), Minimac4 (9) and GLIMPSE (10) – have been developed to infer unobserved genotypes in one individual from the estimated haplotypes in a reference panel, which comprises a large number of markers. Genotype imputation could be beneficial (11) for fine mapping by increasing the chances of identifying a causal variant, meta-analysis by facilitating the combination of results across studies using different genotyping arrays, and increasing the power of association studies by increasing the effective sample size. Therefore, it has been widely used in genetic research, especially in humans (6), which usually involves genotyping SNPs in DNA genotyping microarrays (low-density) and then imputing genotypes at tens of millions of additional sites based on the availability of a large cohort of public haplotype reference panels (HRPs)., e.g., the 1000 Genomes Project (12) and the Haplotype Reference Consortium (HRC) (13).

As one of the most important livestock and animal models of human diseases, the genetic mechanisms of traits in pigs need to be dissected through fine mapping to locate the causal loci. Recently, pig haplotype reference panels have been begun to be used in pig genetic studies. For example, *Yan* et al (14) constructed a reference panel including 403 individuals from 10 populations and applied it to impute 60K SNPs (i.e., the PorcineSNP60 BeadChip) into whole-genome sequences of 418 Sutai individuals for a GWAS of lumbar number. They did not detect any significant signals using the original 60K SNPs but rediscovered the missing QTL for lumbar number in Sutai pigs using imputed-genotypes. Later, Yan et al also constructed a reference panel including 117 individuals and utilized it to impute 60K SNPs onto the whole-genome sequences of 1,020 individuals for a GWAS of haematological traits and found 87 novel quantitative trait loci (QTLs) for 18 haematological traits at three different physiological stages. Moreover, a previous study also suggested that using imputation-based whole-genome sequencing data can improve the accuracy of genomic prediction for combined populations of pigs (15). However, the pig reference panels in previous studies are either publicly unavailable or have a small sample size, which is the major determinant of genotype imputation accuracy (16,17). We only found one publicly available pig reference panel in Animal-ImputeDB (18), which includes a very small sample size (n = 233). With the increasing amounts of publicly available of pig genomic sequencing data (more than 25 terabytes), especially in the last five years, it is urgent to use these resources to construct a reference panel to facilitate a wide application of genotype imputation in pig genetic studies. Therefore, it is essential to develop a convenient database to provide a high-quality reference panel with a large sample size and good evaluation and imputation tools for pig genetic research.

Therefore, the aims of this study are to i) build the largest (in terms of sample size) reference panel of pigs to date and ii) provide a user-friendly online tool for efficient phasing and imputation of missing genotypes based on a high-quality reference panel.

## MATERIALS AND METHODS

In this study, we built a pig haplotype reference panel that integrates 1,006 individuals’ (**Supplementary Figure S1**) whole-genome sequence data from our laboratory (n = 84) and the NCBI SRA database (n = 922) that were publicly available (**Supplementary Table S1**), comprehensively evaluated its imputation performance, further developed an online imputation platform and applied it to GWASs and GSs (**Figure 1**).

**Figure 1.**
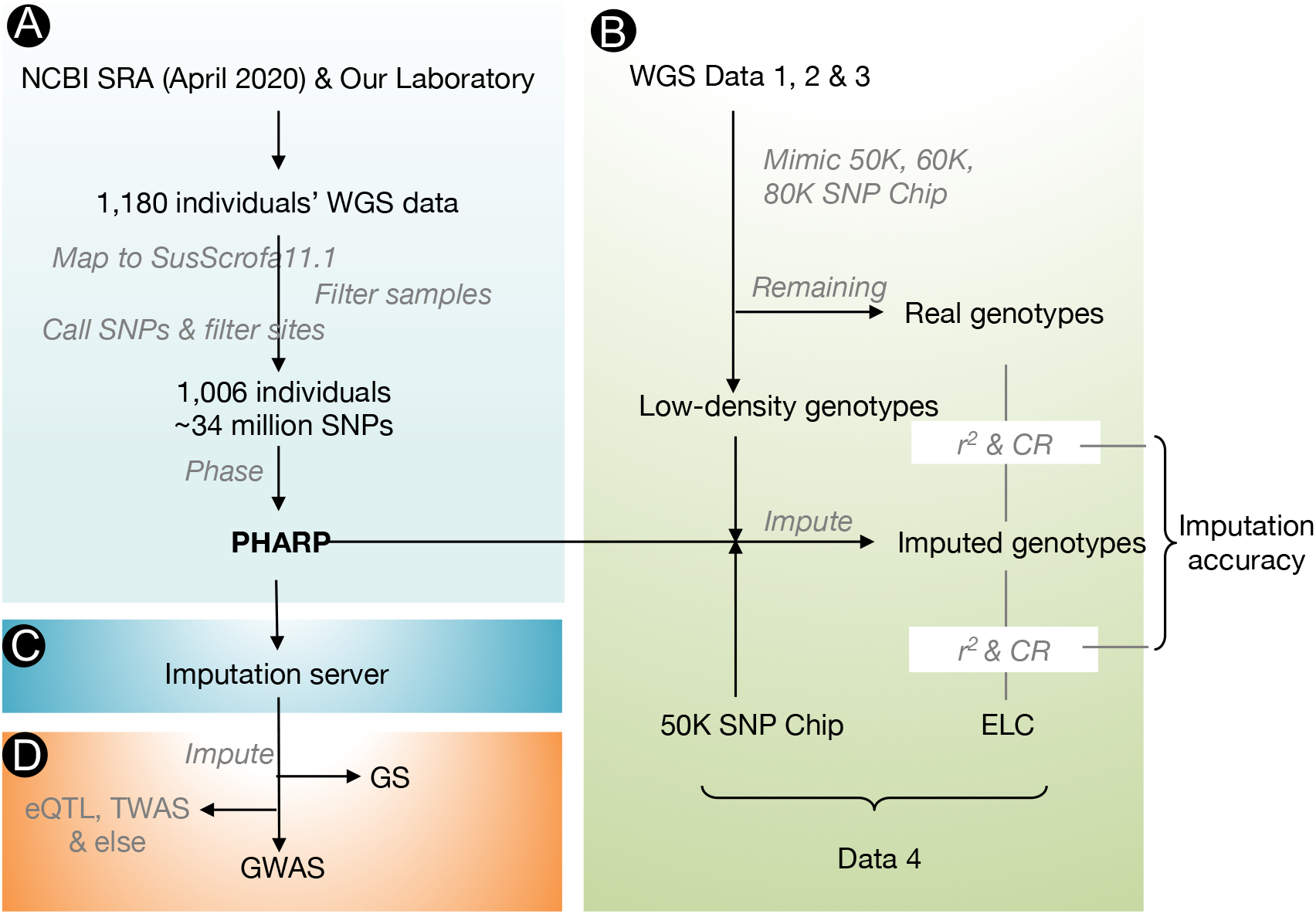
Schematic diagram of the pig haplotype reference panel’s construction, imputation accuracy evaluation, implementation platform and applications. A: Data resources and processing steps used to construct the PHARP. B: Imputation accuracy estimation of PHARP on multiple test datasets. C: Imputation platform development. D: Applications of PHARP in GWASs, GS and other potential studies such as eQTL mapping and TWASs.

### Datasets and their sources

#### WGS for PHARP building

From 52 pig whole genome sequencing projects collected in SRA (as of April 2020), we collected 1,097 individuals’ WGS data, including more than two thousands of experimental datasets (**Supplementary Table S1**). To increase the diversity of the reference panel, we also sequenced an additional 84 individuals’ genomes (including eight breeds). In total, 1,181 individuals’ WGS data from 71 populations were collected (**Supplementary Table S1**).

#### Test data for missing genotype imputation performance based on PHARP

The test data included (i) WGS data (n = 81, Large White pig breed, depth ranging from 9.7 ~ 38×) from PRJEB39374 and PRJEB38156; (ii) WGS data (n = 54, Jiaxinghei pig breed, depth ranging from 3.5 ~ 12×); (iii) WGS data (n = 299, Duroc pig breed, ELC depth ~ 1×); and (iv) individuals genotyped by both the 50K Chip and ELC (n = 20, Duroc pig breed, ELC depth ~ 1×) (**Supplementary Table S1** and **Figure S2**).

#### Genotypes and phenotypes data for the GWAS/GS

In total, 1,432 individual genotypes based on SNP arrays (i.e., the PorcineSNP50 BeadChip, the PorcineSNP60 BeadChip, or the Seek GGP Porcine 80K SNP chip) and phenotypes of 13 traits related to growth and fatness and reproduction in two pig breeds (Duroc and Sujiang) were collected from three previously reported GWASs (19–21) (see details in **Supplementary Table S2**).

### Construction of PHARP

#### Data processing and variant discovery

We used SRA Toolkit (https://github.com/ncbi/sra-tools) to download (`prefetch`) WGS data and convert (`fasterq-dump`) them from SRA to FASTQ format; performed quality control, read filtering and base correction for the raw FASTQ data by using fastp (22) with default parameters; mapped the high-quality reads to the latest version of the pig reference genome (Sscrofa11.1) using `BWA v0.7.17` (23) with the MEM function and the parameters for paired-end data; converted SAM files to BAM files and merged library data from individual and multiple experiments into one dataset using `samtools v1.10` (24); removed duplicated reads with `sambamba v0.7.1` (25); individually calculated coverage and depth with `Mosdepth v0.2.9` (26); and finally applied `GATK v4.1.6` (27) `HaplotypeCaller` to each sample to generate an intermediate GVCF, which was then used in `GenotypeGVCFs` for joint genotyping across all samples.

#### Sample filtering

We first removed samples that could not be successfully converted from SRA to FASTQ format (n = 5, PRJEB29465) and that had a depth less than 4× (n = 166), as suggested by Jiang et al., (28) to reduce false-positive variant detection in pigs. To detect possible duplicates, we counted the number of genotypes (measured by Euclidean distance using the `dist` function in R) that differed between each sample pair using the original genotypes of SNPs after pruning by `PLINk v1.9` (`--indep-pairwise 50 5 0.2`) (29,30). We identified four sample pairs with Euclidean distance outlier (extremely low values less than 2) as duplicates and removed one of the samples in each pair as described in Supplementary Figure S3. These filters resulted in a total of 1,006 samples being used for the final phased reference panel.

#### Site filtering

After sample filtering, we filtered SNPs with the following criteria: (1) “QD < 2.0, FS > 60.0, MQ < 40.0, MQRankSum < −12.5, ReadPosRankSum < −8.0, SOR > 3.0”; (2) a minor allele frequency (MAF) < 0.01; and (3) a call rate < 0.9 `using GATK VariantFiltration`. After applying these filters, in total 34,135,654 SNPs of autosomes were retained in the final site list.

#### Pre-phasing

We applied the `SHAPIT v2` (31) method to individually pre-phase the called genotypes for each autosome, as it was reported that the rephrasing approach substantially improved imputation accuracy when using the haplotypes (16).

### WGS processing of test data

For each genotype imputation performance test dataset from WGS, we used the same procedure and filtering steps as described above in ‘Construction of PHARP’ to obtain their genotypes except i) without MAF filtering and ii) keeping only the site within our reference panel.

### Imputation performance estimation

We used the four experiments data sets (related to the three pig breeds) mentioned above to assess the imputation accuracy performance of our reference panel. Imputation was carried out using `Minmac4` (9), in which parameters were set to default values. The reference/target panel was pre-phased by `SHAPIT v2` (31). We used two measures to evaluate the imputation accuracy: (i) the concordance rate (CR), which is calculated as the percentage of genotypes imputed correctly among the total imputed genotypes, and (ii) *r^2^*, a correlation-based measure, which is the squared correlation between the true and imputed doses of an allele across all imputed samples. Five scenarios of genotype imputation performance estimations are given below.

#### Mimic chips

To mimic a typical imputation analysis, we created three target panels on the basis of three chip lists: the Illumina PorcineSNP50 Genotyping BeadChip (50K), the Illumina PorcineSNP60 v2 Genotyping BeadChip (60K), and the GeneSeek GGP Porcine 80K SNP chip (80K). These pseudo-chip genotypes (from test datasets 1, 2, and 3) were pre-phased by `SHAPIT v2` before being used to impute the remaining genotypes, which were then compared to the held-out genotypes.

#### Density of the genotyping array

To estimate the general imputation accuracy affected by genotype array density, we also created target panels (from test datasets 1, 2, and 3) with a gradient density of sites as follows: we divided autosomes into bins according to the physical location at a specific length (we set up bins with a length of 2.5, 5, 10, 20, 40, 50, 60, 80, 100, 200, and 400 kb), and randomly selected one site from each bin. We used the selected sites to compose new pseudo-chip genotypes, which were then used to impute the genotypes of unselected sites. We then compared the unselected site genotypes between imputed and observed (real) data and repeated the above pseudo-chip genotypes imputation at each specific density five times.

#### Size of the reference panel

Generally, increasing the size of the reference panel will increase the imputation accuracy, as a larger panel provides a larger set of template haplotypes to match against. We randomly selected 10, 30, 50, 100, 200, 400, 600, 800, and 900 samples from PHARP to construct a subset reference panel (repeated 5 times). Test dataset 4 and test dataset 1 mimicking a 50K chip were used to estimate the imputation accuracy.

#### A breed subset of PHARP

We built two reference panels, i.e., a subset of the Large White (n = 115) and Duroc (n = 85) pig breeds with the largest sample sizes from PHARP, to compare the imputation accuracies of different reference panels. The imputation accuracy was estimated the same way as mentioned in the ‘ Size of the reference panel’ section using the same test data sets.

#### Comparison with Animal-ImputeDB

To date, we have only found one publicly available pig reference panel from Animal-ImputeDB (18), containing 233 samples. To keep the assembly of our test dataset 4 consistent with Animal-ImputeDB (based on Sscrofa10.2), we used `liftOver` (the chain file was downloaded from http://hgdownload.soe.ucsc.edu/goldenPath/susScr11/liftOver/) to convert the genome coordinates of test dataset from Sscrofa11.1 to Sscrofa10.2. We conformed their chromosome strand and allele order matched those of the Animal-ImputeDB reference panel using `conform-gt.jar`. We only used sites consistent between the reference panels to enable direct comparison. These 50K genotypes of 20 samples were used to impute the remaining genotypes, which were then compared to the ELC genotypes of the same samples.

### System design and implementation

The current version of PHARP was developed using MySQL 5.7.27 (http://www.mysql.com) and runs on a Linux-based Apache Web server. PHP 7.0.33 (http://www.php.net/) is used for server-side scripting. We designed and built the interactive interface using layui (https://www.layui.com/) with the HTML, CSS and JS frameworks on the Web. We recommend using a modern Web browser such as Google Chrome (preferred), Firefox, or Safari to achieve the best display effect.

### Applications of PHARP

To illustrate the benefits of using the PHARP resource, we applied it to three previously reported GWASs as mentioned in the ‘Data sets and their sources’ section. For each study, we first pre-phased individuals by `SHAPIT v2` (31) using autosomal SNPs after filtering out SNPs with a MAF < 0.01 and a call rate < 0.9. SNPs were also removed if they could not be remapped to the SusScrofa11.1 pig reference genome. We next remotely performed imputation using our PHARP imputation server (http://alphaindex.zju.edu.cn/PHARP/index.php), and subsequently applied it to imputation data sets with an imputation quality threshold of *R^2^* > 0.4 before association testing.

#### GWAS

To make the association results comparable before and after imputation, we adopted the same association model and significance threshold as described in each original paper (19–21). Association analysis using a single marker regression model was implemented in `GEMMA v.0.98.1` software (32). The association plots were produced using the R ggplot2 package.

#### GS

Datasets included in the aforementioned GWAS (19) was also used to carry out genomic prediction based on different SNP datasets. The fivefold cross-validation scheme was adopted to evaluate prediction accuracy. Briefly, the individuals were randomly separated into five groups with the same sample size, and each time, the genomic estimated breeding values (GEBVs) of one tested group were predicted using the phenotypic information of the other four groups and genomic information of all individuals with the following genomic best linear unbiased prediction (GBLUP) model:

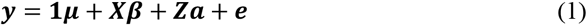

where ***y*** is the phenotypic vector; ***μ*** is the overall mean; ***β*** is the fixed effect used in the original studies; and ***a*** is a vector of breeding values that is assumed to follow the normal distribution 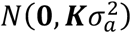, where 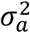 is the additive genetic variance and ***K*** is the genomic relationship matrix (GRM) built from three different SNP datasets using GCTA (33) (*--make-grm-alg 1*), including (i) chip SNPs from the original studies (GBLUP_CHIP); (ii) imputed SNPs (GBLUP_IMP); and (iii) LD-pruned (r^2^ < 0.5) imputed SNPs (GBLUP_PRUNE) using PLINK (*--indep-pairwise 50 5 0.2*). ***e*** is a vector of the residual variance following normal distribution 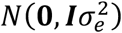, where ***I*** is the identity matrix and 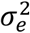 is the residual variance. **1, *X*** and ***Z*** are incidence matrices. The variance components and GEBVs were estimated using the BGLR package (34), and the prediction accuracy was calculated as the Pearson’s correlation coefficient between the GEBVs and true breeding values (TBVs), where TBVs were estimated based on the full dataset using the GBLUP model with the GRM built from the chip SNP dataset as used in Xu et al (19). The fivefold cross validation was repeated ten times.

## RESULTS

### Imputation evaluation

To mimic a typical imputation analysis, we first created three datasets by extracting high-coverage whole-genome sequencing genotypes for 81 Large White pigs at all sites included in the most popular commercial porcine microarray genotyping platform (50K, 60K, and 80K). These were used to impute the remaining genotypes, which were then compared to the held-out genotypes. The imputation accuracy as assessed by the CR value reached 0.971 to 0.978 (with a *r^2^* value from 0.920 to 0.939, **Figure 2A** and **Supplementary Table S3**), indicating that PHARP can be well applied to low-density genotypes from the current popular commercial porcine microarray genotyping platform for imputation with high accuracy performance. We also performed the same imputation accuracy test on the extremely low-coverage whole-genome sequencing genotypes for 299 Duroc pigs. Although the CR value estimated from the Duroc panel decreased slightly compared to that for the Large White panel, it was still greater than 0.9 (CR = 0.934 to 0.947, *r^2^* = 0.876 to 0.901, **Figure 2A** and **Supplementary Table S3**). We speculated that this might be because the low coverage (~ 1×) probably result in a certain proportion of heterozygotes being falsely genotyped, causing incorrect haplotype inference and in turn reducing the imputation accuracy. Finally, we attempted to investigate the imputation performance of PHARP on pig breeds uncovered in our panel using a middle-coverage (average of depth was 6.3×) whole-genome sequencing genotypes for 54 JiaXingHei pigs (JXH, a Chinese indigenous pig breed). The CR value of JXH decreased to approximately 0.81 (**Figure 2A** and **Supplementary Table S3**, *r^2^* was approximately 0.5), which was expected because low genetic similarity between the reference and target panels may decrease imputation accuracy.

**Figure 2.**
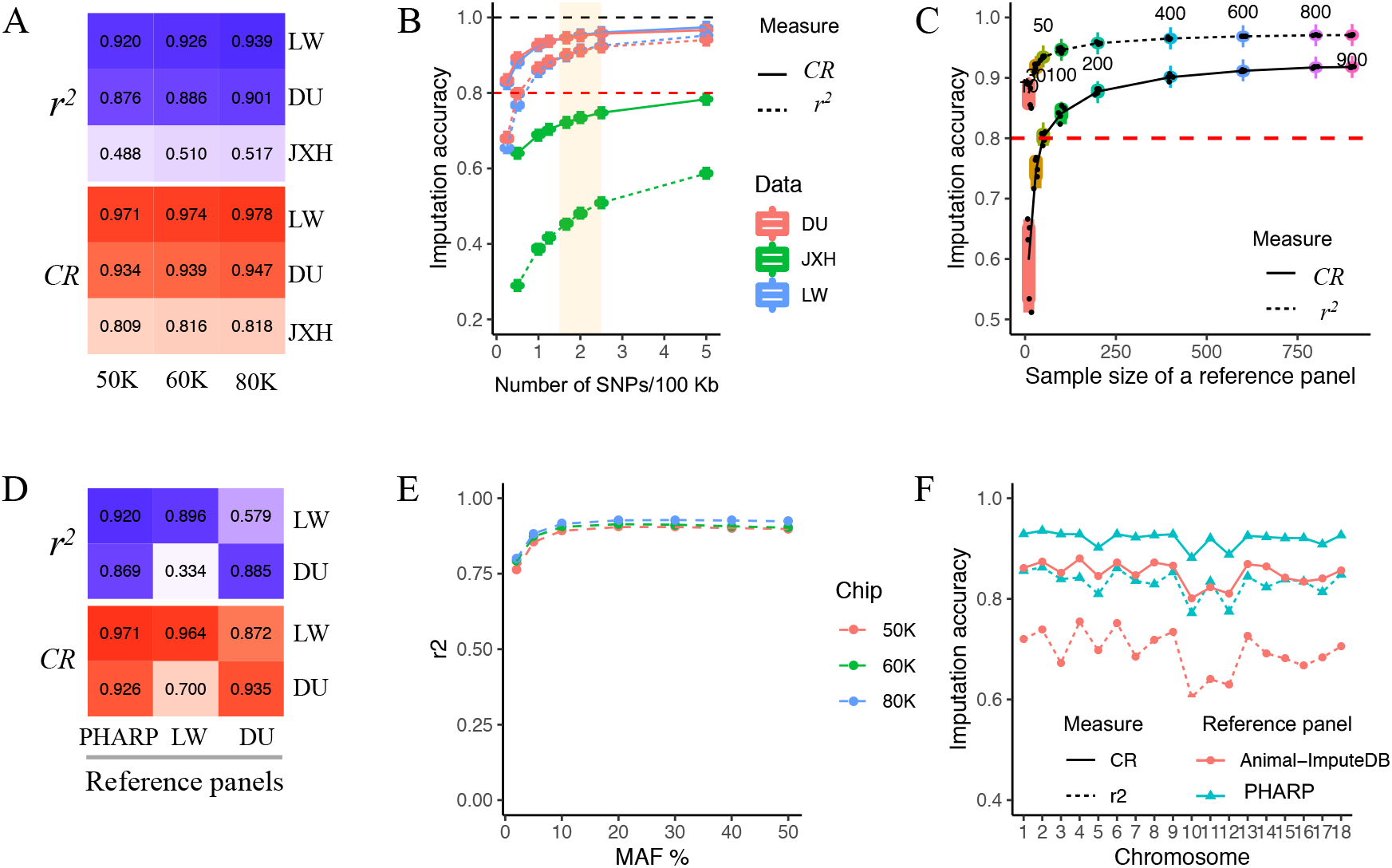
Imputation accuracy under different scenarios. A: Mimicing three popular pig commercial chips (50K, 60K, and 80K) using three datasets by masking all variants (only autosomes were used) except those on the chips; the held-out genotypes were considered as ‘real’ to calculate the CR and *r^2^* values. B: Boxplot of imputation accuracy estimated by mimicking the target panel with different densities of SNPs on chromosome 1 using test datasets 1, 2 and 3 (See **Supplementary Figure S4** for plots of the remaining autosomes). C: Boxplot of the imputation accuracy estimated by mimicking 50K chip genotypes from dataset 1 using different sizes of reference panels constructed by randomly extracting samples from 1,006 individuals (repeated 5 times). D: Mimicking the 50K chip genotypes from dataset 1 and 2 and using reference panels constructed by extracting samples according to pig breed (LW, Large White, n = 114; DU, Duroc, n = 85). E: The imputation accuracies of the different MAF bins ((0, 0.02], (0.02, 0.05], (0.05, 0.1], (0.1, 0.2], (0.2, 0.3], (0.4 0.5]) estimated by mimicking the 50K chip genotypes using dataset 1. F: The imputation accuracy estimated from dataset 4 using our reference panel and that from Animal-ImputeDB. Dataset 1, Large White pig breed, LW, n = 81; dataset 2, Duroc pig breed, DU, n = 299; dataset 3, Jiaxinghei pig breed, JXH, n = 54; dataset 4, Duroc pig breed, n = 20, pigs were genotyped by both a 50K chip and ELC.

We next estimated imputation accuracy in a more general scenario by testing a wide range of densities of SNPs in target panels. As expected, the *r^2^* value increased with a denser SNPs in the target panels. Specifically, it dramatically increased before a density of 60 kb per SNP, achieved an average of more than 0.8 (varying from 0.755 to 0.903 in Large White pigs and 0.840 to 0.902 in Duroc pigs, replicated 5 times, **Figure 2B**, **Supplementary Figure S4** and **Table S4**) for the majority of autosomes, and achieved an average of greater than 0.8 (varying from 0.817 to 0.931 in Large White pigs and 0.876 to 0.925 in Duroc pigs) for all autosomes with a density of 40 kb per SNP. Interestingly, the densities of commonly used commercial porcine SNP arrays (e.g., 50K, 60K, and 80K) are exactly between 40 and 60 kb per SNP and are approximately 50 kb per SNP, suggesting that PHARP has wide applicability to the current porcine SNP arrays for imputation and can achieve high imputation accuracy.

We then explored factors, such as the size and breed subsets of the reference panel and the minor allele frequencies (MAFs) of variants in the reference panel, that can affect imputation accuracy. The *r^2^* value increased with a larger sample size in the reference panels, which is expected because a larger panel provides a larger set of template haplotypes to match against, which improves imputation accuracy. It grew slowly after the size increased to 600 (average of CR = 0.969 and 0.919, *r^2^* = 0.912 and 0.856, in Large White and Duroc pigs, respectively, when the size of the reference panel was 600, **Figure 2C**). Moreover, we examined imputation performance using two subsets of reference panels constructed for the Large White pig breed (n = 114) and the Duroc pig breed (n = 85). We observed that the *r^2^* value was greater when using PHARP (CR = 0.971, *r^2^* = 0.920, **Figure 2D**) than when using only Large White pigs as a reference panel (CR = 0.964, *r^2^* = 0.896, **Figure 2D**), estimated by mimicking 50K SNP array using the test dataset 1 (81 Large White pigs). However, the *r^2^* value obtained using the PHARP (CR = 0.926, *r^2^* = 0.869, **Figure 2D**) was slightly less than obtained using the reference panel (CR = 0.935, *r^2^* = 0.885, **Figure 2D**) including only Duroc pigs estimated from using the test dataset 4 (20 Duroc pigs). Note that for 20 Duroc pigs, ELC-WGS (~ 1×) genotypes were used as ‘real’ genotypes to calculate the imputation accuracy, which might have underestimated imputation performance because the ELC-WGS probably caused heterozygotes to be falsely genotyped due to the limitation of the extremely low sequencing depth. We also investigated imputation accuracy under different MAF bin sizes. Taking the imputation accuracy estimation of the mimic 50K SNP array from the Large White target panel as an example, the *r^2^* value surpassed 0.89 when the MAF of variants was greater than 0.05 and reached 0.762 even when the MAF was less than 0.02 (**Figure 2E**). Imputation accuracy increase in each bin of the MAF for the other two denser mimic SNP arrays (60K and 80K, **Figure 2E**) in comparison with the 50K SNP array.

We finally compared our reference panel with that from Animal-ImputeDB, which is the only one publicly available pig reference panel. Test dataset 4 was used to evaluate performance accuracy. The results showed that PHARP achieved a CR value of 0.921 (*r^2^* = 0.836), whereas Animal-ImputeDB had a CR value of 0.854 (*r^2^* = 0.7, **Figure 2F**), suggesting that PHARP greatly increased the imputation accuracy.

### PHARP imputation server

We developed a user-friendly website to provide an imputation service (http://alphaindex.zju.edu.cn/index.php, **Supplementary Figure S5**) using the PHARP for researchers, which provides the imputation process for a genotype data in variant call format (VCF) (35). The imputation pipeline includes four main steps: pre-processing, phasing, imputation, and post-processing. The pre-processing step for uploaded files consists of checking their format and content validity, summarizing their basic information such as sample size, and modifying their records to be consistent with our reference VCF file. After that, the pre-processed data are phased using `SHAPIT v2` (31) or `Beagle v5.1` (7) without a reference panel. Then the imputation is carried out with `Minmac4` (9). In the final post-processing step, the output is evaluated and provided as bgzip-compressed VCF file. Users will be notified by email after the imputation is completed and the result will be stored on the server for two weeks.

### Applications

To illustrate the benefits of using the PHARP resource, we imputed GWASs of 1,432 samples from three studies. This analysis highlighted potential new associations at the genome-wide suggestive significance threshold of *P*-value < 1/the number of independent markers. For example, for the backfat thickness phenotypes, *Xu et al*. and *Zhang et al*. reported two (*XKR4* and *PENK*) and nine (such as *GRM4, SNRPC, TSHZ1* and *PHLPP1*) associated genes, respectively. Using the PHARP-imputed genotypes, we found novel genes, such as *ANGPTL2, CCL8, TNXB, MC4R, PACSIN1*, and *MLIP* (**Figure 3A and B**, see more results for other phenotypes in Supplementary Table S2). We also found that it is possible for PHARP-based imputation to refine signals of association. For example, the association results using PHARP-based imputation for the backfat thickness phenotype at the *SNRPC, GRM4* and *PACSIN1* loci are shown in **Figure 3C-E**. At the *SNRPC* locus, the original paper reported that MARC0061142 was the most significant SNP (*P*-value = 5.42 × 10^-8^) associated with the backfat thickness phenotype. We found a more significant SNP (chr7:30719820:G:A, *P*-value = 5.42 × 10^-8^) with a tight linkage with this SNP (r^2^ = 0.99), implying a potential causal variant. Similarly, at the *GRM4* and *PACSIN1* loci, we found one (chr7:30191165:T:C, *P*-value = 1.19 × 10^-7^, r^2^ = 0.98) and two (chr7:30509485:C:G and chr7:30509543:C:G, *P*-value = 8.91 × 10^-8^, r^2^ = 0.98) missense variants, potential casual variants, with high linkages with their most significant SNPs (chr7:30250983:G:C, *P*-value = 6.08 × 10^-8^, and chr7:30497108:T:C, *P*-value = 1.44 × 10^-8^, **Figure 3D-E**), respectively.

**Figure 3.**
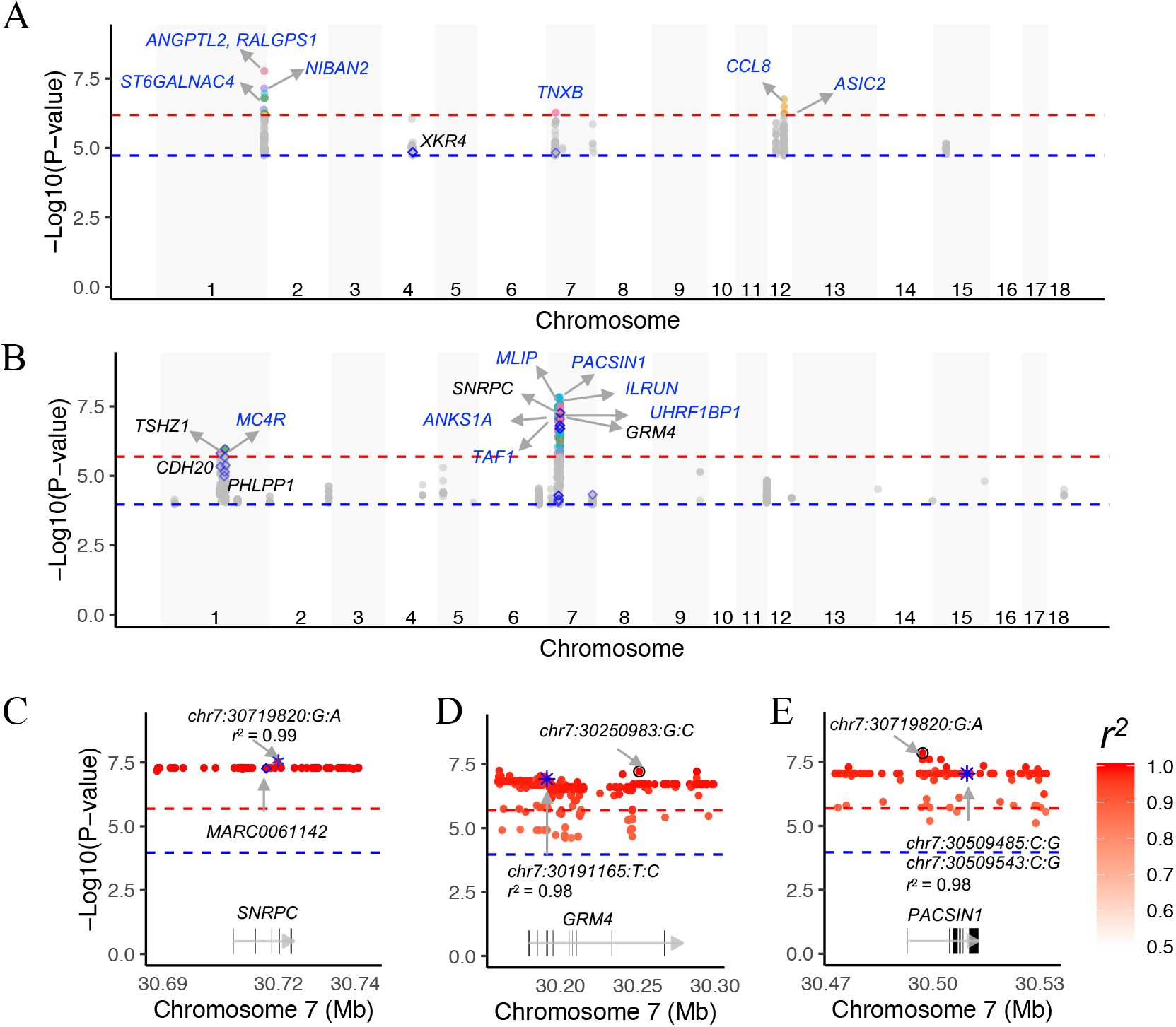
Association signals for growth phenotypes before and after imputation. Association test statistics on the –log10 (*P-value*) scale (y-axis) are plotted for each SNP position (x-axis) for the trait of backfat thickness at an age of 180 days (A), from Zhang et al., and at 100 kg (B), from Fu et al. To simplify the plot, only the variants with a *P-value* less than 1.08×10^-4^ are shown, and they are colored according to the annotated genes. The black-labeled genes are reported in the original paper, and the blue-labeled genes are novel genes detected after imputation. Examples of potential causal variants (marked by blue asterisks) in the *SNRPC* (C), *GRM4* (D) and *PACSIN1* (E) genes. Each dot represents a variant, whose LD (*r^2^*) with the Chip SNP (marked by blue diamonds) or the one with the lowest *P-value* (marked by a black circle) is indicated by the colour of the dot. The two horizontal lines divide SNPs with *P-values* < 2.05×10^-6^ and < 1.08×10^-4^ (A), and *P-values* < 6.46×10^-7^ and < 1.86×10^-5^ (B).

We also carried out genomic prediction using the chip and the same imputed genotypes and phenotypes as in the above GWASs. GBLUP_IMP and GBLUP_PRUNE had higher prediction accuracies across most traits (except for CW) tested in the datasets of Xu et al. (**Supplementary Table S5**).

## DISCUSSION

The first release of the PHARP is the largest pig genetic variation resource thus far with enriched ancestral diversity and has been created by combining data from many different studies. We searched the NCBI SRA database and identified WGS-based studies of pigs to collect together as many whole-genome sequencing datasets as possible and joined them with WGS data from our laboratory to build a much larger combined haplotype reference panel. By doing so, we provide a single centralized resource for pig genetics researchers to carry out genotype imputation.

PHARP achieves high imputation performance. We systematically estimated the imputation performance of PHARP using multiple test datasets. First, PHARP is able to accurately impute porcine commercial SNP array chips (e.g., 50K, 60K and 80K) for the most represented pig breeds worldwide, such as the Large White and Duroc pig breeds. As test dataset 1 includes high-coverage (with an average depth of 17×, Large White pig breed) whole-genome sequencing, implying high-quality genotyping, we do expect the imputation performance estimated from these data to be more reliable and find that the imputation accuracy as assessed by CR value (mimicking a 50K chip) surpasses 0.97 (*r^2^* > 0.92), suggesting high imputation accuracy. Compared to the test dataset 1, the test dataset 2 (with an average depth of 1×, Duroc pig breed) had a slightly decreased imputation accuracy (mimicking a 50K chip, CR = 0.93, *r^2^* = 0.88), possibly caused by a low density of SNPs (sites covered by the 50K chip that were kept after quality control were less abundant than those in test dataset 1) and false genotyping of heterozygotes resulting from low coverage. We also investigated the imputation performance for a pig breed (JXH) that is not covered in PHARP and found that, as expected, imputation accuracy decreased (mimicking a 50K chip, CR = 0.81, *r^2^* = 0.49) because of low genetic similarity between the reference and target panels. To overcome this limitation, we will substantially increase the ancestral diversity of the panel by sequencing more pig breeds in the future. Second, imputation accuracy increases with an increasing SNP density in the target panel and grows slowly after the SNP density surpasses 60 kb per SNP (**Figure 2B**), implying that the SNP density of the most popular SNP array chips (e.g., 50K, 60K and 80K, with SNP densities of 40 ~ 60 kb per SNP) might be enough to achieve high imputation accuracy for these imputed sites. Third, PHARP is able to accurately impute genotypes of rare variants. The *r^2^* value is still high under a low MAF ((0,0.02], CR = 0.996, *r^2^* = 0.76; (0.02,0.05], CR = 0.99, *r^2^* = 0.85; mimicking the 50K chip, test dataset 1). Fourth, PHARP has a better imputation performance than the publicly available pig reference panel in Animal-ImputeDB. The imputation accuracy as assessed by the CR *r^2^* value could be improved from 0.85 (Animal-ImputeDB, *r^2^* = 0.7) to 0.93 (PHARP, *r^2^* = 0.84) (test dataset 4, 20 Duroc pigs, 50K), probably because of the large increase in the sample size in PHARP (n = 1,006). We are planning to sequence decades of breeds that are not included in the first release of PHARP and add more pig WGS data that are publicly available worldwide to enlarge the ancestral diversity and sample size. Therefore, we expect to be able to make future gains in imputation performance.

We have developed centralized imputation server resources to enable pig genetic researchers to easily carry out imputation, which will greatly facilitate the application of reference panels in imputation. Users simply upload phased or unphased genotypes, and imputation is carried out on online servers. The users will be alerted by email once imputation is completed, at which time they can download the imputed datasets.

We demonstrated the good application of PHARP in pig genetic studies. *Increasing the power of association studies*. We could replicate genes well known to be to highly related to a specific trait. For example, the melanocortin-4 receptor (*MC4R*) gene is reported to be related to fatness and growth traits in pigs (36). However, Zhang et al. failed to detect this signal in D100 and L100 (two phenotypes of fatness and growth traits). Interestingly, we were able to rediscover this gene using their imputed-genotypes for those two phenotypes (**Supplementary Table S2**). Moreover, we found novel candidate genes associated with a specific trait. For example, using the imputed-genotypes from Xu et al., we detected the Angiopoietin-like protein 2 (*ANGPTL2*) gene, with the most significant *P* value (1.68 × 10^-8^, **Figure 3A**) for backfat thickness. A previous study reported that *ANGPTL2* treatment can induce lipid accumulation and increase fatty acid synthesis and lipid metabolism-related gene expression in mouse liver (37), suggesting that this gene is an important candidate gene associated with backfat thickness. *Fine mapping*. Imputation increases the number of variants and thus provides a higher-resolution view of a genetic region, thereby increasing the chances of identifying a causal variant. We were able to find functionally annotated variants, such as missense variants, implying a potential causal variant, by pinpointing whether they had an LD score with the most significant SNPs at a locus (**Figure 3D-E**). *Increasing the prediction accuracy of specific traits in GS*. The genomic prediction accuracies obtained using SNP datasets at different densities (SNP chip, WGS or imputed WGS) were compared in several previous studies (38–40), and generally, the sparse chip SNPs were sufficient enough to obtain accurate predictions. The reason might be that the inclusion of many SNPs could also induce more noise, and the sample size did not match the large number of SNPs. However, the prediction accuracies obtained using imputed WGS data will be higher for some traits. We observed this for the traits in the datasets of Xu et al. Some studies also showed that genomic prediction based on LD-pruned WGS-level SNP datasets could result in higher prediction accuracy (41), which could be observed for the dataset of Xu et al., where the pruned SNP dataset had the highest prediction accuracies for most traits (**Supplementary Table S5**). In addition, PHARP might be of great value if we could integrate the results of GWASs based on imputed datasets into genomic prediction using genomic feature BLUP (42) or trait-associated BLUP (43); such a strategy has been validated in several studies (44,45). We did not validate this strategy because the sample size of the test dataset is too small.

There is an additional potential utilization of PHARP. *Meta-analysis*. PHARP can aid in meta-analysis by generating a common set of variants that can be analysed across multiple studies to boost power. Different studies often use difference genotyping platforms (SNP arrays or GGRS), resulting in a small proportion of shared variants. For example, less than 10% of the SNPs included on the 50K SNP array are included in GGRS. *Genome-wide eQTL mapping*. Gene expression levels measured by high-throughput technologies, such as RNA-Seq, are treated as quantitative traits. Genotypes are also called using the RNA-Seq data from the same set of individuals and can be imputed to the WGS level, and statistical analyses are performed to detect associations between imputed markers and expression traits. *Genetic resource identification*. PHARP includes an enriched ancestral diversity that can be used as a control for clustering analysis. PHARP data can be applied to easily pinpoint an unknow individual/population as a novel genetic resource or identify which pig breed it is closely related/similar by doing cluster analysis such as the NJ-trees construction.

## CONCLUSIONS

In summary, we generated a large-scale reference panel for pigs, which will be a highly valuable resource for resolving the deficiency of large-sample-size pig genomic data. We believe that our efforts will markedly contribute to improving the genotype imputation accuracy in pigs, and ultimately facilitate genomic research of variants and their roles in pig complex traits.

In the future, we envisage the reference panel increasing in size and consisting of samples from a more diverse set of breeds. On the one hand, our group will genotype thousands of individuals from more than 100 pig breeds using WGS in the upcoming one/two year and is cooperating with other pig genome research groups to include their WGS data. On the other hand, more pig WGS data are becoming publicly available. Thus, we expect to greatly enlarge the sample size of PHARP, which should lead to further gains in imputation performance, especially for rare variants. Moreover, we will add small indels into the reference panels and continue to update PHARP.

## Supporting information

PHARP-Supplementary-Tables and Figures

## ACKNOWLEDGEMENTS

We thank Zhe Zhang of the College of Animal Science at the South China Agricultural University, Guangzhou, for their sharing the GWAS data and expert advice on the GWAS.

## AUTHOR CONTRIBUTIONS

Y.P., Z.W., Q.W. conceived and designed this study. Z.W. and Z.Z. wrote the manuscript and evaluated the imputation performance. Z.W. and Z.Y.Z collected and processed WGS data to obtain genotypes, and designed and developed PHARP online server. Z.Z. and W.Z. performed the GS analysis. Z.C. and Z.W. conducted the GWAS analysis. H.S., Z.X., J.S., C.C., F.W. and Q.W. provided useful input for the analyses and helped edit the manuscript. All authors read and approved the final manuscript.

## REFERENCES

1. LaFramboise, T. (2009) Single nucleotide polymorphism arrays: a decade of biological, computational and technological advances. Nucleic Acids Research, 37, 4181–4193.

2. Poland, J.A., Brown, P.J., Sorrells, M.E. and Jannink, J.L. (2012) Development of High-Density Genetic Maps for Barley and Wheat Using a Novel Two-Enzyme Genotyping-by-Sequencing Approach. Plos One, 7.

3. Chen, Q., Ma, Y.F., Yang, Y.M., Chen, Z.L., Liao, R.R., Xie, X.X., Wang, Z., He, P.F., Tu, Y.Y., Zhang, X.Z. et al. (2013) Genotyping by Genome Reducing and Sequencing for Outbred Animals. Plos One, 8.

4. Giani, A.M., Gallo, G.R., Gianfranceschi, L. and Formenti, G. (2020) Long walk to genomics: History and current approaches to genome sequencing and assembly. Comput Struct Biotec, 18, 9–19.

5. Hu, Z.L., Park, C.A. and Reecy, J.M. (2019) Building a livestock genetic and genomic information knowledgebase through integrative developments of Animal QTLdb and CorrDB. Nucleic Acids Research, 47, D701–D710.

6. Welter, D., MacArthur, J., Morales, J., Burdett, T., Hall, P., Junkins, H., Klemm, A., Flicek, P., Manolio, T., Hindorff, L. et al. (2014) The NHGRI GWAS Catalog, a curated resource of SNP-trait associations. Nucleic Acids Research, 42, D1001–D1006.

7. Browning, B.L., Zhou, Y. and Browning, S.R. (2018) A One-Penny Imputed Genome from Next-Generation Reference Panels. Am J Hum Genet, 103, 338–348.

8. Marchini, J., Howie, B., Myers, S., McVean, G. and Donnelly, P. (2007) A new multipoint method for genome-wide association studies by imputation of genotypes. Nat Genet, 39, 906–913.

9. Das, S., Forer, L., Schonherr, S., Sidore, C., Locke, A.E., Kwong, A., Vrieze, S.I., Chew, E.Y., Levy, S., McGue, M. et al. (2016) Next-generation genotype imputation service and methods. Nat Genet, 48, 1284–1287.

10. Rubinacci, S., Ribeiro, D.M., Hofmeister, R.J. and Delaneau, O. (2021) Efficient phasing and imputation of low-coverage sequencing data using large reference panels (vol 53, pg 120, 2021). Nat Genet, 53, 412–412.

11. Das, S., Abecasis, G.R. and Browning, B.L. (2018) Genotype Imputation from Large Reference Panels. Annu Rev Genom Hum G, 19, 73–96.

12. Altshuler, D.M., Durbin, R.M., Abecasis, G.R., Bentley, D.R., Chakravarti, A., Clark, A.G., Donnelly, P., Eichler, E.E., Flicek, P., Gabriel, S.B. et al. (2015) A global reference for human genetic variation. Nature, 526, 68-+.

13. McCarthy, S., Das, S., Kretzschmar, W., Delaneau, O., Wood, A.R., Teumer, A., Kang, H.M., Fuchsberger, C., Danecek, P., Sharp, K. et al. (2016) A reference panel of 64,976 haplotypes for genotype imputation. Nat Genet, 48, 1279–1283.

14. Yan, G., Qiao, R., Zhang, F., Xin, W., Xiao, S., Huang, T., Zhang, Z. and Huang, L. (2017) Imputation-Based Whole-Genome Sequence Association Study Rediscovered the Missing QTL for Lumbar Number in Sutai Pigs. Sci Rep, 7, 615.

15. Song, H., Ye, S., Jiang, Y., Zhang, Z., Zhang, Q. and Ding, X. (2019) Using imputation-based whole-genome sequencing data to improve the accuracy of genomic prediction for combined populations in pigs. Genet Sel Evol, 51, 58.

16. Huang, J., Howie, B., McCarthy, S., Memari, Y., Walter, K., Min, J.L., Danecek, P., Malerba, G., Trabetti, E., Zheng, H.F. et al. (2015) Improved imputation of low-frequency and rare variants using the UK10K haplotype reference panel. Nat Commun, 6.

17. Delaneau, O., Zagury, J.F. and Marchini, J. (2013) Improved whole-chromosome phasing for disease and population genetic studies. Nat Methods, 10, 5–6.

18. Yang, W., Yang, Y., Zhao, C., Yang, K., Wang, D., Yang, J., Niu, X. and Gong, J. (2020) Animal-ImputeDB: a comprehensive database with multiple animal reference panels for genotype imputation. Nucleic Acids Res, 48, D659–D667.

19. Xu, P., Ni, L., Tao, Y., Ma, Z., Hu, T., Zhao, X., Yu, Z., Lu, C., Zhao, X. and Ren, J. (2020) Genome-wide association study for growth and fatness traits in Chinese Sujiang pigs. Anim Genet, 51, 314–318.

20. Zhang, Z., Chen, Z., Ye, S., He, Y., Huang, S., Yuan, X., Chen, Z., Zhang, H. and Li, J. (2019) Genome-Wide Association Study for Reproductive Traits in a Duroc Pig Population. Animals (Basel), 9.

21. Zhang, Z., Chen, Z.-t., Diao, S.-q., Ye, S.-p., Wang, J.-y., Gao, N., Yuan, X.-l., Chen, Z.-m., Zhang, H. and Li, J.-q. (2020) Identifying the complex genetic architecture of growth and fatness traits in a Duroc pig population. Journal of Integrative Agriculture, 19.

22. Chen, S., Zhou, Y., Chen, Y. and Gu, J. (2018) fastp: an ultra-fast all-in-one FASTQ preprocessor. Bioinformatics, 34, i884–i890.

23. Li, H. and Durbin, R. (2009) Fast and accurate short read alignment with Burrows-Wheeler transform. Bioinformatics, 25, 1754–1760.

24. Li, H., Handsaker, B., Wysoker, A., Fennell, T., Ruan, J., Homer, N., Marth, G., Abecasis, G., Durbin, R. and Genome Project Data Processing, S. (2009) The Sequence Alignment/Map format and SAMtools. Bioinformatics, 25, 2078–2079.

25. Tarasov, A., Vilella, A.J., Cuppen, E., Nijman, I.J. and Prins, P. (2015) Sambamba: fast processing of NGS alignment formats. Bioinformatics, 31, 2032–2034.

26. Pedersen, B.S. and Quinlan, A.R. (2018) Mosdepth: quick coverage calculation for genomes and exomes. Bioinformatics, 34, 867–868.

27. Van der Auwera, G.A., Carneiro, M.O., Hartl, C., Poplin, R., Del Angel, G., Levy-Moonshine, A., Jordan, T., Shakir, K., Roazen, D., Thibault, J. et al. (2013) From FastQ data to high confidence variant calls: the Genome Analysis Toolkit best practices pipeline. Curr Protoc Bioinformatics, 43, 11 10 11–11 10 33.

28. Jiang, Y.F., Jiang, Y., Wang, S., Zhang, Q. and Ding, X.D. (2019) Optimal sequencing depth design for whole genome re-sequencing in pigs. Bmc Bioinformatics, 20.

29. Chang, C.C., Chow, C.C., Tellier, L.C., Vattikuti, S., Purcell, S.M. and Lee, J.J. (2015) Second-generation PLINK: rising to the challenge of larger and richer datasets. Gigascience, 4, 7.

30. Purcell, S., Neale, B., Todd-Brown, K., Thomas, L., Ferreira, M.A., Bender, D., Maller, J., Sklar, P., de Bakker, P.I., Daly, M.J. et al. (2007) PLINK: a tool set for whole-genome association and population-based linkage analyses. Am J Hum Genet, 81, 559–575.

31. Delaneau, O., Marchini, J. and Zagury, J.F. (2012) A linear complexity phasing method for thousands of genomes. Nat Methods, 9, 179–181.

32. Zhou, X. and Stephens, M. (2012) Genome-wide efficient mixed-model analysis for association studies. Nat Genet, 44, 821–824.

33. Yang, J., Lee, S.H., Goddard, M.E. and Visscher, P.M. (2011) GCTA: a tool for genome-wide complex trait analysis. Am J Hum Genet, 88, 76–82.

34. Perez, P. and de los Campos, G. (2014) Genome-wide regression and prediction with the BGLR statistical package. Genetics, 198, 483–495.

35. Danecek, P., Auton, A., Abecasis, G., Albers, C.A., Banks, E., DePristo, M.A., Handsaker, R.E., Lunter, G., Marth, G.T., Sherry, S.T. et al. (2011) The variant call format and VCFtools. Bioinformatics, 27, 2156–2158.

36. Fan, B., Onteru, S.K., Plastow, G.S. and Rothschild, M.F. (2009) Detailed characterization of the porcine MC4R gene in relation to fatness and growth. Anim Genet, 40, 401–409.

37. Sasaki, Y., Ohta, M., Desai, D., Figueiredo, J.L., Whelan, M.C., Sugano, T., Yamabi, M., Yano, W., Faits, T., Yabusaki, K. et al. (2015) Angiopoietin Like Protein 2 (ANGPTL2) Promotes Adipose Tissue Macrophage and T lymphocyte Accumulation and Leads to Insulin Resistance. Plos One, 10, e0131176.

38. Frischknecht, M., Meuwissen, T.H.E., Bapst, B., Seefried, F.R., Flury, C., Garrick, D., Signer-Hasler, H., Stricker, C., Bieber, A., Fries, R. et al. (2018) Genomic prediction using imputed whole-genome sequence variants in Brown Swiss Cattle. J Dairy Sci, 101, 1292–1296.

39. Ober, U., Ayroles, J.F., Stone, E.A., Richards, S., Zhu, D.H., Gibbs, R.A., Stricker, C., Gianola, D., Schlather, M., Mackay, T.F.C. et al. (2012) Using Whole-Genome Sequence Data to Predict Quantitative Trait Phenotypes in Drosophila melanogaster. Plos Genet, 8.

40. van Binsbergen, R., Calus, M.P.L., Bink, M.C.A.M., van Eeuwijk, F.A., Schrooten, C. and Veerkamp, R.F. (2015) Genomic prediction using imputed whole-genome sequence data in Holstein Friesian cattle. Genetics Selection Evolution, 47.

41. Mathew, B., Leon, J. and Sillanpaa, M.J. (2018) A novel linkage-disequilibrium corrected genomic relationship matrix for SNP-heritability estimation and genomic prediction. Heredity (Edinb), 120, 356–368.

42. Edwards, S.M., Sorensen, I.F., Sarup, P., Mackay, T.F.C. and Sorensen, P. (2016) Genomic Prediction for Quantitative Traits Is Improved by Mapping Variants to Gene Ontology Categories in Drosophila melanogaster. Genetics, 203, 1871-+.

43. Zhang, Z., Liu, J.F., Ding, X.D., Bijma, P., de Koning, D.J. and Zhang, Q. (2010) Best Linear Unbiased Prediction of Genomic Breeding Values Using a Trait-Specific Marker-Derived Relationship Matrix. Plos One, 5.

44. Al Kalaldeh, M., Gibson, J., Duijvesteijn, N., Daetwyler, H.D., MacLeod, I., Moghaddar, N., Lee, S.H. and van der Werf, J.H.J. (2019) Using imputed wholegenome sequence data to improve the accuracy of genomic prediction for parasite resistance in Australian sheep. Genetics Selection Evolution, 51.

45. Song, H.L., Ye, S.P., Jiang, Y.F., Zhang, Z., Zhang, Q. and Ding, X.D. (2019) Using imputation-based whole-genome sequencing data to improve the accuracy of genomic prediction for combined populations in pigs. Genetics Selection Evolution, 51.

