## Supplementary material for "PHARP: A pig haplotype reference panel for genotype imputation": PHARP-Supplementary-Tables and Figures

**Table S1. WGS data source for pig haplotypes reference panel construction and imputation accuracy assessment.**

| Type | Accession | Pig breeds | Sample size | Ref |
|---|---|---|---|---|
| WGS data for PHARP construction | - | AQ, DU, DY, HBBZ, JH, JS, LW, XDH | 84 | Lab* |
| | PRJEB1683 | EWB, AWB, DU, HA, PI, LR, LW, XZ, JQH, MS | 72 | [1-4] |
| | PRJNA144099 | WZS | 1 | [5] |
| | PRJNA41185 | DU | 1 | - |
| | PRJNA176189 | GM | 1 | [6] |
| | PRJNA231897 | RC | 6 | - |
| | PRJNA186497 | AWB, TWB, PZ, WJ, YN, NJ, JH | 49 | [7] |
| | PRJNA238851 | TC | 4 | [8] |
| | PRJNA260763 | DU, LR, YM, KWB, LW | 70 | [9, 10] |
| | PRJEB9115 | DU | 1 | - |
| | PRJNA213179 | AWB, EHL, HT, LWU, LC, MI, MX, TP, WZS | 69 | [11] |
| | PRJNA221763 | BK | 3 | [12] |
| | PRJNA239399 | MA, DU | 4 | [13] |
| | PRJNA190683 | IB | 1 | - |
| | PRJNA255085 | EWB, GC, IB, MP, LW | 6 | [14] |
| | PRJEB9326 | PI | 10 | [15] |
| | PRJEB9922 | EWB, AWB, WS, TP, LS, AS, BB, BK, BS, CM, CS, CA, CT, GO, HA, LB, LI, MA, MW, IB, NS, TA, RE, JQH, XZ | 91 | [16] |
| | PRJNA281548 | BK | 10 | [17] |
| | PRJNA254936 | KP | 14 | [18] |
| | PRJNA273907 | LW, LR | 2 | - |
| | PRJNA438040 | WZS | 1 | - |
| | PRJNA506339 | LW | 16 | - |
| | PRJNA524263 | WB | 20 | - |
| | PRJNA369600 | IB, LS | 7 | [19] |
| | PRJNA418771 | NS | 1 | - |
| | PRJNA493166 | LW | 3 | [20] |
| | PRJNA487172 | DU | 4 | [21] |
| | PRJEB27654 | GM | 12 | [22] |
| | PRJNA343658 | LS, DU, LW | 72 | [23] |
| | PRJNA305975 | DSE | 31 | [24] |

| | | | | |
|---|---|---|---|---|
| | PRJNA322309 | RC, DU | 8 | - |
| | PRJNA320525 | WB, IB, ICF, YM, TA | 9 | [25, 26] |
| | PRJNA354435 | DP | 1 | [27] |
| | PRJNA378496 | MS, DU, AWB, DW, DSE, TP, LW, RC, MZ | 71 | [28] |
| | PRJEB29465 | DU, PL | 5 | |
| | PRJNA507853 | DLY, LL, LW, DU, LR | 10 | [29] |
| | PRJNA305081 | MZ, AWB | 11 | - |
| | PRJNA291011 | GM | 1 | [30] |
| | PRJNA482384 | USMARC | 29 | [31] |
| | PRJNA414091 | USMARC | 108 | [31] |
| | PRJNA309108 | BaM, BK, HA, JH, LR, LW, MS, PI, RC | 9 | [32] |
| | PRJNA553106 | PLL, PI, MS, LW, LL | 115 | [33] |
| | PRJEB35180 | Ssc | 5 | - |
| | PRJNA488327 | EHL, DU | 19 | [34] |
| | PRJNA398176 | JH, NJ, BaM, BS | 24 | [35] |
| | PRJNA550237 | DU, EHL, LW, TP | 63 | [36] |
| | PRJEB37956 | LW | 5 | [37] |
| | PRJNA622908 | LR | 5 | [37] |
| | PRJNA626370 | DU | 5 | - |
| | PRJNA488960 | TC, LW | 12 | - |
| **Total** | | | **1,181** | |
| Test dataset 1 | PRJEB39374 | LW | 76 | [37] |
| | PRJEB38156 | LW | 5 | [37] |
| **Total** | | | **81** | |
| Test dataset 2 | - | JXH | 54 | Lab* |
| Test dataset 3 | - | DU | 299 | Lab* |
| Test dataset 4 | - | DU | 20 | Lab* |

Note: * WGS data generated by our laboratory.

AQ: Anqingliubai, AS: Angler Sattleschwein, AWB: Asian wild boar, BaM: Bamei, BB: Bunte Bentheimer, BK: Berkshire, BS: Baoshan, BS: British Saddleback, CA: Calabrese, CM: Chato Murciano, CS: Cinta Senese, CT: Casertana, DLY: Duroc x (Landrace x Yorkshire), DP: Duroc x Pietrain, DSE: Diannan small ear, DU: Duroc, DW: Daweizi, DY: Dingyuan, EHL: Erhualian, EWB: European wild boar, GC: Guatemala Creole pig, GM: GÌ_ttingen minipig, GO: Gloucester Old Spot, HA: Hampshire, HBBZ: Huibeibai, HT: Hetao, IB: Iberian, ICF: Isla del Coco feral pigs (Costa Rica), JH: Jinhua, JQH: Jiangquhai, JS: Jishen, JXH: Jiaxinghei, KP: Korean pig, KWB: Korean wild boar, LB: Large Black, LC: Luchuan, LI: Linderodsvin, LL: Landrace x Large White, LR: Landrace, LS: Leping Spotted, LW: Large White, LWU: Laiwu, MA: Mangalica, MI:

Mizhu, MP: 16th century pig, MS: Meishan, MW: Middle White, MX: Bamaxiang, MZ: Min, NJ: Neijiang, NS: Nera Siciliana, PI: Pietrain, PL: Pietrain x Swiss Landrace , PLL: Pietrain x (Landrace x Large White), PZ: Penzhou, RC: Rongchang, RE: Retinto, Ssc: Sus scrofa, TA: Tamworth, TC: Tongcheng, TP: Tibetan pig, TWB: Tibetan wild boar, USMARC: USDA-ARS-USMARC, WB: Wild boar, WJ: Wujin, WS: Wannan Spotted, WZS: Wuzhishan, XDH: Xiaduhei, XZ: Xiang, YM: Yucatan miniature pig, YN: Yanan,

**Table S2. GWAS datasets and candidate genes reported in the original paper and identified using imputed-genotypes.**

| Traits | Samples | Phenotypes | Associations (candidate genes) | |
|---|---|---|---|---|
| | | | [1]Original report | [2]Imputed-genotypes |
| [3]Growth and fatness traits | N=1,067, Duroc pig breed, 60K & 80K, [38] | D100 | *VPS4B*<br>*PHLPP1*<br>*CDH20*<br>*ENSSSCG00000034988*<br>*ENSSSCG00000004911* | *PHLPP1*<br>*ENSSSCG00000034988*<br>*ENSSSCG00000004911*<br>***TNFRSF11A***<br>***MC4R***<br>***CCBE1***<br>***LMAN1***<br>***CPLX4***<br>***ENSSSCG00000048150***<br>***ENSSSCG00000039179***<br>***ENSSSCG00000041383***<br>***ENSSSCG00000048538***<br>***ENSSSCG00000050691***<br>***ENSSSCG00000045579***<br>***ENSSSCG00000047436*** |
| | | L100 | *ADAMTS2* | ***MC4R***<br>***TSHZ1***<br>***DXO***<br>***STK19***<br>***PRRT1***<br>***TINAG***<br>***MLIP***<br>***GRM4***<br>***NUDT3***<br>***HMGA1***<br>***PACSIN1***<br>***SPDEF***<br>***ILRUN***<br>***SNRPC***<br>***UHRF1BP1***<br>***TAF11***<br>***ANKS1A***<br>***ENSSSCG00000037476***<br>***ENSSSCG00000023160***<br>***ENSSSCG00000048538***<br>***ENSSSCG00000050691***<br>***ENSSSCG00000045579*** |
| | | B100 | *SNRPC*<br>*GRM4*<br>*TXN2*<br>*TSHZ1*<br>*PHLLP1*<br>*NUDT3*<br>*ENSSSCG00000034988*<br>*ENSSSCG00000004911* | *ENSSSCG00000051188*<br>***MAML1***<br>***CANX***<br>***HNRNPH1***<br>***RUFY1***<br>***ADAMTS2***<br>***ZNF879***<br>***GRM6***<br>***ZNF454***<br>***ZFP2***<br>***ZNF354B***<br>***PROP1***<br>***CLK4***<br>***COL23A1***<br>***RNF44***<br>***CLTB***<br>***ssc-mir-1271***<br>***ARL10***<br>***KIAA1191***<br>***SIMC1***<br>***THOC3***<br>***SFXN1*** |

| | | | | |
|---|---|---|---|---|
| | | | | ***TMEM171***<br>***ENSSSCG00000046532***<br>***ENSSSCG00000044467***<br>***ENSSSCG00000041652***<br>***ENSSSCG00000034785***<br>***ENSSSCG00000042891***<br>***ENSSSCG00000042242*** |
| [3]Reproduction traits | N=1,067, Duroc pig breed, 60K & 80K, [39] | LSB | *TNX*<br>*KCNA1*<br>*ZDHHC18*<br>*MAP2K6*<br>*BICC1* | *BICC1*<br>***NCF4***<br>***IFT27***<br>***CACNG2***<br>***ENSSSCG00000000145***<br>***ENSSSCG00000041362***<br>***ENSSSCG00000036761*** |
| | | LWB | *FAM135B*<br>*EPHB2*<br>*SEMA4D* | ***PEBP4***<br>***PLEK***<br>***CNRIP1***<br>***PNO1***<br>***WDR92***<br>***ZC3H3***<br>***PTK2***<br>***SPATC1***<br>***ENSSSCG00000036115***<br>***ENSSSCG00000034561***<br>***ENSSSCG00000039331***<br>***ENSSSCG00000041622*** |
| | | LSW | *TMEM132D*<br>*TBX3*<br>*FAM110A* | - |
| [4]Growth and fatness traits | N=365, Sujiang pig breed, [40] | BW | *GABRB3*<br>*ZNF106* | - |
| | | BL | - | - |
| | | BH | - | ***ENSSSCG00000032052*** |
| | | CC | - | - |
| | | CW | - | - |
| | | HW | - | - |
| | | BF | *XKR4*<br>*MGAM*<br>*TAS2R38* | ***RABEPK***<br>***LMX1B***<br>***RALGPS1***<br>***ANGPTL2***<br>***LRSAM1***<br>***NIBAN2***<br>***SH2D3C***<br>***ST6GALNAC4***<br>***PIP5KL1***<br>***DPM2***<br>***FAM102A***<br>***CCL8***<br>***ASIC2***<br>***TNXB***<br>***ENSSSCG00000022295*** |

Notes:

[1]Candidate genes reported in the original paper.

[2]Candidate genes identified using imputed-genotypes and novel genes were bolded.

[3]The original study used 32,147 SNPs for GWAS and took the genome-wide suggestive significant threshold as to be $1.08 \times 10^{-4}$; imputed-genotypes after using the same filtrations as in the ordinal study included 10,856,918 SNPs and took the genome-wide suggestive significant threshold as to be $2.05 \times 10^{-6}$ (1/ 488,357).

[4]The original study used 53,702 SNPs and took the genome-wide significant threshold as to be $9.31 \times 10^{-7}$; the imputed-genotype after using the same filtration as in the ordinal study included 19,401,022 SNPs and took the genome-wide suggestive significant threshold as to be $6.46 \times 10^{-7}$ (1/1,548,065).

D100 = days to 100 kg; L100 = loin muscle area at 100 kg; B100 = backfat thickness at 100 kg; LSB = litter size at birth; LWB= litter weight at birth; LSW = litter size at weaning; BW = body weight; BL = body length; BH = body height; CC = chest circumference; CW = chest width, HW = hip width; BF = backfat thickness.

**Table S3. Imputation accuracy evaluated from test datasets.**

| Test datasets | [1]Mimic chips | No. of imputed genotypes | No. of consistent genotypes | [2]Inconsistent genotypes (percentage) | | | | CR | $r^2$ |
|---|---|---|---|---|---|---|---|---|---|
| | | | | No. | Ho_Ho | He_Ho | Ho_He | | |
| Test dataset 1 (LW, n=81) | 50K | 2,628,610,057 | 75,145,634 | 2,553,464,423 | 0.057 | 0.488 | 0.456 | 0.971 | 0.920 |
| | 60K | 2,628,136,817 | 69,129,000 | 2,559,007,817 | 0.055 | 0.489 | 0.455 | 0.974 | 0.926 |
| | 80K | 2,627,590,208 | 58,066,262 | 2,569,523,946 | 0.050 | 0.480 | 0.469 | 0.978 | 0.939 |
| Test dataset 2 (DU, n=299) | 50K | 2,327,943,137 | 153,430,902 | 2,174,512,235 | 0.041 | 0.503 | 0.456 | 0.934 | 0.876 |
| | 60K | 2,327,099,658 | 141,230,486 | 2,185,869,172 | 0.039 | 0.495 | 0.467 | 0.939 | 0.886 |
| | 80K | 2,325,327,186 | 124,017,370 | 2,201,309,816 | 0.036 | 0.492 | 0.472 | 0.947 | 0.901 |
| Test dataset 3 (JXH, n=54) | 50K | 1,093,333,104 | 208,940,405 | 884,392,699 | 0.257 | 0.308 | 0.435 | 0.809 | 0.488 |
| | 60K | 1,093,163,149 | 201,269,754 | 891,893,395 | 0.248 | 0.305 | 0.447 | 0.816 | 0.510 |
| | 80K | 1,093,001,361 | 199,369,412 | 893,631,949 | 0.245 | 0.305 | 0.451 | 0.818 | 0.517 |
| [3]Test dataset 4 (DU, n = 20) | 50K | 160,966,260 | 149,064,448 | 11,901,812 | 0.040 | 0.537 | 0.423 | 0.926 | 0.869 |

Notes:

[1]Mimicing the most popular commercial porcine SNP chips (50K, 60K and 80K) by extracting SNPs for test datasets at all sites included in these commercial porcine microarray genotyping platforms.

[2]Inconsistent genotypes types: Ho_Ho, a homozygote was imputed to be another homozygote; He_Ho, a heterozygote was imputed to be a homozygote; Ho_He, a homozygote was imputed to be a heterozygote.

[3]Duroc pigs (n = 20) genotyped by 50K were used as input for imputation and the imputed-genotypes were compared with the same individuals genotyped by ELC (regarded as real).

**Table S4. Imputation accuracy estimated by mimicking the target panel with different densities (repeated 5 times) of SNPs on chromosomes using test datasets.**

| Test datasets | Chr. | Density Kb per SNP | No. of imputed genotypes | No. of consistent genotypes | Inconsistent genotypes (percentage) | | | | CR | $r^2$ |
|---|---|---|---|---|---|---|---|---|---|---|
| | | | | | No. | Ho_Ho | He_Ho | Ho_He | | |
| Test dataset 1 | chr1 | 2.5 | 92.32±0 | 91.1±0.01 | 1.22±0.01 | 0.021±0 | 0.384±0.002 | 0.595±0.002 | 0.987±0 | 0.976±0 |
| | | 5 | 96.02±0 | 94.58±0.01 | 1.44±0.01 | 0.023±0 | 0.412±0.003 | 0.565±0.003 | 0.985±0 | 0.973±0 |
| | | 10 | 98.05±0 | 96.22±0.01 | 1.83±0.01 | 0.029±0.001 | 0.44±0.004 | 0.531±0.003 | 0.981±0 | 0.966±0 |
| | | 20 | 99.12±0 | 96.55±0.04 | 2.57±0.04 | 0.036±0.002 | 0.471±0.003 | 0.493±0.002 | 0.974±0 | 0.952±0.001 |
| | | 40 | 99.67±0 | 95.7±0.05 | 3.97±0.05 | 0.046±0.002 | 0.495±0.004 | 0.459±0.004 | 0.96±0 | 0.925±0.001 |
| | | 50 | 99.78±0 | 95.15±0.12 | 4.63±0.12 | 0.046±0.001 | 0.496±0.001 | 0.458±0.001 | 0.954±0.001 | 0.913±0.002 |
| | | 60 | 99.85±0 | 94.55±0.1 | 5.3±0.1 | 0.052±0.001 | 0.506±0.007 | 0.442±0.007 | 0.947±0.001 | 0.899±0.002 |
| | | 80 | 99.95±0 | 93.64±0.1 | 6.31±0.1 | 0.056±0.003 | 0.503±0.005 | 0.44±0.006 | 0.937±0.001 | 0.88±0.002 |
| | | 100 | 100±0 | 92.58±0.14 | 7.42±0.14 | 0.061±0.002 | 0.511±0.003 | 0.427±0.005 | 0.926±0.001 | 0.858±0.003 |
| | | 200 | 100.11±0 | 88.17±0.23 | 11.94±0.23 | 0.079±0.001 | 0.525±0.002 | 0.396±0.001 | 0.881±0.002 | 0.768±0.004 |
| | | 400 | 100.17±0 | 82.64±0.28 | 17.53±0.28 | 0.105±0.004 | 0.547±0.001 | 0.348±0.004 | 0.825±0.003 | 0.654±0.005 |
| | chr2 | 2.5 | 61.53±0 | 60.56±0.02 | 0.97±0.02 | 0.022±0 | 0.439±0.008 | 0.539±0.008 | 0.984±0 | 0.972±0 |
| | | 5 | 63.43±0 | 62.26±0.01 | 1.16±0.01 | 0.025±0.001 | 0.457±0.003 | 0.518±0.003 | 0.982±0 | 0.967±0 |
| | | 10 | 64.46±0 | 62.92±0.01 | 1.53±0.01 | 0.03±0.001 | 0.477±0.006 | 0.493±0.005 | 0.976±0 | 0.957±0 |
| | | 20 | 65±0 | 62.71±0.04 | 2.29±0.04 | 0.039±0.001 | 0.495±0.005 | 0.466±0.004 | 0.965±0.001 | 0.935±0.001 |
| | | 40 | 65.28±0 | 61.59±0.04 | 3.69±0.04 | 0.049±0.002 | 0.511±0.005 | 0.44±0.005 | 0.943±0.001 | 0.894±0.001 |
| | | 50 | 65.34±0 | 60.87±0.11 | 4.47±0.11 | 0.056±0.002 | 0.512±0.005 | 0.432±0.006 | 0.932±0.002 | 0.87±0.004 |
| | | 60 | 65.37±0 | 60.3±0.07 | 5.08±0.07 | 0.058±0.002 | 0.516±0.003 | 0.426±0.002 | 0.922±0.001 | 0.853±0.002 |
| | | 80 | 65.42±0 | 59.15±0.08 | 6.28±0.08 | 0.066±0.002 | 0.514±0.005 | 0.421±0.006 | 0.904±0.001 | 0.816±0.003 |
| | | 100 | 65.45±0 | 58.24±0.15 | 7.21±0.15 | 0.067±0.004 | 0.517±0.002 | 0.416±0.004 | 0.89±0.002 | 0.79±0.004 |
| | | 200 | 65.51±0 | 53.79±0.12 | 11.71±0.12 | 0.092±0.003 | 0.528±0.004 | 0.38±0.006 | 0.821±0.002 | 0.655±0.002 |
| | | 400 | 65.54±0 | 48.96±0.27 | 16.58±0.27 | 0.122±0.005 | 0.542±0.005 | 0.336±0.004 | 0.747±0.004 | 0.509±0.01 |
| | chr3 | 2.5 | 60.08±0 | 59.25±0 | 0.83±0 | 0.022±0.001 | 0.371±0.002 | 0.607±0.002 | 0.986±0 | 0.974±0 |
| | | 5 | 62.02±0 | 60.99±0.01 | 1.03±0.01 | 0.026±0.001 | 0.405±0.003 | 0.568±0.002 | 0.983±0 | 0.969±0 |
| | | 10 | 63.04±0 | 61.63±0.02 | 1.4±0.02 | 0.034±0.002 | 0.441±0.003 | 0.525±0.004 | 0.978±0 | 0.957±0.001 |
| | | 20 | 63.57±0 | 61.41±0.03 | 2.16±0.03 | 0.041±0.002 | 0.48±0.005 | 0.479±0.005 | 0.966±0 | 0.934±0.001 |
| | | 40 | 63.83±0 | 60.3±0.04 | 3.54±0.04 | 0.053±0.003 | 0.503±0.007 | 0.444±0.005 | 0.945±0.001 | 0.89±0.002 |
| | | 50 | 63.89±0 | 59.71±0.05 | 4.17±0.05 | 0.056±0.002 | 0.509±0.004 | 0.435±0.005 | 0.935±0.001 | 0.87±0.002 |
| | | 60 | 63.92±0 | 59.15±0.06 | 4.77±0.06 | 0.059±0.002 | 0.514±0.004 | 0.427±0.004 | 0.925±0.001 | 0.851±0.002 |

| | | | | | | | | | | |
|---|---|---|---|---|---|---|---|---|---|---|
| | | 80 | 63.97±0 | 58.2±0.1 | 5.76±0.1 | 0.065±0.001 | 0.524±0.005 | 0.411±0.004 | 0.91±0.002 | 0.82±0.003 |
| | | 100 | 63.99±0 | 57.11±0.12 | 6.89±0.12 | 0.07±0.004 | 0.523±0.008 | 0.407±0.006 | 0.892±0.002 | 0.784±0.005 |
| | | 200 | 64.05±0 | 53.68±0.19 | 10.37±0.19 | 0.088±0.004 | 0.535±0.002 | 0.377±0.004 | 0.838±0.003 | 0.673±0.008 |
| | | 400 | 64.07±0 | 49.73±0.28 | 14.35±0.28 | 0.109±0.005 | 0.555±0.005 | 0.336±0.006 | 0.776±0.004 | 0.549±0.009 |
| | chr4 | 2.5 | 58.82±0 | 58.04±0 | 0.79±0 | 0.029±0.001 | 0.353±0.002 | 0.617±0.002 | 0.987±0 | 0.975±0 |
| | | 5 | 60.67±0 | 59.67±0.01 | 1±0.01 | 0.034±0.001 | 0.391±0.004 | 0.575±0.003 | 0.984±0 | 0.969±0 |
| | | 10 | 61.66±0 | 60.21±0.02 | 1.44±0.02 | 0.042±0.002 | 0.431±0.004 | 0.526±0.005 | 0.977±0 | 0.955±0.001 |
| | | 20 | 62.17±0 | 59.92±0.1 | 2.25±0.1 | 0.053±0.001 | 0.462±0.003 | 0.484±0.003 | 0.964±0.002 | 0.928±0.003 |
| | | 40 | 62.43±0 | 58.74±0.08 | 3.7±0.08 | 0.061±0.003 | 0.486±0.003 | 0.453±0.004 | 0.941±0.001 | 0.882±0.003 |
| | | 50 | 62.49±0 | 58.16±0.12 | 4.33±0.12 | 0.065±0.002 | 0.485±0.004 | 0.451±0.004 | 0.931±0.002 | 0.861±0.004 |
| | | 60 | 62.52±0 | 57.63±0.14 | 4.89±0.14 | 0.067±0.004 | 0.496±0.005 | 0.437±0.006 | 0.922±0.002 | 0.843±0.005 |
| | | 80 | 62.57±0 | 56.71±0.13 | 5.86±0.13 | 0.072±0.002 | 0.5±0.003 | 0.428±0.003 | 0.906±0.002 | 0.812±0.004 |
| | | 100 | 62.59±0 | 55.75±0.11 | 6.84±0.11 | 0.077±0.003 | 0.508±0.003 | 0.415±0.003 | 0.891±0.002 | 0.78±0.005 |
| | | 200 | 62.64±0 | 51.99±0.21 | 10.66±0.21 | 0.094±0.001 | 0.525±0.005 | 0.381±0.005 | 0.83±0.003 | 0.658±0.007 |
| | | 400 | 62.67±0 | 48.05±0.22 | 14.62±0.22 | 0.117±0.003 | 0.547±0.006 | 0.336±0.008 | 0.767±0.003 | 0.533±0.007 |
| | chr5 | 2.5 | 51.14±0 | 50.51±0 | 0.63±0 | 0.023±0.001 | 0.349±0.003 | 0.628±0.002 | 0.988±0 | 0.977±0 |
| | | 5 | 52.69±0 | 51.92±0.01 | 0.77±0.01 | 0.026±0.001 | 0.382±0.004 | 0.592±0.004 | 0.985±0 | 0.973±0 |
| | | 10 | 53.5±0 | 52.45±0.02 | 1.05±0.02 | 0.03±0.001 | 0.427±0.003 | 0.543±0.003 | 0.98±0 | 0.963±0.001 |
| | | 20 | 53.92±0 | 52.25±0.03 | 1.67±0.03 | 0.037±0.002 | 0.47±0.003 | 0.493±0.004 | 0.969±0.001 | 0.941±0.001 |
| | | 40 | 54.12±0 | 51.17±0.03 | 2.95±0.03 | 0.046±0.002 | 0.493±0.003 | 0.461±0.004 | 0.945±0.001 | 0.894±0.001 |
| | | 50 | 54.17±0 | 50.69±0.12 | 3.48±0.12 | 0.05±0.004 | 0.502±0.005 | 0.448±0.005 | 0.936±0.002 | 0.875±0.006 |
| | | 60 | 54.19±0 | 50.11±0.1 | 4.08±0.1 | 0.057±0.003 | 0.505±0.007 | 0.438±0.007 | 0.925±0.002 | 0.852±0.004 |
| | | 80 | 54.23±0 | 49.14±0.09 | 5.09±0.09 | 0.062±0.002 | 0.51±0.004 | 0.429±0.004 | 0.906±0.002 | 0.815±0.004 |
| | | 100 | 54.25±0 | 48.14±0.12 | 6.11±0.12 | 0.069±0.003 | 0.512±0.006 | 0.418±0.005 | 0.887±0.002 | 0.776±0.005 |
| | | 200 | 54.29±0 | 44.83±0.28 | 9.46±0.28 | 0.087±0.003 | 0.529±0.002 | 0.384±0.003 | 0.826±0.005 | 0.653±0.011 |
| | | 400 | 54.31±0 | 41.06±0.31 | 13.26±0.31 | 0.108±0.004 | 0.552±0.009 | 0.34±0.007 | 0.756±0.006 | 0.518±0.012 |
| | chr6 | 2.5 | 71.2±0 | 70.13±0.01 | 1.07±0.01 | 0.025±0.001 | 0.411±0.003 | 0.564±0.003 | 0.985±0 | 0.972±0 |
| | | 5 | 73.56±0 | 72.27±0.01 | 1.29±0.01 | 0.03±0.001 | 0.434±0.002 | 0.536±0.002 | 0.982±0 | 0.967±0 |
| | | 10 | 74.83±0 | 73.15±0.02 | 1.67±0.02 | 0.035±0.002 | 0.46±0.005 | 0.505±0.006 | 0.978±0 | 0.958±0.001 |
| | | 20 | 75.49±0 | 73.07±0.03 | 2.42±0.03 | 0.043±0.001 | 0.482±0.004 | 0.475±0.003 | 0.968±0 | 0.938±0.001 |
| | | 40 | 75.82±0 | 72.13±0.03 | 3.69±0.03 | 0.049±0.001 | 0.5±0.005 | 0.451±0.005 | 0.951±0 | 0.906±0.001 |
| | | 50 | 75.89±0 | 71.48±0.07 | 4.42±0.07 | 0.054±0.001 | 0.507±0.005 | 0.438±0.006 | 0.942±0.001 | 0.887±0.002 |
| | | 60 | 75.94±0 | 70.86±0.15 | 5.08±0.15 | 0.056±0.002 | 0.508±0.003 | 0.436±0.003 | 0.933±0.002 | 0.87±0.004 |
| | | 80 | 76±0 | 69.66±0.08 | 6.34±0.08 | 0.065±0.002 | 0.511±0.002 | 0.424±0.002 | 0.917±0.001 | 0.836±0.002 |
| | | 100 | 76.03±0 | 68.64±0.09 | 7.39±0.09 | 0.07±0.003 | 0.514±0.003 | 0.416±0.003 | 0.903±0.001 | 0.808±0.003 |

| | | | | | | | | | | |
|---|---|---|---|---|---|---|---|---|---|---|
| | | 200 | 76.1±0 | 63.97±0.21 | 12.13±0.21 | 0.087±0.002 | 0.526±0.005 | 0.387±0.003 | 0.841±0.003 | 0.684±0.005 |
| | | 400 | 76.13±0 | 58.49±0.29 | 17.64±0.29 | 0.109±0.002 | 0.54±0.003 | 0.351±0.005 | 0.768±0.004 | 0.543±0.007 |
| | chr7 | 2.5 | 60.41±0 | 59.23±0.01 | 1.18±0.01 | 0.025±0.001 | 0.494±0.005 | 0.481±0.005 | 0.98±0 | 0.964±0 |
| | | 5 | 62.21±0 | 60.8±0.02 | 1.41±0.02 | 0.029±0.001 | 0.51±0.004 | 0.461±0.004 | 0.977±0 | 0.957±0.001 |
| | | 10 | 63.16±0 | 61.36±0.03 | 1.8±0.03 | 0.035±0.001 | 0.522±0.005 | 0.443±0.006 | 0.972±0 | 0.946±0.001 |
| | | 20 | 63.64±0 | 61.01±0.05 | 2.63±0.05 | 0.044±0.002 | 0.526±0.002 | 0.43±0.002 | 0.959±0.001 | 0.92±0.001 |
| | | 40 | 63.88±0 | 59.68±0.07 | 4.2±0.07 | 0.057±0.001 | 0.527±0.001 | 0.416±0.001 | 0.934±0.001 | 0.87±0.002 |
| | | 50 | 63.93±0 | 59.03±0.09 | 4.9±0.09 | 0.064±0.001 | 0.528±0.005 | 0.408±0.005 | 0.923±0.001 | 0.847±0.003 |
| | | 60 | 63.97±0 | 58.36±0.08 | 5.61±0.08 | 0.066±0.001 | 0.527±0.003 | 0.407±0.002 | 0.912±0.001 | 0.825±0.003 |
| | | 80 | 64.01±0 | 57.13±0.15 | 6.88±0.15 | 0.072±0.003 | 0.527±0.002 | 0.401±0.004 | 0.892±0.002 | 0.785±0.005 |
| | | 100 | 64.03±0 | 56.01±0.14 | 8.02±0.14 | 0.076±0.002 | 0.528±0.003 | 0.396±0.005 | 0.875±0.002 | 0.75±0.005 |
| | | 200 | 64.08±0 | 51.96±0.34 | 12.12±0.34 | 0.096±0.001 | 0.543±0.003 | 0.361±0.004 | 0.811±0.005 | 0.622±0.009 |
| | | 400 | 64.11±0 | 47.48±0.4 | 16.63±0.4 | 0.121±0.005 | 0.559±0.002 | 0.321±0.006 | 0.741±0.006 | 0.484±0.014 |
| | chr8 | 2.5 | 66.79±0 | 65.94±0.01 | 0.85±0.01 | 0.024±0.001 | 0.345±0.002 | 0.631±0.003 | 0.987±0 | 0.976±0 |
| | | 5 | 68.75±0 | 67.72±0.01 | 1.04±0.01 | 0.029±0 | 0.376±0.004 | 0.595±0.004 | 0.985±0 | 0.971±0 |
| | | 10 | 69.79±0 | 68.38±0.02 | 1.41±0.02 | 0.035±0.001 | 0.414±0.005 | 0.551±0.005 | 0.98±0 | 0.961±0.001 |
| | | 20 | 70.33±0 | 68.15±0.04 | 2.18±0.04 | 0.042±0.001 | 0.458±0.003 | 0.499±0.003 | 0.969±0.001 | 0.939±0.001 |
| | | 40 | 70.6±0 | 66.94±0.07 | 3.66±0.07 | 0.053±0.003 | 0.486±0.005 | 0.461±0.004 | 0.948±0.001 | 0.897±0.002 |
| | | 50 | 70.66±0 | 66.29±0.19 | 4.37±0.19 | 0.057±0.003 | 0.491±0.003 | 0.452±0.003 | 0.938±0.003 | 0.877±0.006 |
| | | 60 | 70.69±0 | 65.45±0.13 | 5.24±0.13 | 0.064±0.003 | 0.496±0.005 | 0.44±0.004 | 0.926±0.002 | 0.851±0.004 |
| | | 80 | 70.74±0 | 64.26±0.09 | 6.48±0.09 | 0.069±0.002 | 0.506±0.004 | 0.425±0.005 | 0.908±0.001 | 0.815±0.003 |
| | | 100 | 70.77±0 | 62.88±0.3 | 7.89±0.3 | 0.075±0.004 | 0.511±0.002 | 0.413±0.004 | 0.889±0.004 | 0.774±0.01 |
| | | 200 | 70.82±0 | 58.35±0.38 | 12.47±0.38 | 0.095±0.004 | 0.522±0.005 | 0.383±0.009 | 0.824±0.005 | 0.642±0.012 |
| | | 400 | 70.85±0 | 53.07±0.23 | 17.78±0.23 | 0.121±0.002 | 0.547±0.01 | 0.332±0.01 | 0.749±0.003 | 0.494±0.007 |
| | chr9 | 2.5 | 39.7±0 | 39.02±0.01 | 0.68±0.01 | 0.026±0.001 | 0.444±0.002 | 0.529±0.002 | 0.983±0 | 0.968±0 |
| | | 5 | 40.76±0 | 39.97±0 | 0.8±0 | 0.03±0.001 | 0.46±0.003 | 0.51±0.004 | 0.98±0 | 0.963±0 |
| | | 10 | 41.31±0 | 40.25±0.03 | 1.06±0.03 | 0.036±0.002 | 0.481±0.003 | 0.483±0.004 | 0.974±0.001 | 0.951±0.002 |
| | | 20 | 41.59±0 | 40.03±0.03 | 1.56±0.03 | 0.045±0.003 | 0.502±0.003 | 0.453±0.004 | 0.962±0.001 | 0.927±0.002 |
| | | 40 | 41.73±0 | 39.18±0.07 | 2.55±0.07 | 0.058±0.003 | 0.507±0.007 | 0.434±0.008 | 0.939±0.002 | 0.878±0.004 |
| | | 50 | 41.76±0 | 38.7±0.08 | 3.05±0.08 | 0.059±0.003 | 0.511±0.005 | 0.43±0.003 | 0.927±0.002 | 0.855±0.004 |
| | | 60 | 41.78±0 | 38.3±0.1 | 3.47±0.1 | 0.066±0.003 | 0.506±0.007 | 0.427±0.007 | 0.917±0.002 | 0.833±0.005 |
| | | 80 | 41.8±0 | 37.43±0.08 | 4.37±0.08 | 0.074±0.004 | 0.511±0.006 | 0.416±0.008 | 0.896±0.002 | 0.789±0.004 |
| | | 100 | 41.81±0 | 36.71±0.16 | 5.1±0.16 | 0.079±0.003 | 0.512±0.003 | 0.409±0.006 | 0.878±0.004 | 0.753±0.007 |
| | | 200 | 41.84±0 | 34.06±0.23 | 7.78±0.23 | 0.101±0.005 | 0.519±0.002 | 0.381±0.004 | 0.814±0.005 | 0.621±0.012 |
| | | 400 | 41.86±0 | 30.63±0.27 | 11.22±0.27 | 0.126±0.005 | 0.533±0.004 | 0.341±0.005 | 0.732±0.006 | 0.458±0.014 |

| | | | | | | | | | | |
|---|---|---|---|---|---|---|---|---|---|---|
| | chr10 | 2.5 | 49.44±0 | 48.58±0.01 | 0.86±0.01 | 0.029±0.001 | 0.417±0.003 | 0.554±0.003 | 0.983±0 | 0.967±0 |
| | | 5 | 50.51±0 | 49.43±0.01 | 1.08±0.01 | 0.034±0.001 | 0.442±0.002 | 0.524±0.003 | 0.979±0 | 0.96±0 |
| | | 10 | 51.06±0 | 49.53±0.03 | 1.53±0.03 | 0.044±0.001 | 0.469±0.002 | 0.486±0.002 | 0.97±0 | 0.942±0.001 |
| | | 20 | 51.33±0 | 48.94±0.03 | 2.4±0.03 | 0.052±0.003 | 0.484±0.004 | 0.464±0.006 | 0.953±0.001 | 0.909±0.001 |
| | | 40 | 51.47±0 | 47.38±0.1 | 4.1±0.1 | 0.066±0.003 | 0.5±0.002 | 0.435±0.004 | 0.92±0.002 | 0.842±0.005 |
| | | 50 | 51.5±0 | 46.55±0.12 | 4.95±0.12 | 0.07±0.002 | 0.506±0.006 | 0.425±0.004 | 0.904±0.002 | 0.809±0.006 |
| | | 60 | 51.52±0 | 45.93±0.17 | 5.59±0.17 | 0.07±0.002 | 0.509±0.005 | 0.422±0.005 | 0.892±0.003 | 0.786±0.007 |
| | | 80 | 51.54±0 | 44.61±0.22 | 6.94±0.22 | 0.078±0.003 | 0.516±0.006 | 0.406±0.007 | 0.865±0.004 | 0.733±0.009 |
| | | 100 | 51.56±0 | 43.41±0.3 | 8.15±0.3 | 0.086±0.002 | 0.512±0.005 | 0.401±0.004 | 0.842±0.006 | 0.685±0.011 |
| | | 200 | 51.59±0 | 39.33±0.34 | 12.26±0.34 | 0.107±0.004 | 0.527±0.007 | 0.366±0.008 | 0.762±0.007 | 0.531±0.014 |
| | | 400 | 51.6±0 | 35.53±0.26 | 16.07±0.26 | 0.138±0.003 | 0.547±0.006 | 0.315±0.007 | 0.688±0.005 | 0.388±0.008 |
| | chr11 | 2.5 | 49.87±0 | 49.3±0 | 0.57±0 | 0.022±0.001 | 0.305±0.003 | 0.673±0.003 | 0.989±0 | 0.979±0 |
| | | 5 | 51.1±0 | 50.38±0.01 | 0.72±0.01 | 0.027±0.001 | 0.353±0.004 | 0.62±0.003 | 0.986±0 | 0.973±0 |
| | | 10 | 51.73±0 | 50.72±0.02 | 1.01±0.02 | 0.032±0.002 | 0.405±0.005 | 0.563±0.004 | 0.98±0 | 0.962±0.001 |
| | | 20 | 52.05±0 | 50.31±0.04 | 1.73±0.04 | 0.04±0.001 | 0.454±0.008 | 0.506±0.008 | 0.967±0.001 | 0.935±0.001 |
| | | 40 | 52.2±0 | 49.23±0.06 | 2.98±0.06 | 0.049±0.002 | 0.492±0.004 | 0.459±0.004 | 0.943±0.001 | 0.887±0.002 |
| | | 50 | 52.24±0 | 48.67±0.07 | 3.57±0.07 | 0.055±0.003 | 0.5±0.005 | 0.446±0.004 | 0.932±0.001 | 0.864±0.003 |
| | | 60 | 52.26±0 | 48.18±0.12 | 4.07±0.12 | 0.057±0.004 | 0.502±0.001 | 0.441±0.005 | 0.922±0.002 | 0.845±0.005 |
| | | 80 | 52.28±0 | 47.19±0.14 | 5.09±0.14 | 0.063±0.004 | 0.513±0.003 | 0.424±0.006 | 0.903±0.003 | 0.805±0.006 |
| | | 100 | 52.3±0 | 46.29±0.18 | 6.01±0.18 | 0.07±0.003 | 0.514±0.003 | 0.416±0.004 | 0.885±0.003 | 0.769±0.007 |
| | | 200 | 52.33±0 | 42.9±0.32 | 9.44±0.32 | 0.092±0.005 | 0.535±0.005 | 0.373±0.008 | 0.82±0.006 | 0.635±0.013 |
| | | 400 | 52.35±0 | 39.19±0.27 | 13.16±0.27 | 0.115±0.002 | 0.559±0.005 | 0.326±0.006 | 0.749±0.005 | 0.495±0.008 |
| | chr12 | 2.5 | 34.37±0 | 33.7±0.01 | 0.67±0.01 | 0.029±0.001 | 0.413±0.003 | 0.559±0.004 | 0.981±0 | 0.963±0 |
| | | 5 | 35.29±0 | 34.42±0.01 | 0.88±0.01 | 0.037±0.003 | 0.443±0.002 | 0.52±0.003 | 0.975±0 | 0.951±0.001 |
| | | 10 | 35.77±0 | 34.51±0.02 | 1.26±0.02 | 0.045±0.003 | 0.473±0.005 | 0.483±0.006 | 0.965±0.001 | 0.93±0.002 |
| | | 20 | 36.02±0 | 34.01±0.05 | 2.01±0.05 | 0.054±0.002 | 0.5±0.005 | 0.446±0.005 | 0.944±0.002 | 0.888±0.003 |
| | | 40 | 36.14±0 | 32.87±0.06 | 3.27±0.06 | 0.064±0.003 | 0.51±0.002 | 0.426±0.003 | 0.909±0.002 | 0.817±0.003 |
| | | 50 | 36.16±0 | 32.3±0.12 | 3.86±0.12 | 0.068±0.003 | 0.517±0.004 | 0.415±0.006 | 0.893±0.003 | 0.784±0.007 |
| | | 60 | 36.18±0 | 31.77±0.1 | 4.41±0.1 | 0.071±0.004 | 0.52±0.004 | 0.409±0.007 | 0.878±0.003 | 0.755±0.007 |
| | | 80 | 36.2±0 | 30.9±0.09 | 5.3±0.09 | 0.077±0.003 | 0.524±0.004 | 0.398±0.004 | 0.854±0.002 | 0.705±0.005 |
| | | 100 | 36.21±0 | 30.15±0.17 | 6.06±0.17 | 0.082±0.004 | 0.527±0.008 | 0.391±0.008 | 0.833±0.005 | 0.663±0.01 |
| | | 200 | 36.24±0 | 27.25±0.33 | 8.99±0.33 | 0.103±0.007 | 0.532±0.005 | 0.366±0.008 | 0.752±0.009 | 0.507±0.022 |
| | | 400 | 36.25±0 | 24.52±0.23 | 11.73±0.23 | 0.13±0.007 | 0.548±0.004 | 0.321±0.01 | 0.677±0.006 | 0.363±0.015 |
| | chr13 | 2.5 | 67.15±0 | 66.34±0.01 | 0.81±0.01 | 0.023±0.001 | 0.318±0.003 | 0.659±0.003 | 0.988±0 | 0.979±0 |
| | | 5 | 69.99±0 | 69.04±0 | 0.95±0 | 0.027±0.001 | 0.342±0.002 | 0.631±0.003 | 0.986±0 | 0.976±0 |

| | | | | | | | | | | |
|---|---|---|---|---|---|---|---|---|---|---|
| | | 10 | 71.55±0 | 70.36±0.01 | 1.19±0.01 | 0.032±0.002 | 0.377±0.003 | 0.591±0.004 | 0.983±0 | 0.971±0 |
| | | 20 | 72.36±0 | 70.67±0.02 | 1.69±0.02 | 0.041±0.002 | 0.423±0.006 | 0.536±0.005 | 0.977±0 | 0.958±0.001 |
| | | 40 | 72.77±0 | 70.03±0.06 | 2.74±0.06 | 0.054±0.004 | 0.455±0.006 | 0.492±0.006 | 0.962±0.001 | 0.931±0.002 |
| | | 50 | 72.86±0 | 69.64±0.08 | 3.21±0.08 | 0.057±0.001 | 0.465±0.006 | 0.478±0.007 | 0.956±0.001 | 0.918±0.002 |
| | | 60 | 72.91±0 | 69.13±0.12 | 3.79±0.12 | 0.063±0.002 | 0.471±0.006 | 0.466±0.006 | 0.948±0.002 | 0.903±0.003 |
| | | 80 | 72.98±0 | 68.15±0.14 | 4.83±0.14 | 0.072±0.005 | 0.477±0.005 | 0.451±0.007 | 0.934±0.002 | 0.874±0.005 |
| | | 100 | 73.02±0 | 67.44±0.13 | 5.59±0.13 | 0.076±0.001 | 0.482±0.003 | 0.442±0.004 | 0.924±0.002 | 0.854±0.003 |
| | | 200 | 73.11±0 | 63.88±0.18 | 9.23±0.18 | 0.104±0.005 | 0.496±0.005 | 0.4±0.009 | 0.874±0.002 | 0.752±0.006 |
| | | 400 | 73.15±0 | 59.53±0.22 | 13.62±0.22 | 0.131±0.003 | 0.514±0.004 | 0.354±0.007 | 0.814±0.003 | 0.631±0.006 |
| | chr14 | 2.5 | 62.71±0 | 61.79±0.01 | 0.92±0.01 | 0.022±0.001 | 0.379±0.003 | 0.599±0.004 | 0.985±0 | 0.973±0 |
| | | 5 | 64.71±0 | 63.58±0.01 | 1.13±0.01 | 0.029±0.001 | 0.403±0.005 | 0.568±0.004 | 0.983±0 | 0.967±0 |
| | | 10 | 65.74±0 | 64.25±0.01 | 1.5±0.01 | 0.036±0.001 | 0.438±0.004 | 0.526±0.004 | 0.977±0 | 0.957±0 |
| | | 20 | 66.27±0 | 64.06±0.03 | 2.21±0.03 | 0.043±0.001 | 0.466±0.004 | 0.491±0.005 | 0.967±0 | 0.936±0.001 |
| | | 40 | 66.54±0 | 63.06±0.11 | 3.48±0.11 | 0.052±0.002 | 0.489±0.005 | 0.459±0.005 | 0.948±0.002 | 0.898±0.003 |
| | | 50 | 66.59±0 | 62.42±0.09 | 4.17±0.09 | 0.056±0.002 | 0.497±0.006 | 0.447±0.006 | 0.937±0.001 | 0.877±0.003 |
| | | 60 | 66.62±0 | 61.86±0.19 | 4.76±0.19 | 0.06±0.002 | 0.496±0.005 | 0.444±0.007 | 0.929±0.003 | 0.859±0.006 |
| | | 80 | 66.67±0 | 60.72±0.18 | 5.95±0.18 | 0.067±0.002 | 0.505±0.005 | 0.428±0.006 | 0.911±0.003 | 0.822±0.006 |
| | | 100 | 66.7±0 | 59.97±0.06 | 6.73±0.06 | 0.071±0.002 | 0.512±0.006 | 0.417±0.007 | 0.899±0.001 | 0.798±0.002 |
| | | 200 | 66.75±0 | 56.05±0.22 | 10.7±0.22 | 0.088±0.003 | 0.529±0.006 | 0.383±0.005 | 0.84±0.003 | 0.68±0.007 |
| | | 400 | 66.78±0 | 51.48±0.3 | 15.3±0.3 | 0.116±0.001 | 0.549±0.006 | 0.335±0.007 | 0.771±0.005 | 0.54±0.007 |
| | chr15 | 2.5 | 52.16±0 | 51.48±0 | 0.68±0 | 0.018±0 | 0.393±0.004 | 0.588±0.003 | 0.987±0 | 0.977±0 |
| | | 5 | 53.99±0 | 53.19±0.01 | 0.8±0.01 | 0.02±0.001 | 0.417±0.003 | 0.563±0.004 | 0.985±0 | 0.974±0 |
| | | 10 | 55.01±0 | 53.99±0.01 | 1.02±0.01 | 0.023±0.001 | 0.443±0.002 | 0.535±0.002 | 0.981±0 | 0.967±0.001 |
| | | 20 | 55.55±0 | 54.03±0.03 | 1.52±0.03 | 0.029±0.002 | 0.477±0.006 | 0.494±0.007 | 0.973±0.001 | 0.95±0.001 |
| | | 40 | 55.83±0 | 53.32±0.04 | 2.51±0.04 | 0.036±0.002 | 0.501±0.004 | 0.463±0.003 | 0.955±0.001 | 0.918±0.001 |
| | | 50 | 55.89±0 | 52.84±0.1 | 3.05±0.1 | 0.041±0.003 | 0.507±0.004 | 0.452±0.005 | 0.945±0.002 | 0.899±0.004 |
| | | 60 | 55.93±0 | 52.37±0.09 | 3.56±0.09 | 0.042±0.001 | 0.515±0.006 | 0.443±0.006 | 0.936±0.002 | 0.883±0.003 |
| | | 80 | 55.97±0 | 51.52±0.11 | 4.46±0.11 | 0.048±0.001 | 0.52±0.008 | 0.432±0.009 | 0.92±0.002 | 0.852±0.004 |
| | | 100 | 56±0 | 50.7±0.2 | 5.31±0.2 | 0.052±0.002 | 0.523±0.005 | 0.425±0.006 | 0.905±0.004 | 0.824±0.007 |
| | | 200 | 56.06±0 | 47.01±0.22 | 9.05±0.22 | 0.077±0.002 | 0.532±0.005 | 0.391±0.007 | 0.839±0.004 | 0.694±0.007 |
| | | 400 | 56.09±0 | 43.17±0.29 | 12.92±0.29 | 0.105±0.003 | 0.554±0.006 | 0.341±0.008 | 0.77±0.005 | 0.56±0.01 |
| | chr16 | 2.5 | 40.93±0 | 40.42±0.01 | 0.52±0.01 | 0.027±0.001 | 0.319±0.001 | 0.654±0.001 | 0.987±0 | 0.978±0 |
| | | 5 | 42.04±0 | 41.4±0.01 | 0.64±0.01 | 0.033±0.001 | 0.356±0.004 | 0.611±0.006 | 0.985±0 | 0.973±0 |
| | | 10 | 42.64±0 | 41.76±0.02 | 0.88±0.02 | 0.039±0.001 | 0.396±0.004 | 0.565±0.004 | 0.979±0 | 0.963±0.001 |
| | | 20 | 42.95±0 | 41.57±0.05 | 1.39±0.05 | 0.052±0.005 | 0.436±0.004 | 0.513±0.003 | 0.968±0.001 | 0.941±0.003 |

| | | | | | | | | | | |
|---|---|---|---|---|---|---|---|---|---|---|
| | | 40 | 43.11±0 | 40.8±0.05 | 2.31±0.05 | 0.063±0.002 | 0.467±0.002 | 0.47±0.003 | 0.946±0.001 | 0.901±0.002 |
| | | 50 | 43.15±0 | 40.32±0.1 | 2.83±0.1 | 0.072±0.004 | 0.476±0.007 | 0.452±0.008 | 0.935±0.002 | 0.877±0.005 |
| | | 60 | 43.17±0 | 39.86±0.09 | 3.31±0.09 | 0.073±0.001 | 0.489±0.009 | 0.439±0.01 | 0.923±0.002 | 0.856±0.004 |
| | | 80 | 43.19±0 | 39.01±0.22 | 4.19±0.22 | 0.081±0.007 | 0.494±0.006 | 0.425±0.005 | 0.903±0.005 | 0.817±0.012 |
| | | 100 | 43.21±0 | 38.34±0.1 | 4.87±0.1 | 0.084±0.002 | 0.494±0.004 | 0.422±0.006 | 0.887±0.002 | 0.788±0.005 |
| | | 200 | 43.24±0 | 35.61±0.11 | 7.63±0.11 | 0.104±0.006 | 0.508±0.003 | 0.388±0.007 | 0.824±0.002 | 0.667±0.007 |
| | | 400 | 43.26±0 | 32.23±0.26 | 11.03±0.26 | 0.127±0.002 | 0.522±0.006 | 0.351±0.006 | 0.745±0.006 | 0.525±0.011 |
| | chr17 | 2.5 | 35.53±0 | 34.97±0.01 | 0.56±0.01 | 0.032±0.002 | 0.402±0.004 | 0.566±0.005 | 0.984±0 | 0.971±0 |
| | | 5 | 36.47±0 | 35.78±0.01 | 0.69±0.01 | 0.042±0.002 | 0.426±0.007 | 0.532±0.007 | 0.981±0 | 0.964±0.001 |
| | | 10 | 36.97±0 | 35.99±0.02 | 0.97±0.02 | 0.054±0.004 | 0.457±0.005 | 0.489±0.004 | 0.974±0 | 0.948±0.001 |
| | | 20 | 37.22±0 | 35.77±0.04 | 1.45±0.04 | 0.061±0.005 | 0.477±0.003 | 0.462±0.007 | 0.961±0.001 | 0.923±0.003 |
| | | 40 | 37.35±0 | 34.92±0.03 | 2.42±0.03 | 0.068±0.003 | 0.497±0.007 | 0.434±0.009 | 0.935±0.001 | 0.871±0.002 |
| | | 50 | 37.37±0 | 34.48±0.07 | 2.89±0.07 | 0.072±0.001 | 0.499±0.007 | 0.429±0.007 | 0.923±0.002 | 0.846±0.004 |
| | | 60 | 37.39±0 | 34.14±0.05 | 3.25±0.05 | 0.076±0.002 | 0.506±0.005 | 0.418±0.005 | 0.913±0.001 | 0.826±0.003 |
| | | 80 | 37.41±0 | 33.32±0.14 | 4.09±0.14 | 0.08±0.004 | 0.511±0.008 | 0.409±0.009 | 0.891±0.004 | 0.782±0.007 |
| | | 100 | 37.42±0 | 32.69±0.14 | 4.73±0.14 | 0.082±0.003 | 0.51±0.002 | 0.408±0.003 | 0.874±0.004 | 0.75±0.007 |
| | | 200 | 37.45±0 | 29.81±0.12 | 7.64±0.12 | 0.099±0.002 | 0.524±0.004 | 0.377±0.005 | 0.796±0.003 | 0.601±0.007 |
| | | 400 | 37.46±0 | 26.25±0.31 | 11.21±0.31 | 0.122±0.004 | 0.536±0.008 | 0.341±0.006 | 0.701±0.008 | 0.429±0.016 |
| | chr18 | 2.5 | 26.11±0 | 25.75±0 | 0.36±0 | 0.022±0.001 | 0.394±0.005 | 0.584±0.005 | 0.986±0 | 0.975±0 |
| | | 5 | 26.88±0 | 26.44±0.01 | 0.44±0.01 | 0.024±0.001 | 0.422±0.006 | 0.554±0.006 | 0.984±0 | 0.97±0.001 |
| | | 10 | 27.29±0 | 26.71±0.01 | 0.58±0.01 | 0.027±0.001 | 0.456±0.005 | 0.517±0.005 | 0.979±0 | 0.96±0.001 |
| | | 20 | 27.51±0 | 26.62±0.01 | 0.89±0.01 | 0.033±0.001 | 0.49±0.006 | 0.477±0.006 | 0.968±0.001 | 0.939±0.001 |
| | | 40 | 27.63±0 | 26.07±0.04 | 1.55±0.04 | 0.041±0.004 | 0.52±0.007 | 0.439±0.004 | 0.944±0.001 | 0.893±0.003 |
| | | 50 | 27.65±0 | 25.73±0.05 | 1.92±0.05 | 0.046±0.004 | 0.523±0.007 | 0.431±0.007 | 0.931±0.002 | 0.867±0.005 |
| | | 60 | 27.66±0 | 25.39±0.17 | 2.28±0.17 | 0.053±0.004 | 0.525±0.011 | 0.422±0.011 | 0.918±0.006 | 0.84±0.012 |
| | | 80 | 27.68±0 | 24.97±0.09 | 2.71±0.09 | 0.056±0.003 | 0.528±0.01 | 0.416±0.011 | 0.902±0.003 | 0.81±0.006 |
| | | 100 | 27.69±0 | 24.37±0.11 | 3.33±0.11 | 0.063±0.003 | 0.525±0.005 | 0.413±0.003 | 0.88±0.004 | 0.766±0.008 |
| | | 200 | 27.72±0 | 22.59±0.14 | 5.13±0.14 | 0.088±0.005 | 0.533±0.004 | 0.379±0.005 | 0.815±0.005 | 0.633±0.012 |
| | | 400 | 27.73±0 | 20.48±0.2 | 7.25±0.2 | 0.121±0.005 | 0.549±0.008 | 0.33±0.01 | 0.739±0.007 | 0.477±0.013 |
| Test dataset 2 | chr1 | 2.5 | 181.47±0 | 177.16±0.02 | 4.31±0.02 | 0.019±0 | 0.444±0.002 | 0.536±0.002 | 0.976±0 | 0.958±0 |
| | | 5 | 191.64±0 | 186.76±0.02 | 4.88±0.02 | 0.02±0 | 0.436±0.001 | 0.544±0.002 | 0.975±0 | 0.955±0 |
| | | 10 | 197.8±0 | 192.15±0.02 | 5.65±0.02 | 0.022±0 | 0.437±0.002 | 0.542±0.002 | 0.971±0 | 0.949±0 |
| | | 20 | 201.33±0 | 194.59±0.06 | 6.74±0.06 | 0.025±0 | 0.445±0.002 | 0.53±0.002 | 0.967±0 | 0.94±0.001 |
| | | 40 | 203.24±0 | 194.48±0.18 | 8.76±0.18 | 0.032±0.001 | 0.466±0.004 | 0.502±0.005 | 0.957±0.001 | 0.922±0.002 |
| | | 50 | 203.63±0 | 193.76±0.18 | 9.86±0.18 | 0.036±0.001 | 0.474±0.005 | 0.49±0.004 | 0.952±0.001 | 0.912±0.002 |

| | | | | | | | | | | |
|---|---|---|---|---|---|---|---|---|---|---|
| | | 60 | 203.89±0 | 192.94±0.1 | 10.96±0.1 | 0.04±0.001 | 0.48±0.005 | 0.479±0.006 | 0.946±0 | 0.901±0.001 |
| | | 80 | 204.23±0 | 191.37±0.34 | 12.86±0.34 | 0.044±0.002 | 0.489±0.004 | 0.466±0.004 | 0.937±0.002 | 0.884±0.004 |
| | | 100 | 204.43±0 | 189.8±0.18 | 14.63±0.18 | 0.05±0.001 | 0.493±0.004 | 0.457±0.004 | 0.928±0.001 | 0.866±0.002 |
| | | 200 | 204.84±0 | 182.89±0.79 | 21.95±0.79 | 0.066±0.006 | 0.506±0.006 | 0.428±0.006 | 0.893±0.004 | 0.796±0.01 |
| | | 400 | 205.04±0 | 171.22±0.83 | 33.82±0.83 | 0.091±0.007 | 0.515±0.008 | 0.394±0.011 | 0.835±0.004 | 0.68±0.011 |
| | chr2 | 2.5 | 132.22±0 | 128.82±0.01 | 3.4±0.01 | 0.02±0 | 0.431±0.002 | 0.549±0.002 | 0.974±0 | 0.954±0 |
| | | 5 | 138.31±0 | 134.46±0.01 | 3.85±0.01 | 0.02±0 | 0.424±0.003 | 0.556±0.003 | 0.972±0 | 0.951±0 |
| | | 10 | 141.79±0 | 137.39±0.03 | 4.4±0.03 | 0.021±0 | 0.426±0.001 | 0.552±0.001 | 0.969±0 | 0.945±0 |
| | | 20 | 143.71±0 | 138.37±0.08 | 5.34±0.08 | 0.024±0.001 | 0.438±0.005 | 0.538±0.005 | 0.963±0.001 | 0.934±0.001 |
| | | 40 | 144.73±0 | 137.5±0.16 | 7.23±0.16 | 0.03±0.001 | 0.465±0.004 | 0.505±0.005 | 0.95±0.001 | 0.91±0.002 |
| | | 50 | 144.93±0 | 136.78±0.07 | 8.16±0.07 | 0.033±0.001 | 0.475±0.004 | 0.491±0.004 | 0.944±0 | 0.898±0.001 |
| | | 60 | 145.07±0 | 136.02±0.05 | 9.05±0.05 | 0.036±0.001 | 0.482±0.003 | 0.482±0.003 | 0.938±0 | 0.887±0.001 |
| | | 80 | 145.25±0 | 134.34±0.26 | 10.9±0.26 | 0.04±0.003 | 0.489±0.003 | 0.471±0.002 | 0.925±0.002 | 0.863±0.004 |
| | | 100 | 145.36±0 | 132.75±0.28 | 12.61±0.28 | 0.044±0.001 | 0.498±0.006 | 0.458±0.006 | 0.913±0.002 | 0.841±0.004 |
| | | 200 | 145.57±0 | 125.28±0.64 | 20.29±0.64 | 0.063±0.003 | 0.511±0.004 | 0.426±0.002 | 0.861±0.004 | 0.74±0.009 |
| | | 400 | 145.68±0 | 113.96±0.73 | 31.72±0.73 | 0.092±0.005 | 0.515±0.002 | 0.393±0.004 | 0.782±0.005 | 0.59±0.011 |
| | chr3 | 2.5 | 129.43±0 | 126.36±0.01 | 3.07±0.01 | 0.019±0 | 0.431±0.001 | 0.549±0.001 | 0.976±0 | 0.955±0 |
| | | 5 | 134.99±0 | 131.53±0.01 | 3.45±0.01 | 0.02±0 | 0.423±0.002 | 0.557±0.002 | 0.974±0 | 0.951±0 |
| | | 10 | 138.17±0 | 134.24±0.03 | 3.93±0.03 | 0.022±0.001 | 0.426±0.003 | 0.553±0.004 | 0.972±0 | 0.945±0 |
| | | 20 | 139.92±0 | 135.21±0.02 | 4.71±0.02 | 0.025±0 | 0.44±0.003 | 0.535±0.003 | 0.966±0 | 0.935±0 |
| | | 40 | 140.84±0 | 134.5±0.09 | 6.34±0.09 | 0.03±0 | 0.47±0.005 | 0.5±0.005 | 0.955±0.001 | 0.912±0.001 |
| | | 50 | 141.03±0 | 133.68±0.28 | 7.35±0.28 | 0.035±0.003 | 0.477±0.004 | 0.488±0.003 | 0.948±0.002 | 0.897±0.004 |
| | | 60 | 141.16±0 | 133.07±0.1 | 8.09±0.1 | 0.036±0.002 | 0.482±0.002 | 0.482±0.004 | 0.943±0.001 | 0.887±0.002 |
| | | 80 | 141.32±0 | 131.7±0.16 | 9.62±0.16 | 0.041±0.003 | 0.492±0.005 | 0.467±0.006 | 0.932±0.001 | 0.865±0.003 |
| | | 100 | 141.42±0 | 129.69±0.56 | 11.73±0.56 | 0.049±0.003 | 0.495±0.004 | 0.456±0.007 | 0.917±0.004 | 0.833±0.009 |
| | | 200 | 141.61±0 | 122.35±1.01 | 19.27±1.01 | 0.069±0.006 | 0.512±0.003 | 0.419±0.009 | 0.864±0.007 | 0.722±0.017 |
| | | 400 | 141.71±0 | 111.64±0.77 | 30.07±0.77 | 0.101±0.008 | 0.522±0.006 | 0.377±0.005 | 0.788±0.005 | 0.559±0.016 |
| | chr4 | 2.5 | 138.7±0 | 135.31±0.01 | 3.38±0.01 | 0.019±0 | 0.433±0.001 | 0.548±0.001 | 0.976±0 | 0.954±0 |
| | | 5 | 144.47±0 | 140.66±0.01 | 3.81±0.01 | 0.019±0 | 0.424±0 | 0.557±0 | 0.974±0 | 0.95±0 |
| | | 10 | 147.74±0 | 143.4±0.03 | 4.33±0.03 | 0.019±0.001 | 0.423±0.003 | 0.557±0.003 | 0.971±0 | 0.945±0 |
| | | 20 | 149.52±0 | 144.21±0.04 | 5.3±0.04 | 0.022±0.001 | 0.44±0.003 | 0.538±0.003 | 0.965±0 | 0.933±0.001 |
| | | 40 | 150.44±0 | 143.02±0.16 | 7.42±0.16 | 0.028±0.001 | 0.469±0.004 | 0.502±0.005 | 0.951±0.001 | 0.906±0.002 |
| | | 50 | 150.63±0 | 142.21±0.2 | 8.42±0.2 | 0.031±0.002 | 0.479±0.006 | 0.49±0.006 | 0.944±0.001 | 0.893±0.003 |
| | | 60 | 150.76±0 | 141.2±0.14 | 9.56±0.14 | 0.035±0.001 | 0.484±0.006 | 0.481±0.006 | 0.937±0.001 | 0.878±0.002 |
| | | 80 | 150.92±0 | 139.29±0.35 | 11.63±0.35 | 0.039±0.003 | 0.489±0.008 | 0.472±0.008 | 0.923±0.002 | 0.851±0.005 |

| | | | | | | | | | | |
|---|---|---|---|---|---|---|---|---|---|---|
| | | 100 | 151.01±0 | 137.93±0.23 | 13.08±0.23 | 0.041±0.002 | 0.498±0.007 | 0.461±0.007 | 0.913±0.002 | 0.832±0.003 |
| | | 200 | 151.21±0 | 129.14±1.12 | 22.06±1.12 | 0.062±0.004 | 0.512±0.002 | 0.426±0.004 | 0.854±0.007 | 0.713±0.016 |
| | | 400 | 151.3±0 | 116.54±0.53 | 34.76±0.53 | 0.095±0.004 | 0.533±0.008 | 0.372±0.006 | 0.77±0.004 | 0.543±0.009 |
| | chr5 | 2.5 | 101.71±0 | 99.35±0 | 2.36±0 | 0.019±0 | 0.444±0.001 | 0.538±0.001 | 0.977±0 | 0.959±0 |
| | | 5 | 105.77±0 | 103.04±0.02 | 2.72±0.02 | 0.02±0.001 | 0.442±0.002 | 0.539±0.003 | 0.974±0 | 0.955±0 |
| | | 10 | 108.16±0 | 104.92±0.03 | 3.24±0.03 | 0.022±0.001 | 0.446±0.004 | 0.532±0.004 | 0.97±0 | 0.947±0.001 |
| | | 20 | 109.5±0 | 105.4±0.05 | 4.09±0.05 | 0.025±0 | 0.467±0.005 | 0.508±0.005 | 0.963±0 | 0.934±0.001 |
| | | 40 | 110.22±0 | 104.09±0.11 | 6.14±0.11 | 0.035±0.001 | 0.495±0.006 | 0.47±0.006 | 0.944±0.001 | 0.899±0.002 |
| | | 50 | 110.37±0 | 103.52±0.15 | 6.86±0.15 | 0.036±0.001 | 0.505±0.007 | 0.459±0.007 | 0.938±0.001 | 0.888±0.003 |
| | | 60 | 110.47±0 | 102.59±0.22 | 7.88±0.22 | 0.04±0.001 | 0.505±0.004 | 0.455±0.004 | 0.929±0.002 | 0.87±0.004 |
| | | 80 | 110.6±0 | 101.15±0.18 | 9.46±0.18 | 0.044±0.001 | 0.511±0.007 | 0.445±0.008 | 0.915±0.002 | 0.844±0.003 |
| | | 100 | 110.68±0 | 99.86±0.17 | 10.82±0.17 | 0.047±0.002 | 0.515±0.003 | 0.437±0.003 | 0.902±0.002 | 0.822±0.003 |
| | | 200 | 110.83±0 | 93.61±0.69 | 17.22±0.69 | 0.064±0.002 | 0.524±0.006 | 0.412±0.006 | 0.845±0.006 | 0.714±0.011 |
| | | 400 | 110.91±0 | 85.58±0.68 | 25.33±0.68 | 0.093±0.007 | 0.541±0.007 | 0.366±0.006 | 0.772±0.006 | 0.575±0.015 |
| | chr6 | 2.5 | 148.9±0 | 145.39±0.01 | 3.51±0.01 | 0.019±0 | 0.437±0.002 | 0.545±0.002 | 0.976±0 | 0.961±0 |
| | | 5 | 156.42±0 | 152.43±0.01 | 3.99±0.01 | 0.019±0 | 0.428±0.002 | 0.553±0.001 | 0.974±0 | 0.957±0 |
| | | 10 | 160.69±0 | 156.12±0.02 | 4.57±0.02 | 0.02±0.001 | 0.429±0.002 | 0.551±0.002 | 0.972±0 | 0.952±0 |
| | | 20 | 163.01±0 | 157.54±0.03 | 5.47±0.03 | 0.024±0 | 0.443±0.001 | 0.533±0.002 | 0.966±0 | 0.943±0 |
| | | 40 | 164.23±0 | 157.02±0.06 | 7.21±0.06 | 0.028±0.001 | 0.469±0.002 | 0.503±0.002 | 0.956±0 | 0.925±0.001 |
| | | 50 | 164.47±0 | 156.34±0.09 | 8.13±0.09 | 0.031±0.001 | 0.478±0.005 | 0.492±0.004 | 0.951±0.001 | 0.916±0.001 |
| | | 60 | 164.64±0 | 155.57±0.22 | 9.07±0.22 | 0.036±0.001 | 0.485±0.003 | 0.48±0.003 | 0.945±0.001 | 0.905±0.002 |
| | | 80 | 164.85±0 | 154.32±0.28 | 10.53±0.28 | 0.038±0.002 | 0.493±0.002 | 0.469±0.004 | 0.936±0.002 | 0.89±0.003 |
| | | 100 | 164.97±0 | 152.22±0.28 | 12.75±0.28 | 0.044±0.002 | 0.497±0.004 | 0.459±0.004 | 0.923±0.002 | 0.866±0.003 |
| | | 200 | 165.23±0 | 145.37±0.67 | 19.86±0.67 | 0.06±0.002 | 0.514±0.001 | 0.425±0.002 | 0.88±0.004 | 0.787±0.008 |
| | | 400 | 165.35±0 | 134.25±0.56 | 31.1±0.56 | 0.085±0.003 | 0.518±0.007 | 0.397±0.005 | 0.812±0.003 | 0.662±0.007 |
| | chr7 | 2.5 | 118.72±26.9 | 115.33±26.06 | 3.39±0.84 | 0.022±0.001 | 0.451±0.015 | 0.527±0.016 | 0.972±0.001 | 0.947±0.002 |
| | | 5 | 136.21±0 | 132±0.06 | 4.21±0.06 | 0.023±0 | 0.448±0.006 | 0.529±0.007 | 0.969±0 | 0.942±0.001 |
| | | 10 | 139.26±0 | 134.41±0.02 | 4.85±0.02 | 0.024±0 | 0.452±0.004 | 0.524±0.004 | 0.965±0 | 0.935±0 |
| | | 20 | 140.9±0 | 135±0.1 | 5.9±0.1 | 0.028±0.001 | 0.464±0.002 | 0.508±0.003 | 0.958±0.001 | 0.921±0.002 |
| | | 40 | 141.76±0 | 133.79±0.09 | 7.97±0.09 | 0.035±0.001 | 0.481±0.005 | 0.483±0.004 | 0.944±0.001 | 0.893±0.001 |
| | | 50 | 141.93±0 | 133±0.15 | 8.93±0.15 | 0.038±0.001 | 0.494±0.004 | 0.468±0.004 | 0.937±0.001 | 0.88±0.002 |
| | | 60 | 142.05±0 | 132±0.36 | 10.05±0.36 | 0.042±0.002 | 0.497±0.007 | 0.461±0.008 | 0.929±0.003 | 0.864±0.005 |
| | | 80 | 142.2±0 | 130.14±0.17 | 12.06±0.17 | 0.045±0.002 | 0.504±0.006 | 0.451±0.008 | 0.915±0.001 | 0.837±0.002 |
| | | 100 | 142.28±0 | 128.39±0.28 | 13.9±0.28 | 0.05±0.003 | 0.507±0.002 | 0.443±0.003 | 0.902±0.002 | 0.812±0.005 |
| | | 200 | 142.46±0 | 120.88±0.53 | 21.58±0.53 | 0.068±0.003 | 0.515±0.007 | 0.417±0.005 | 0.849±0.004 | 0.705±0.009 |

| | | | | | | | | | | |
|---|---|---|---|---|---|---|---|---|---|---|
| | | 400 | 142.55±0 | 108.93±0.46 | 33.63±0.46 | 0.099±0.004 | 0.53±0.004 | 0.371±0.002 | 0.764±0.003 | 0.537±0.009 |
| | chr8 | 2.5 | 148.86±0 | 145.22±0.01 | 3.64±0.01 | 0.02±0 | 0.417±0.001 | 0.562±0.001 | 0.976±0 | 0.952±0 |
| | | 5 | 155.52±0 | 151.5±0.02 | 4.02±0.02 | 0.021±0 | 0.406±0.002 | 0.573±0.003 | 0.974±0 | 0.949±0 |
| | | 10 | 159.22±0 | 154.77±0.01 | 4.45±0.01 | 0.02±0.001 | 0.401±0.002 | 0.579±0.002 | 0.972±0 | 0.945±0 |
| | | 20 | 161.2±0 | 155.97±0.06 | 5.23±0.06 | 0.022±0 | 0.416±0.003 | 0.562±0.004 | 0.968±0 | 0.936±0.001 |
| | | 40 | 162.21±0 | 155.26±0.19 | 6.95±0.19 | 0.027±0.002 | 0.452±0.006 | 0.521±0.008 | 0.957±0.001 | 0.915±0.003 |
| | | 50 | 162.42±0 | 154.75±0.24 | 7.67±0.24 | 0.03±0.002 | 0.46±0.006 | 0.51±0.007 | 0.953±0.001 | 0.906±0.003 |
| | | 60 | 162.56±0 | 153.84±0.16 | 8.72±0.16 | 0.033±0.001 | 0.475±0.003 | 0.492±0.002 | 0.946±0.001 | 0.893±0.002 |
| | | 80 | 162.73±0 | 152.5±0.15 | 10.23±0.15 | 0.036±0.001 | 0.482±0.011 | 0.482±0.011 | 0.937±0.001 | 0.874±0.002 |
| | | 100 | 162.83±0 | 151.08±0.19 | 11.75±0.19 | 0.04±0.002 | 0.494±0.004 | 0.465±0.005 | 0.928±0.001 | 0.855±0.003 |
| | | 200 | 163.04±0 | 144.18±0.45 | 18.86±0.45 | 0.055±0.002 | 0.512±0.003 | 0.433±0.004 | 0.884±0.003 | 0.765±0.006 |
| | | 400 | 163.15±0 | 132.52±0.52 | 30.62±0.52 | 0.083±0.004 | 0.539±0.006 | 0.378±0.003 | 0.812±0.003 | 0.613±0.008 |
| | chr9 | 2.5 | 64.51±0 | 62.93±0.01 | 1.58±0.01 | 0.024±0.001 | 0.43±0.001 | 0.547±0.002 | 0.975±0 | 0.959±0 |
| | | 5 | 67.3±0 | 65.51±0.01 | 1.79±0.01 | 0.024±0 | 0.424±0.002 | 0.552±0.002 | 0.973±0 | 0.955±0 |
| | | 10 | 68.95±0 | 66.91±0.02 | 2.04±0.02 | 0.027±0.001 | 0.423±0.003 | 0.55±0.003 | 0.97±0 | 0.95±0.001 |
| | | 20 | 69.88±0 | 67.43±0.06 | 2.45±0.06 | 0.031±0.004 | 0.433±0.004 | 0.535±0.005 | 0.965±0.001 | 0.94±0.002 |
| | | 40 | 70.37±0 | 67.04±0.07 | 3.32±0.07 | 0.039±0.003 | 0.462±0.003 | 0.5±0.003 | 0.953±0.001 | 0.918±0.002 |
| | | 50 | 70.47±0 | 66.82±0.12 | 3.64±0.12 | 0.04±0.006 | 0.474±0.005 | 0.487±0.007 | 0.948±0.002 | 0.91±0.004 |
| | | 60 | 70.53±0 | 66.33±0.15 | 4.2±0.15 | 0.047±0.008 | 0.476±0.008 | 0.477±0.009 | 0.94±0.002 | 0.895±0.006 |
| | | 80 | 70.62±0 | 65.62±0.26 | 4.99±0.26 | 0.051±0.003 | 0.487±0.007 | 0.462±0.004 | 0.929±0.004 | 0.875±0.007 |
| | | 100 | 70.67±0 | 65.06±0.12 | 5.61±0.12 | 0.052±0.004 | 0.487±0.005 | 0.46±0.009 | 0.921±0.002 | 0.86±0.004 |
| | | 200 | 70.77±0 | 61.52±0.3 | 9.25±0.3 | 0.065±0.005 | 0.499±0.015 | 0.436±0.015 | 0.869±0.004 | 0.768±0.009 |
| | | 400 | 70.82±0 | 56.3±0.56 | 14.52±0.56 | 0.084±0.007 | 0.51±0.007 | 0.406±0.005 | 0.795±0.008 | 0.638±0.017 |
| | chr10 | 2.5 | 112.86±0 | 110.06±0.01 | 2.8±0.01 | 0.022±0.001 | 0.434±0.001 | 0.544±0.001 | 0.975±0 | 0.954±0 |
| | | 5 | 116.41±0 | 113.25±0.01 | 3.17±0.01 | 0.022±0.001 | 0.428±0.002 | 0.55±0.003 | 0.973±0 | 0.949±0 |
| | | 10 | 118.33±0 | 114.57±0.03 | 3.76±0.03 | 0.024±0.001 | 0.433±0.002 | 0.543±0.002 | 0.968±0 | 0.94±0.001 |
| | | 20 | 119.33±0 | 114.49±0.11 | 4.84±0.11 | 0.028±0.001 | 0.448±0.004 | 0.524±0.004 | 0.959±0.001 | 0.923±0.002 |
| | | 40 | 119.84±0 | 112.76±0.06 | 7.09±0.06 | 0.034±0.002 | 0.479±0.003 | 0.487±0.004 | 0.941±0.001 | 0.887±0.001 |
| | | 50 | 119.95±0 | 111.42±0.14 | 8.52±0.14 | 0.04±0.002 | 0.489±0.007 | 0.471±0.006 | 0.929±0.001 | 0.863±0.003 |
| | | 60 | 120.02±0 | 110.33±0.41 | 9.68±0.41 | 0.044±0.003 | 0.493±0.006 | 0.462±0.009 | 0.919±0.003 | 0.844±0.007 |
| | | 80 | 120.1±0 | 108.36±0.47 | 11.74±0.47 | 0.047±0.003 | 0.502±0.002 | 0.45±0.004 | 0.902±0.004 | 0.811±0.008 |
| | | 100 | 120.15±0 | 106.46±0.45 | 13.69±0.45 | 0.051±0.003 | 0.508±0.003 | 0.441±0.003 | 0.886±0.004 | 0.78±0.008 |
| | | 200 | 120.26±0 | 97.75±0.49 | 22.51±0.49 | 0.075±0.005 | 0.521±0.007 | 0.404±0.007 | 0.813±0.004 | 0.635±0.009 |
| | | 400 | 120.31±0 | 86.78±0.27 | 33.53±0.27 | 0.101±0.003 | 0.529±0.003 | 0.369±0.005 | 0.721±0.002 | 0.463±0.004 |
| | chr11 | 2.5 | 93.82±0 | 91.6±0 | 2.22±0 | 0.02±0.001 | 0.428±0.001 | 0.552±0.001 | 0.976±0 | 0.957±0 |

| | | | | | | | | | | |
|---|---|---|---|---|---|---|---|---|---|---|
| | | 5 | 97.61±0 | 95.07±0.02 | 2.54±0.02 | 0.021±0.001 | 0.422±0.003 | 0.558±0.003 | 0.974±0 | 0.953±0 |
| | | 10 | 99.73±0 | 96.78±0.02 | 2.95±0.02 | 0.021±0 | 0.423±0.002 | 0.556±0.002 | 0.97±0 | 0.946±0 |
| | | 20 | 100.86±0 | 97.07±0.03 | 3.79±0.03 | 0.024±0.001 | 0.444±0.001 | 0.532±0.002 | 0.962±0 | 0.931±0.001 |
| | | 40 | 101.44±0 | 95.95±0.16 | 5.49±0.16 | 0.031±0.002 | 0.469±0.004 | 0.5±0.005 | 0.946±0.002 | 0.9±0.003 |
| | | 50 | 101.55±0 | 95.2±0.17 | 6.35±0.17 | 0.036±0.003 | 0.478±0.002 | 0.486±0.004 | 0.937±0.002 | 0.883±0.004 |
| | | 60 | 101.63±0 | 94.42±0.22 | 7.21±0.22 | 0.039±0.003 | 0.486±0.003 | 0.475±0.005 | 0.929±0.002 | 0.867±0.005 |
| | | 80 | 101.73±0 | 92.83±0.21 | 8.9±0.21 | 0.044±0.003 | 0.492±0.004 | 0.464±0.006 | 0.912±0.002 | 0.836±0.005 |
| | | 100 | 101.79±0 | 91.4±0.39 | 10.39±0.39 | 0.048±0.003 | 0.497±0.005 | 0.455±0.005 | 0.898±0.004 | 0.808±0.008 |
| | | 200 | 101.91±0 | 85.99±0.33 | 15.92±0.33 | 0.068±0.004 | 0.503±0.005 | 0.429±0.006 | 0.844±0.003 | 0.701±0.006 |
| | | 400 | 101.97±0 | 77.64±0.56 | 24.32±0.56 | 0.095±0.003 | 0.524±0.001 | 0.382±0.004 | 0.761±0.005 | 0.544±0.011 |
| | chr12 | 2.5 | 76.27±0 | 74.31±0.01 | 1.96±0.01 | 0.02±0 | 0.425±0.002 | 0.554±0.002 | 0.974±0 | 0.953±0 |
| | | 5 | 79.16±0 | 76.89±0.01 | 2.28±0.01 | 0.021±0.001 | 0.423±0.002 | 0.555±0.003 | 0.971±0 | 0.947±0 |
| | | 10 | 80.78±0 | 78.05±0.02 | 2.73±0.02 | 0.024±0.001 | 0.434±0.005 | 0.542±0.005 | 0.966±0 | 0.938±0.001 |
| | | 20 | 81.64±0 | 78.03±0.14 | 3.61±0.14 | 0.031±0.004 | 0.459±0.008 | 0.51±0.011 | 0.956±0.002 | 0.917±0.004 |
| | | 40 | 82.08±0 | 76.69±0.09 | 5.39±0.09 | 0.039±0.002 | 0.482±0.002 | 0.479±0.004 | 0.934±0.001 | 0.876±0.002 |
| | | 50 | 82.17±0 | 76.1±0.22 | 6.08±0.22 | 0.043±0.005 | 0.485±0.006 | 0.472±0.004 | 0.926±0.003 | 0.859±0.007 |
| | | 60 | 82.23±0 | 75.31±0.17 | 6.92±0.17 | 0.046±0.004 | 0.499±0.01 | 0.455±0.011 | 0.916±0.002 | 0.84±0.005 |
| | | 80 | 82.31±0 | 73.91±0.25 | 8.4±0.25 | 0.051±0.004 | 0.5±0.006 | 0.449±0.007 | 0.898±0.003 | 0.806±0.006 |
| | | 100 | 82.35±0 | 72.61±0.32 | 9.74±0.32 | 0.055±0.003 | 0.505±0.01 | 0.441±0.01 | 0.882±0.004 | 0.774±0.008 |
| | | 200 | 82.44±0 | 67.27±0.75 | 15.17±0.75 | 0.072±0.004 | 0.515±0.012 | 0.413±0.009 | 0.816±0.009 | 0.649±0.019 |
| | | 400 | 82.49±0 | 60.6±0.91 | 21.9±0.91 | 0.098±0.003 | 0.519±0.01 | 0.383±0.008 | 0.735±0.011 | 0.494±0.021 |
| | chr13 | 2.5 | 159.41±0 | 155.55±0.01 | 3.86±0.01 | 0.017±0 | 0.432±0.001 | 0.551±0.001 | 0.976±0 | 0.955±0 |
| | | 5 | 144.16±53.6 | 140.4±52.31 | 3.76±1.37 | 0.017±0.002 | 0.424±0.001 | 0.559±0.002 | 0.974±0.001 | 0.952±0 |
| | | 10 | 173.23±0 | 168.25±0.02 | 4.98±0.02 | 0.018±0 | 0.423±0.001 | 0.559±0.001 | 0.971±0 | 0.947±0 |
| | | 20 | 176.04±0 | 170.12±0.07 | 5.92±0.07 | 0.02±0 | 0.431±0.005 | 0.549±0.005 | 0.966±0 | 0.938±0.001 |
| | | 40 | 177.51±0 | 169.87±0.1 | 7.64±0.1 | 0.023±0.001 | 0.456±0.004 | 0.521±0.004 | 0.957±0.001 | 0.92±0.001 |
| | | 50 | 177.81±0 | 169.36±0.18 | 8.46±0.18 | 0.026±0.001 | 0.463±0.001 | 0.511±0.002 | 0.952±0.001 | 0.911±0.002 |
| | | 60 | 178.02±0 | 168.76±0.18 | 9.26±0.18 | 0.03±0.001 | 0.468±0.003 | 0.502±0.003 | 0.948±0.001 | 0.902±0.002 |
| | | 80 | 178.27±0 | 167.24±0.3 | 11.03±0.3 | 0.035±0.001 | 0.476±0.002 | 0.489±0.003 | 0.938±0.002 | 0.882±0.003 |
| | | 100 | 178.42±0 | 165.98±0.26 | 12.44±0.26 | 0.037±0.002 | 0.486±0.006 | 0.478±0.008 | 0.93±0.001 | 0.867±0.003 |
| | | 200 | 178.73±0 | 159.04±0.31 | 19.69±0.31 | 0.056±0.002 | 0.504±0.007 | 0.44±0.008 | 0.89±0.002 | 0.785±0.004 |
| | | 400 | 178.89±0 | 148.7±1.17 | 30.19±1.17 | 0.083±0.004 | 0.527±0.003 | 0.39±0.005 | 0.831±0.007 | 0.663±0.014 |
| | chr14 | 2.5 | 141.3±0 | 137.98±0.01 | 3.32±0.01 | 0.02±0 | 0.428±0.002 | 0.552±0.002 | 0.977±0 | 0.954±0 |
| | | 5 | 147.95±0 | 144.18±0.02 | 3.77±0.02 | 0.02±0 | 0.423±0.002 | 0.557±0.002 | 0.975±0 | 0.95±0 |
| | | 10 | 151.64±0 | 147.26±0.06 | 4.38±0.06 | 0.022±0.001 | 0.428±0.004 | 0.55±0.004 | 0.971±0 | 0.943±0.001 |

| | | | | | | | | | | |
|---|---|---|---|---|---|---|---|---|---|---|
| | | 20 | 153.59±0 | 148.07±0.07 | 5.52±0.07 | 0.028±0 | 0.449±0.003 | 0.523±0.003 | 0.964±0 | 0.929±0.001 |
| | | 40 | 154.59±0 | 146.93±0.07 | 7.66±0.07 | 0.036±0.002 | 0.475±0.002 | 0.489±0.003 | 0.95±0 | 0.9±0.001 |
| | | 50 | 154.8±0 | 146.12±0.13 | 8.68±0.13 | 0.038±0.001 | 0.484±0.006 | 0.478±0.006 | 0.944±0.001 | 0.887±0.002 |
| | | 60 | 154.93±0 | 145.24±0.22 | 9.69±0.22 | 0.043±0.002 | 0.482±0.005 | 0.476±0.003 | 0.937±0.001 | 0.872±0.003 |
| | | 80 | 155.11±0 | 143.52±0.34 | 11.58±0.34 | 0.048±0.004 | 0.489±0.005 | 0.463±0.006 | 0.925±0.002 | 0.846±0.006 |
| | | 100 | 155.21±0 | 141.8±0.19 | 13.41±0.19 | 0.052±0.002 | 0.501±0.005 | 0.448±0.005 | 0.914±0.001 | 0.822±0.003 |
| | | 200 | 155.42±0 | 134.87±0.3 | 20.55±0.3 | 0.07±0.002 | 0.518±0.004 | 0.413±0.003 | 0.868±0.002 | 0.724±0.005 |
| | | 400 | 155.52±0 | 123.84±0.67 | 31.69±0.67 | 0.104±0.002 | 0.532±0.008 | 0.364±0.009 | 0.796±0.004 | 0.565±0.008 |
| | chr15 | 2.5 | 133.47±0 | 130.26±0 | 3.22±0 | 0.019±0 | 0.444±0.002 | 0.537±0.002 | 0.976±0 | 0.956±0 |
| | | 5 | 139.55±0 | 135.96±0.02 | 3.58±0.02 | 0.019±0 | 0.433±0.004 | 0.549±0.004 | 0.974±0 | 0.954±0 |
| | | 10 | 142.88±0 | 138.93±0.02 | 3.96±0.02 | 0.019±0 | 0.428±0.002 | 0.554±0.002 | 0.972±0 | 0.95±0 |
| | | 20 | 144.65±0 | 139.96±0.02 | 4.69±0.02 | 0.021±0.001 | 0.438±0.003 | 0.541±0.003 | 0.968±0 | 0.941±0 |
| | | 40 | 145.57±0 | 139.38±0.12 | 6.19±0.12 | 0.027±0.001 | 0.46±0.003 | 0.514±0.004 | 0.957±0.001 | 0.922±0.002 |
| | | 50 | 145.76±0 | 138.72±0.13 | 7.03±0.13 | 0.029±0.002 | 0.472±0.002 | 0.498±0.002 | 0.952±0.001 | 0.911±0.002 |
| | | 60 | 145.88±0 | 138.05±0.17 | 7.84±0.17 | 0.032±0.002 | 0.476±0.005 | 0.492±0.005 | 0.946±0.001 | 0.901±0.002 |
| | | 80 | 146.04±0 | 136.68±0.1 | 9.36±0.1 | 0.038±0.002 | 0.485±0.004 | 0.476±0.002 | 0.936±0.001 | 0.88±0.002 |
| | | 100 | 146.14±0 | 135.26±0.39 | 10.88±0.39 | 0.041±0.001 | 0.493±0.003 | 0.466±0.003 | 0.926±0.003 | 0.861±0.005 |
| | | 200 | 146.33±0 | 128.31±0.78 | 18.02±0.78 | 0.06±0.006 | 0.513±0.004 | 0.428±0.006 | 0.877±0.005 | 0.766±0.013 |
| | | 400 | 146.43±0 | 117.9±0.85 | 28.53±0.85 | 0.086±0.004 | 0.53±0.009 | 0.384±0.008 | 0.805±0.006 | 0.626±0.012 |
| | chr16 | 2.5 | 88.73±0 | 86.43±0.01 | 2.3±0.01 | 0.023±0.001 | 0.425±0.001 | 0.552±0.001 | 0.974±0 | 0.955±0 |
| | | 5 | 92.33±0 | 89.74±0.02 | 2.59±0.02 | 0.023±0.001 | 0.417±0.004 | 0.56±0.004 | 0.972±0 | 0.951±0 |
| | | 10 | 94.37±0 | 91.41±0.02 | 2.97±0.02 | 0.025±0 | 0.42±0.003 | 0.555±0.003 | 0.969±0 | 0.945±0 |
| | | 20 | 95.47±0 | 91.78±0.06 | 3.69±0.06 | 0.032±0.001 | 0.434±0.005 | 0.534±0.005 | 0.961±0.001 | 0.931±0.001 |
| | | 40 | 96.05±0 | 90.79±0.21 | 5.25±0.21 | 0.038±0.003 | 0.46±0.004 | 0.503±0.006 | 0.945±0.002 | 0.902±0.004 |
| | | 50 | 96.16±0 | 90.05±0.09 | 6.11±0.09 | 0.044±0.003 | 0.468±0.007 | 0.488±0.009 | 0.936±0.001 | 0.885±0.002 |
| | | 60 | 96.24±0 | 89.61±0.24 | 6.63±0.24 | 0.045±0.003 | 0.475±0.002 | 0.48±0.003 | 0.931±0.003 | 0.875±0.005 |
| | | 80 | 96.34±0 | 88.01±0.22 | 8.33±0.22 | 0.053±0.005 | 0.478±0.004 | 0.469±0.005 | 0.913±0.002 | 0.841±0.006 |
| | | 100 | 96.4±0 | 86.6±0.36 | 9.8±0.36 | 0.061±0.004 | 0.481±0.004 | 0.458±0.004 | 0.898±0.004 | 0.811±0.007 |
| | | 200 | 96.52±0 | 80.25±0.25 | 16.27±0.25 | 0.089±0.005 | 0.491±0.007 | 0.42±0.007 | 0.831±0.003 | 0.68±0.008 |
| | | 400 | 96.58±0 | 72.97±0.31 | 23.61±0.31 | 0.112±0.002 | 0.502±0.005 | 0.386±0.005 | 0.756±0.003 | 0.539±0.007 |
| | chr17 | 2.5 | 83.78±0 | 81.56±0.01 | 2.23±0.01 | 0.024±0.001 | 0.448±0.001 | 0.528±0.001 | 0.973±0 | 0.951±0 |
| | | 5 | 86.78±0 | 84.23±0.01 | 2.55±0.01 | 0.025±0.001 | 0.445±0.003 | 0.53±0.003 | 0.971±0 | 0.946±0 |
| | | 10 | 88.43±0 | 85.46±0.05 | 2.97±0.05 | 0.027±0.001 | 0.453±0.003 | 0.52±0.004 | 0.966±0.001 | 0.938±0.001 |
| | | 20 | 89.31±0 | 85.61±0.08 | 3.7±0.08 | 0.032±0.002 | 0.469±0.004 | 0.5±0.005 | 0.959±0.001 | 0.922±0.002 |
| | | 40 | 89.77±0 | 84.75±0.09 | 5.02±0.09 | 0.037±0.003 | 0.492±0.005 | 0.471±0.006 | 0.944±0.001 | 0.895±0.003 |

| | | | | | | | | | | |
|---|---|---|---|---|---|---|---|---|---|---|
| | | 50 | 89.86±0 | 84.13±0.22 | 5.74±0.22 | 0.039±0.003 | 0.504±0.007 | 0.457±0.006 | 0.936±0.002 | 0.88±0.005 |
| | | 60 | 89.93±0 | 83.66±0.18 | 6.27±0.18 | 0.04±0.003 | 0.509±0.003 | 0.451±0.004 | 0.93±0.002 | 0.869±0.004 |
| | | 80 | 90±0 | 82.36±0.18 | 7.64±0.18 | 0.045±0.002 | 0.514±0.006 | 0.441±0.008 | 0.915±0.002 | 0.839±0.004 |
| | | 100 | 90.05±0 | 81.15±0.32 | 8.9±0.32 | 0.049±0.003 | 0.522±0.01 | 0.429±0.008 | 0.901±0.004 | 0.813±0.007 |
| | | 200 | 90.14±0 | 75.96±0.23 | 14.18±0.23 | 0.063±0.002 | 0.53±0.01 | 0.407±0.01 | 0.843±0.003 | 0.702±0.005 |
| | | 400 | 90.19±0 | 68.67±0.31 | 21.52±0.31 | 0.086±0.004 | 0.534±0.005 | 0.38±0.006 | 0.761±0.003 | 0.549±0.008 |
| | chr18 | 2.5 | 60.34±0 | 58.83±0.01 | 1.51±0.01 | 0.023±0.001 | 0.438±0.005 | 0.54±0.005 | 0.975±0 | 0.956±0 |
| | | 5 | 62.8±0 | 61.09±0.01 | 1.71±0.01 | 0.024±0.001 | 0.429±0.004 | 0.547±0.005 | 0.973±0 | 0.952±0 |
| | | 10 | 64.21±0 | 62.28±0.01 | 1.93±0.01 | 0.025±0.001 | 0.429±0.003 | 0.546±0.003 | 0.97±0 | 0.947±0 |
| | | 20 | 64.98±0 | 62.7±0.05 | 2.27±0.05 | 0.027±0.002 | 0.434±0.006 | 0.539±0.004 | 0.965±0.001 | 0.938±0.002 |
| | | 40 | 65.38±0 | 62.41±0.04 | 2.97±0.04 | 0.034±0.002 | 0.46±0.01 | 0.506±0.009 | 0.955±0.001 | 0.918±0.001 |
| | | 50 | 65.46±0 | 62.06±0.09 | 3.4±0.09 | 0.036±0.002 | 0.473±0.001 | 0.492±0.004 | 0.948±0.001 | 0.906±0.003 |
| | | 60 | 65.52±0 | 61.67±0.12 | 3.85±0.12 | 0.039±0.004 | 0.477±0.014 | 0.484±0.017 | 0.941±0.002 | 0.893±0.004 |
| | | 80 | 65.59±0 | 60.98±0.18 | 4.61±0.18 | 0.042±0.001 | 0.491±0.005 | 0.467±0.005 | 0.93±0.003 | 0.872±0.005 |
| | | 100 | 65.63±0 | 60.28±0.19 | 5.35±0.19 | 0.048±0.003 | 0.483±0.006 | 0.469±0.005 | 0.918±0.003 | 0.85±0.006 |
| | | 200 | 65.71±0 | 56.46±0.44 | 9.25±0.44 | 0.066±0.004 | 0.484±0.007 | 0.449±0.004 | 0.859±0.007 | 0.738±0.014 |
| | | 400 | 65.75±0 | 51.41±0.79 | 14.35±0.79 | 0.09±0.011 | 0.494±0.011 | 0.416±0.011 | 0.782±0.012 | 0.592±0.028 |
| Test dataset 3 | chr1 | 2.5 | 32.72±0 | 27.94±0.01 | 4.78±0.01 | 0.135±0.001 | 0.296±0.001 | 0.569±0.001 | 0.854±0 | 0.73±0.001 |
| | | 5 | 34.34±0 | 28.75±0.02 | 5.59±0.02 | 0.144±0.001 | 0.32±0.002 | 0.536±0.002 | 0.837±0 | 0.697±0.001 |
| | | 10 | 35.25±0 | 28.7±0.02 | 6.55±0.02 | 0.156±0 | 0.34±0.001 | 0.504±0.001 | 0.814±0.001 | 0.65±0.001 |
| | | 20 | 35.76±0 | 28.01±0.04 | 7.75±0.04 | 0.173±0.002 | 0.355±0.003 | 0.472±0.004 | 0.783±0.001 | 0.586±0.003 |
| | | 40 | 36.05±0 | 26.95±0.04 | 9.1±0.04 | 0.198±0.001 | 0.359±0.003 | 0.444±0.004 | 0.748±0.001 | 0.509±0.002 |
| | | 50 | 36.11±0 | 26.51±0.1 | 9.61±0.1 | 0.205±0.002 | 0.363±0.002 | 0.432±0.003 | 0.734±0.003 | 0.481±0.005 |
| | | 60 | 36.16±0 | 26.09±0.04 | 10.07±0.04 | 0.214±0.003 | 0.361±0.002 | 0.425±0.004 | 0.722±0.001 | 0.454±0.003 |
| | | 80 | 36.22±0 | 25.48±0.01 | 10.73±0.01 | 0.225±0.004 | 0.363±0.001 | 0.412±0.004 | 0.704±0 | 0.417±0.003 |
| | | 100 | 36.25±0 | 24.97±0.09 | 11.28±0.09 | 0.234±0.002 | 0.366±0.002 | 0.401±0.004 | 0.689±0.003 | 0.387±0.005 |
| | | 200 | 36.32±0 | 23.26±0.03 | 13.07±0.03 | 0.266±0.003 | 0.365±0.003 | 0.369±0.006 | 0.64±0.001 | 0.289±0.002 |
| | chr2 | 2.5 | 22.88±0 | 19.07±0.01 | 3.82±0.01 | 0.132±0.001 | 0.327±0.002 | 0.541±0.003 | 0.833±0.001 | 0.681±0.001 |
| | | 5 | 23.85±0 | 19.31±0.03 | 4.54±0.03 | 0.145±0.001 | 0.351±0.002 | 0.504±0.003 | 0.809±0.001 | 0.631±0.003 |
| | | 10 | 24.39±0 | 18.92±0.04 | 5.47±0.04 | 0.164±0.001 | 0.37±0.002 | 0.466±0.002 | 0.776±0.002 | 0.561±0.003 |
| | | 20 | 24.69±0 | 18.11±0.02 | 6.57±0.02 | 0.187±0.001 | 0.38±0.001 | 0.433±0.002 | 0.734±0.001 | 0.473±0.002 |
| | | 40 | 24.85±0 | 16.99±0.09 | 7.86±0.09 | 0.213±0.002 | 0.384±0.004 | 0.402±0.005 | 0.684±0.004 | 0.372±0.007 |
| | | 50 | 24.89±0 | 16.55±0.05 | 8.33±0.05 | 0.222±0.002 | 0.386±0.002 | 0.392±0.004 | 0.665±0.002 | 0.337±0.005 |
| | | 60 | 24.91±0 | 16.33±0.06 | 8.58±0.06 | 0.227±0.003 | 0.39±0.003 | 0.383±0.003 | 0.656±0.002 | 0.32±0.006 |
| | | 80 | 24.94±0 | 15.87±0.08 | 9.07±0.08 | 0.236±0.002 | 0.389±0.002 | 0.375±0.003 | 0.636±0.003 | 0.284±0.006 |

| | | | | | | | | | | |
|---|---|---|---|---|---|---|---|---|---|---|
| | | 100 | 24.96±0 | 15.58±0.08 | 9.38±0.08 | 0.244±0.003 | 0.392±0.002 | 0.365±0.004 | 0.624±0.003 | 0.261±0.003 |
| | | 200 | 25±0 | 14.7±0.11 | 10.3±0.11 | 0.268±0.006 | 0.398±0.003 | 0.334±0.006 | 0.588±0.005 | 0.196±0.011 |
| | chr3 | 2.5 | 24.47±10.3 | 21.2±8.88 | 3.27±1.42 | 0.115±0.002 | 0.277±0.004 | 0.608±0.002 | 0.868±0.006 | 0.754±0.008 |
| | | 5 | 25.31±10.69 | 21.44±9.04 | 3.87±1.65 | 0.128±0.005 | 0.313±0.006 | 0.559±0.011 | 0.848±0.002 | 0.711±0.004 |
| | | 10 | 30.64±0 | 25.09±0.03 | 5.55±0.03 | 0.142±0.002 | 0.342±0.001 | 0.516±0.001 | 0.819±0.001 | 0.655±0.002 |
| | | 20 | 30.94±0 | 24.19±0.03 | 6.75±0.03 | 0.16±0.001 | 0.367±0.003 | 0.473±0.005 | 0.782±0.001 | 0.58±0.002 |
| | | 40 | 31.1±0 | 22.89±0.04 | 8.21±0.04 | 0.185±0.002 | 0.379±0.003 | 0.436±0.002 | 0.736±0.001 | 0.486±0.004 |
| | | 50 | 31.13±0 | 22.47±0.11 | 8.66±0.11 | 0.193±0.004 | 0.382±0.004 | 0.425±0.003 | 0.722±0.003 | 0.456±0.009 |
| | | 60 | 31.15±0 | 22.07±0.07 | 9.09±0.07 | 0.199±0.003 | 0.388±0.003 | 0.413±0.002 | 0.708±0.002 | 0.431±0.006 |
| | | 80 | 31.18±0 | 21.42±0.12 | 9.76±0.12 | 0.21±0.003 | 0.391±0.003 | 0.399±0.003 | 0.687±0.004 | 0.389±0.009 |
| | | 100 | 31.2±0 | 20.93±0.17 | 10.27±0.17 | 0.218±0.006 | 0.391±0.002 | 0.39±0.006 | 0.671±0.006 | 0.358±0.013 |
| | | 200 | 31.23±0 | 19.38±0.03 | 11.85±0.03 | 0.247±0.007 | 0.396±0.002 | 0.358±0.008 | 0.62±0.001 | 0.264±0.005 |
| | chr4 | 2.5 | 25.6±0 | 21.41±0.01 | 4.19±0.01 | 0.128±0.001 | 0.294±0.001 | 0.579±0.002 | 0.836±0 | 0.687±0.001 |
| | | 5 | 23.75±6.3 | 19.37±5.16 | 4.38±1.14 | 0.142±0.003 | 0.315±0.002 | 0.542±0.003 | 0.815±0.002 | 0.64±0.006 |
| | | 10 | 27.09±0 | 21.36±0.05 | 5.73±0.05 | 0.158±0.003 | 0.334±0.001 | 0.507±0.003 | 0.788±0.002 | 0.583±0.006 |
| | | 20 | 27.37±0 | 20.54±0.05 | 6.83±0.05 | 0.179±0.002 | 0.349±0.001 | 0.471±0.002 | 0.75±0.002 | 0.501±0.004 |
| | | 40 | 27.53±0 | 19.46±0.03 | 8.07±0.03 | 0.199±0.004 | 0.36±0.004 | 0.441±0.006 | 0.707±0.001 | 0.414±0.004 |
| | | 50 | 27.56±0 | 19.05±0.05 | 8.51±0.05 | 0.207±0.003 | 0.361±0.001 | 0.432±0.004 | 0.691±0.002 | 0.381±0.005 |
| | | 60 | 27.58±0 | 18.73±0.07 | 8.85±0.07 | 0.213±0.004 | 0.364±0.004 | 0.423±0.005 | 0.679±0.002 | 0.357±0.007 |
| | | 80 | 27.61±0 | 18.18±0.05 | 9.44±0.05 | 0.222±0.005 | 0.364±0.003 | 0.414±0.006 | 0.658±0.002 | 0.317±0.006 |
| | | 100 | 27.63±0 | 17.79±0.06 | 9.84±0.06 | 0.226±0.004 | 0.368±0.003 | 0.406±0.004 | 0.644±0.002 | 0.293±0.006 |
| | | 200 | 27.66±0 | 16.63±0.11 | 11.03±0.11 | 0.249±0.004 | 0.375±0.002 | 0.376±0.005 | 0.601±0.004 | 0.216±0.008 |
| | chr5 | 2.5 | 19.68±0 | 16.54±0.01 | 3.13±0.01 | 0.138±0.002 | 0.299±0.002 | 0.563±0.003 | 0.841±0.001 | 0.693±0.002 |
| | | 5 | 17.5±6.56 | 14.29±5.32 | 3.2±1.24 | 0.159±0.008 | 0.329±0.003 | 0.512±0.011 | 0.818±0.005 | 0.641±0.006 |
| | | 10 | 20.84±0 | 16.31±0.02 | 4.53±0.02 | 0.175±0.002 | 0.351±0.003 | 0.474±0.004 | 0.782±0.001 | 0.567±0.002 |
| | | 20 | 21.07±0 | 15.53±0.04 | 5.54±0.04 | 0.201±0.002 | 0.361±0.001 | 0.438±0.003 | 0.737±0.002 | 0.47±0.005 |
| | | 40 | 21.19±0 | 14.52±0.05 | 6.67±0.05 | 0.228±0.004 | 0.368±0.003 | 0.404±0.003 | 0.685±0.002 | 0.364±0.007 |
| | | 50 | 21.22±0 | 14.24±0.03 | 6.98±0.03 | 0.233±0.003 | 0.37±0.002 | 0.397±0.002 | 0.671±0.001 | 0.338±0.002 |
| | | 60 | 21.23±0 | 13.93±0.06 | 7.3±0.06 | 0.239±0.002 | 0.367±0.003 | 0.393±0.004 | 0.656±0.003 | 0.31±0.006 |
| | | 80 | 21.26±0 | 13.48±0.09 | 7.77±0.09 | 0.25±0.001 | 0.369±0.003 | 0.381±0.002 | 0.634±0.004 | 0.27±0.007 |
| | | 100 | 21.27±0 | 13.1±0.03 | 8.17±0.03 | 0.259±0.005 | 0.37±0.004 | 0.371±0.009 | 0.616±0.001 | 0.236±0.005 |
| | | 200 | 21.3±0 | 12.22±0.1 | 9.08±0.1 | 0.28±0.006 | 0.372±0.003 | 0.348±0.007 | 0.574±0.005 | 0.166±0.008 |
| | chr6 | 2.5 | 31.75±0 | 27.01±0.01 | 4.74±0.01 | 0.122±0.001 | 0.29±0.002 | 0.588±0.002 | 0.851±0 | 0.718±0.001 |
| | | 5 | 33.04±0 | 27.47±0.02 | 5.57±0.02 | 0.135±0.001 | 0.316±0.002 | 0.549±0.002 | 0.831±0.001 | 0.677±0.001 |
| | | 10 | 33.75±0 | 27.08±0.01 | 6.67±0.01 | 0.157±0.001 | 0.339±0.001 | 0.503±0.001 | 0.802±0 | 0.611±0.001 |

| | | | | | | | | | | |
|---|---|---|---|---|---|---|---|---|---|---|
| | | 20 | 34.13±0 | 26.07±0.04 | 8.06±0.04 | 0.179±0.002 | 0.357±0.003 | 0.464±0.004 | 0.764±0.001 | 0.529±0.004 |
| | | 40 | 34.33±0 | 24.58±0.08 | 9.74±0.08 | 0.206±0.003 | 0.368±0.004 | 0.426±0.006 | 0.716±0.002 | 0.429±0.005 |
| | | 50 | 34.37±0 | 24.09±0.16 | 10.28±0.16 | 0.213±0.003 | 0.372±0.003 | 0.415±0.004 | 0.701±0.005 | 0.399±0.01 |
| | | 60 | 34.4±0 | 23.64±0.09 | 10.76±0.09 | 0.221±0.002 | 0.374±0.003 | 0.405±0.002 | 0.687±0.003 | 0.371±0.006 |
| | | 80 | 34.44±0 | 22.88±0.05 | 11.56±0.05 | 0.233±0.002 | 0.374±0.003 | 0.394±0.003 | 0.664±0.001 | 0.326±0.002 |
| | | 100 | 34.46±0 | 22.3±0.07 | 12.16±0.07 | 0.24±0.002 | 0.376±0.002 | 0.384±0.003 | 0.647±0.002 | 0.295±0.005 |
| | | 200 | 34.5±0 | 20.74±0.09 | 13.77±0.09 | 0.265±0.002 | 0.382±0.003 | 0.353±0.003 | 0.601±0.003 | 0.212±0.005 |
| | chr7 | 2.5 | 26.81±0 | 22.63±0.01 | 4.18±0.01 | 0.12±0.001 | 0.298±0.003 | 0.581±0.003 | 0.844±0 | 0.707±0.001 |
| | | 5 | 27.74±0 | 22.88±0.01 | 4.86±0.01 | 0.132±0.001 | 0.325±0.001 | 0.543±0.002 | 0.825±0 | 0.667±0.002 |
| | | 10 | 28.24±0 | 22.54±0.02 | 5.7±0.02 | 0.147±0.002 | 0.348±0.002 | 0.504±0.004 | 0.798±0.001 | 0.611±0.001 |
| | | 20 | 28.5±0 | 21.78±0.04 | 6.73±0.04 | 0.163±0.002 | 0.369±0.002 | 0.469±0.003 | 0.764±0.001 | 0.542±0.003 |
| | | 40 | 28.65±0 | 20.65±0.04 | 7.99±0.04 | 0.188±0.002 | 0.381±0.002 | 0.431±0.003 | 0.721±0.002 | 0.452±0.004 |
| | | 50 | 28.68±0 | 20.33±0.03 | 8.34±0.03 | 0.19±0.004 | 0.386±0.002 | 0.424±0.002 | 0.709±0.001 | 0.432±0.004 |
| | | 60 | 28.7±0 | 19.94±0.11 | 8.76±0.11 | 0.202±0.004 | 0.385±0.003 | 0.414±0.005 | 0.695±0.004 | 0.4±0.008 |
| | | 80 | 28.72±0 | 19.46±0.11 | 9.26±0.11 | 0.21±0.005 | 0.387±0.002 | 0.403±0.007 | 0.677±0.004 | 0.367±0.009 |
| | | 100 | 28.74±0 | 19.03±0.1 | 9.71±0.1 | 0.219±0.004 | 0.389±0.002 | 0.392±0.004 | 0.662±0.004 | 0.336±0.003 |
| | | 200 | 28.77±0 | 17.81±0.04 | 10.96±0.04 | 0.244±0.004 | 0.396±0.004 | 0.36±0.006 | 0.619±0.001 | 0.256±0.004 |
| | chr8 | 2.5 | 21.64±0 | 17.59±0.01 | 4.05±0.01 | 0.126±0.001 | 0.321±0.001 | 0.552±0.001 | 0.813±0 | 0.633±0.001 |
| | | 5 | 22.44±0 | 17.63±0.01 | 4.8±0.01 | 0.137±0 | 0.347±0.002 | 0.516±0.002 | 0.786±0.001 | 0.578±0.001 |
| | | 10 | 22.88±0 | 17.22±0.03 | 5.66±0.03 | 0.15±0 | 0.364±0.002 | 0.487±0.002 | 0.753±0.001 | 0.511±0.002 |
| | | 20 | 23.13±0 | 16.46±0.03 | 6.67±0.03 | 0.164±0.001 | 0.374±0.003 | 0.462±0.003 | 0.712±0.001 | 0.43±0.002 |
| | | 40 | 23.27±0 | 15.54±0.06 | 7.73±0.06 | 0.181±0.002 | 0.383±0.002 | 0.437±0.001 | 0.668±0.003 | 0.346±0.006 |
| | | 50 | 23.3±0 | 15.19±0.13 | 8.11±0.13 | 0.188±0.002 | 0.385±0.003 | 0.427±0.003 | 0.652±0.006 | 0.317±0.01 |
| | | 60 | 23.32±0 | 14.99±0.04 | 8.33±0.04 | 0.192±0.002 | 0.388±0.001 | 0.42±0.002 | 0.643±0.002 | 0.299±0.003 |
| | | 80 | 23.35±0 | 14.51±0.05 | 8.84±0.05 | 0.202±0.002 | 0.388±0.002 | 0.409±0.004 | 0.622±0.002 | 0.259±0.004 |
| | | 100 | 23.37±0 | 14.25±0.07 | 9.11±0.07 | 0.208±0.003 | 0.392±0.001 | 0.399±0.004 | 0.61±0.003 | 0.239±0.007 |
| | | 200 | 23.4±0 | 13.48±0.02 | 9.92±0.02 | 0.229±0.004 | 0.402±0.002 | 0.368±0.004 | 0.576±0.001 | 0.177±0.004 |
| | chr9 | 2.5 | 15.77±0 | 13.14±0.01 | 2.63±0 | 0.126±0.001 | 0.31±0.001 | 0.564±0.002 | 0.833±0 | 0.683±0.001 |
| | | 5 | 16.3±0 | 13.17±0.01 | 3.13±0.01 | 0.14±0.001 | 0.339±0.002 | 0.52±0.003 | 0.808±0.001 | 0.631±0.002 |
| | | 10 | 16.58±0 | 12.85±0.03 | 3.73±0.03 | 0.158±0.003 | 0.36±0.003 | 0.482±0.005 | 0.775±0.002 | 0.562±0.004 |
| | | 20 | 16.74±0 | 12.28±0.03 | 4.46±0.03 | 0.178±0.001 | 0.376±0.002 | 0.447±0.002 | 0.733±0.002 | 0.478±0.002 |
| | | 40 | 16.82±0 | 11.57±0.09 | 5.25±0.09 | 0.202±0.005 | 0.382±0.002 | 0.416±0.005 | 0.688±0.005 | 0.386±0.013 |
| | | 50 | 16.83±0 | 11.37±0.07 | 5.46±0.07 | 0.208±0.003 | 0.387±0.003 | 0.405±0.007 | 0.675±0.004 | 0.361±0.008 |
| | | 60 | 16.85±0 | 11.19±0.03 | 5.66±0.03 | 0.212±0.004 | 0.387±0.004 | 0.402±0.008 | 0.664±0.002 | 0.341±0.003 |
| | | 80 | 16.86±0 | 10.91±0.07 | 5.95±0.07 | 0.221±0.006 | 0.389±0.004 | 0.39±0.005 | 0.647±0.004 | 0.309±0.009 |

| | | | | | | | | | | |
|---|---|---|---|---|---|---|---|---|---|---|
| | | 100 | 16.87±0 | 10.66±0.05 | 6.21±0.05 | 0.231±0.005 | 0.388±0.002 | 0.381±0.006 | 0.632±0.003 | 0.278±0.007 |
| | | 200 | 16.89±0 | 10.09±0.1 | 6.8±0.1 | 0.247±0.011 | 0.394±0.004 | 0.358±0.014 | 0.597±0.006 | 0.22±0.013 |
| | chcr10 | 2.5 | 18.69±0 | 15.48±0.01 | 3.21±0.01 | 0.137±0.001 | 0.306±0.001 | 0.557±0.002 | 0.828±0.001 | 0.674±0.002 |
| | | 5 | 19.2±0 | 15.42±0.01 | 3.79±0.01 | 0.152±0.001 | 0.33±0.001 | 0.518±0.001 | 0.803±0 | 0.621±0.001 |
| | | 10 | 19.48±0 | 14.89±0.03 | 4.58±0.03 | 0.174±0.002 | 0.349±0.002 | 0.476±0.003 | 0.765±0.002 | 0.54±0.004 |
| | | 20 | 19.62±0 | 14.07±0.05 | 5.55±0.05 | 0.2±0.004 | 0.36±0.003 | 0.439±0.005 | 0.717±0.003 | 0.441±0.006 |
| | | 40 | 19.7±0 | 13.15±0.06 | 6.55±0.06 | 0.228±0.006 | 0.367±0.002 | 0.405±0.007 | 0.668±0.003 | 0.341±0.009 |
| | | 50 | 19.71±0 | 12.81±0.05 | 6.9±0.05 | 0.238±0.005 | 0.366±0.002 | 0.397±0.005 | 0.65±0.003 | 0.306±0.008 |
| | | 60 | 19.72±0 | 12.55±0.05 | 7.17±0.05 | 0.244±0.003 | 0.368±0.003 | 0.388±0.006 | 0.636±0.003 | 0.282±0.005 |
| | | 80 | 19.73±0 | 12.17±0.03 | 7.56±0.03 | 0.253±0.004 | 0.368±0.002 | 0.379±0.005 | 0.617±0.001 | 0.247±0.002 |
| | | 100 | 19.74±0 | 11.91±0.04 | 7.83±0.04 | 0.261±0.005 | 0.371±0.002 | 0.368±0.006 | 0.603±0.002 | 0.223±0.004 |
| | | 200 | 19.76±0 | 11.23±0.04 | 8.53±0.04 | 0.284±0.005 | 0.379±0.002 | 0.337±0.007 | 0.568±0.002 | 0.163±0.006 |
| | chr11 | 2.5 | 18.79±0 | 15.79±0.01 | 3±0.01 | 0.129±0.001 | 0.287±0.002 | 0.584±0.002 | 0.84±0.001 | 0.689±0.001 |
| | | 5 | 19.35±0 | 15.81±0.01 | 3.54±0.01 | 0.145±0.001 | 0.317±0.002 | 0.538±0.003 | 0.817±0 | 0.638±0.001 |
| | | 10 | 19.64±0 | 15.42±0.03 | 4.23±0.03 | 0.165±0.004 | 0.344±0.003 | 0.491±0.005 | 0.785±0.001 | 0.568±0.005 |
| | | 20 | 19.8±0 | 14.74±0.02 | 5.06±0.02 | 0.188±0.002 | 0.364±0.003 | 0.448±0.002 | 0.744±0.001 | 0.481±0.004 |
| | | 40 | 19.89±0 | 13.87±0.12 | 6.02±0.12 | 0.211±0.004 | 0.375±0.003 | 0.414±0.005 | 0.697±0.006 | 0.386±0.012 |
| | | 50 | 19.91±0 | 13.47±0.1 | 6.43±0.1 | 0.22±0.005 | 0.378±0.004 | 0.402±0.004 | 0.677±0.005 | 0.346±0.01 |
| | | 60 | 19.92±0 | 13.24±0.09 | 6.68±0.09 | 0.228±0.006 | 0.38±0.002 | 0.392±0.004 | 0.664±0.005 | 0.321±0.012 |
| | | 80 | 19.94±0 | 12.83±0.11 | 7.11±0.11 | 0.237±0.006 | 0.384±0.005 | 0.379±0.008 | 0.643±0.006 | 0.283±0.013 |
| | | 100 | 19.95±0 | 12.63±0.06 | 7.32±0.06 | 0.244±0.006 | 0.385±0.005 | 0.371±0.009 | 0.633±0.003 | 0.263±0.008 |
| | | 200 | 19.97±0 | 11.7±0.05 | 8.26±0.05 | 0.267±0.003 | 0.397±0.006 | 0.337±0.008 | 0.586±0.002 | 0.185±0.005 |
| | chr12 | 2.5 | 15.74±0 | 13.48±0.01 | 2.26±0.01 | 0.126±0.002 | 0.306±0.001 | 0.568±0.002 | 0.856±0 | 0.725±0.001 |
| | | 5 | 16.27±0 | 13.46±0.02 | 2.81±0.02 | 0.146±0.002 | 0.344±0.003 | 0.51±0.004 | 0.827±0.001 | 0.663±0.003 |
| | | 10 | 16.55±0 | 13.06±0.02 | 3.49±0.02 | 0.166±0.001 | 0.375±0.002 | 0.459±0.003 | 0.789±0.001 | 0.584±0.002 |
| | | 20 | 16.7±0 | 12.32±0.07 | 4.39±0.07 | 0.192±0.002 | 0.392±0.003 | 0.416±0.002 | 0.737±0.004 | 0.48±0.008 |
| | | 40 | 16.78±0 | 11.43±0.09 | 5.35±0.09 | 0.218±0.005 | 0.398±0.004 | 0.384±0.005 | 0.681±0.005 | 0.371±0.011 |
| | | 50 | 16.8±0 | 11.17±0.07 | 5.62±0.07 | 0.22±0.002 | 0.4±0.003 | 0.38±0.003 | 0.665±0.004 | 0.347±0.008 |
| | | 60 | 16.81±0 | 10.85±0.11 | 5.95±0.11 | 0.233±0.007 | 0.405±0.002 | 0.362±0.005 | 0.646±0.007 | 0.309±0.015 |
| | | 80 | 16.82±0 | 10.52±0.07 | 6.31±0.07 | 0.239±0.003 | 0.399±0.002 | 0.362±0.003 | 0.625±0.004 | 0.274±0.007 |
| | | 100 | 16.83±0 | 10.2±0.08 | 6.63±0.08 | 0.251±0.006 | 0.404±0.006 | 0.345±0.008 | 0.606±0.005 | 0.241±0.011 |
| | | 200 | 16.85±0 | 9.31±0.15 | 7.54±0.15 | 0.276±0.005 | 0.398±0.004 | 0.326±0.006 | 0.553±0.009 | 0.157±0.014 |
| | chr13 | 2.5 | 16.25±0 | 13.45±0.01 | 2.8±0.01 | 0.139±0.001 | 0.321±0.002 | 0.54±0.002 | 0.828±0.001 | 0.671±0.001 |
| | | 5 | 17.18±0 | 13.86±0.02 | 3.33±0.02 | 0.153±0 | 0.34±0.002 | 0.507±0.002 | 0.806±0.001 | 0.626±0.002 |
| | | 10 | 17.73±0 | 13.82±0.03 | 3.92±0.03 | 0.169±0.003 | 0.354±0.002 | 0.476±0.003 | 0.779±0.001 | 0.568±0.004 |

| | | | | | | | | | | |
|---|---|---|---|---|---|---|---|---|---|---|
| | | 20 | 18.05±0 | 13.51±0.04 | 4.54±0.04 | 0.186±0.002 | 0.361±0.001 | 0.453±0.003 | 0.749±0.002 | 0.504±0.006 |
| | | 40 | 18.24±0 | 13±0.02 | 5.24±0.02 | 0.206±0.004 | 0.367±0.001 | 0.428±0.004 | 0.713±0.001 | 0.43±0.005 |
| | | 50 | 18.28±0 | 12.78±0.04 | 5.5±0.04 | 0.21±0.002 | 0.368±0.001 | 0.422±0.002 | 0.699±0.002 | 0.405±0.005 |
| | | 60 | 18.31±0 | 12.51±0.04 | 5.8±0.04 | 0.222±0.004 | 0.366±0.004 | 0.412±0.005 | 0.683±0.002 | 0.369±0.007 |
| | | 80 | 18.35±0 | 12.22±0.05 | 6.13±0.05 | 0.233±0.002 | 0.37±0.002 | 0.397±0.003 | 0.666±0.003 | 0.334±0.006 |
| | | 100 | 18.37±0 | 11.99±0.05 | 6.39±0.05 | 0.239±0.005 | 0.371±0.002 | 0.39±0.005 | 0.652±0.002 | 0.309±0.007 |
| | chr14 | 2.5 | 28.26±0 | 23.98±0.01 | 4.27±0.01 | 0.13±0.001 | 0.288±0.002 | 0.582±0.002 | 0.849±0 | 0.708±0.002 |
| | | 5 | 29.34±0 | 24.31±0.02 | 5.03±0.02 | 0.144±0.001 | 0.315±0.001 | 0.541±0.002 | 0.829±0.001 | 0.663±0.002 |
| | | 10 | 29.93±0 | 23.96±0.03 | 5.96±0.03 | 0.165±0.002 | 0.337±0.003 | 0.498±0.003 | 0.801±0.001 | 0.6±0.003 |
| | | 20 | 30.24±0 | 23.09±0.05 | 7.15±0.05 | 0.187±0.003 | 0.354±0.002 | 0.459±0.004 | 0.764±0.002 | 0.519±0.005 |
| | | 40 | 30.41±0 | 22±0.05 | 8.41±0.05 | 0.209±0.003 | 0.366±0.003 | 0.425±0.003 | 0.724±0.001 | 0.434±0.005 |
| | | 50 | 30.45±0 | 21.52±0.06 | 8.93±0.06 | 0.218±0.003 | 0.37±0.001 | 0.412±0.004 | 0.707±0.002 | 0.399±0.005 |
| | | 60 | 30.47±0 | 21.19±0.14 | 9.28±0.14 | 0.22±0.003 | 0.371±0.003 | 0.409±0.003 | 0.695±0.005 | 0.379±0.011 |
| | | 80 | 30.5±0 | 20.65±0.07 | 9.85±0.07 | 0.229±0.001 | 0.374±0.001 | 0.397±0.003 | 0.677±0.002 | 0.343±0.004 |
| | | 100 | 30.52±0 | 20.17±0.09 | 10.35±0.09 | 0.237±0.001 | 0.375±0.003 | 0.388±0.002 | 0.661±0.003 | 0.311±0.006 |
| | | 200 | 30.56±0 | 18.91±0.15 | 11.65±0.15 | 0.259±0.004 | 0.389±0.003 | 0.352±0.006 | 0.619±0.005 | 0.235±0.009 |
| | chr15 | 2.5 | 20.08±0 | 16.89±0.01 | 3.19±0.01 | 0.135±0.001 | 0.304±0.002 | 0.561±0.003 | 0.841±0.001 | 0.702±0.002 |
| | | 5 | 20.92±0 | 17.18±0.02 | 3.74±0.02 | 0.15±0.001 | 0.326±0.002 | 0.524±0.003 | 0.821±0.001 | 0.659±0.002 |
| | | 10 | 21.39±0 | 16.98±0.03 | 4.41±0.03 | 0.168±0.003 | 0.345±0.001 | 0.487±0.004 | 0.794±0.002 | 0.599±0.004 |
| | | 20 | 21.65±0 | 16.45±0.02 | 5.2±0.02 | 0.187±0.003 | 0.355±0.003 | 0.459±0.003 | 0.76±0.001 | 0.528±0.003 |
| | | 40 | 21.8±0 | 15.76±0.04 | 6.04±0.04 | 0.206±0.003 | 0.363±0.003 | 0.431±0.002 | 0.723±0.002 | 0.452±0.005 |
| | | 50 | 21.83±0 | 15.42±0.05 | 6.4±0.05 | 0.215±0.003 | 0.363±0.002 | 0.422±0.005 | 0.707±0.002 | 0.419±0.006 |
| | | 60 | 21.85±0 | 15.13±0.07 | 6.72±0.07 | 0.223±0.003 | 0.362±0.004 | 0.414±0.005 | 0.693±0.003 | 0.39±0.007 |
| | | 80 | 21.88±0 | 14.81±0.07 | 7.07±0.07 | 0.228±0.004 | 0.364±0.004 | 0.407±0.006 | 0.677±0.003 | 0.362±0.007 |
| | | 100 | 21.9±0 | 14.46±0.08 | 7.44±0.08 | 0.24±0.002 | 0.367±0.002 | 0.393±0.003 | 0.66±0.004 | 0.328±0.007 |
| | | 200 | 21.93±0 | 13.59±0.15 | 8.35±0.15 | 0.262±0.005 | 0.369±0.005 | 0.369±0.004 | 0.62±0.007 | 0.252±0.011 |
| | chr16 | 2.5 | 11.47±0 | 9.66±0.01 | 1.81±0.01 | 0.14±0.002 | 0.29±0.001 | 0.571±0.002 | 0.842±0.001 | 0.692±0.002 |
| | | 5 | 11.92±0 | 9.72±0.01 | 2.2±0.01 | 0.16±0.002 | 0.321±0.001 | 0.519±0.003 | 0.816±0.001 | 0.632±0.002 |
| | | 10 | 12.16±0 | 9.49±0.04 | 2.68±0.04 | 0.183±0.005 | 0.344±0.003 | 0.473±0.004 | 0.78±0.003 | 0.554±0.009 |
| | | 20 | 12.3±0 | 9.04±0.03 | 3.26±0.03 | 0.211±0.002 | 0.353±0.003 | 0.436±0.003 | 0.735±0.003 | 0.456±0.004 |
| | | 40 | 12.37±0 | 8.51±0.07 | 3.87±0.07 | 0.239±0.005 | 0.358±0.004 | 0.403±0.009 | 0.688±0.005 | 0.355±0.011 |
| | | 50 | 12.39±0 | 8.36±0.04 | 4.03±0.04 | 0.248±0.003 | 0.364±0.003 | 0.387±0.005 | 0.675±0.003 | 0.328±0.006 |
| | | 60 | 12.4±0 | 8.23±0.03 | 4.17±0.03 | 0.254±0.006 | 0.362±0.003 | 0.384±0.007 | 0.664±0.003 | 0.306±0.007 |
| | | 80 | 12.42±0 | 8.01±0.05 | 4.41±0.05 | 0.26±0.005 | 0.366±0.004 | 0.373±0.007 | 0.645±0.004 | 0.274±0.007 |
| | | 100 | 12.42±0 | 7.84±0.06 | 4.59±0.06 | 0.271±0.005 | 0.364±0.001 | 0.365±0.005 | 0.631±0.005 | 0.244±0.01 |

| | | | | | | | | | | |
|---|---|---|---|---|---|---|---|---|---|---|
| | | 200 | 12.44±0 | 7.36±0.04 | 5.08±0.04 | 0.29±0.007 | 0.369±0.005 | 0.341±0.007 | 0.592±0.003 | 0.179±0.008 |
| | chr17 | 2.5 | 15.04±0 | 12.79±0.01 | 2.25±0.01 | 0.12±0.001 | 0.293±0.002 | 0.587±0.002 | 0.85±0 | 0.718±0.001 |
| | | 5 | 15.55±0 | 12.93±0.01 | 2.62±0.01 | 0.13±0.002 | 0.322±0.001 | 0.548±0.002 | 0.832±0.001 | 0.68±0.003 |
| | | 10 | 15.82±0 | 12.7±0.01 | 3.11±0.01 | 0.144±0.001 | 0.351±0.002 | 0.505±0.003 | 0.803±0.001 | 0.622±0.002 |
| | | 20 | 15.96±0 | 12.2±0.03 | 3.76±0.03 | 0.16±0.002 | 0.371±0.001 | 0.469±0.002 | 0.765±0.002 | 0.544±0.003 |
| | | 40 | 16.04±0 | 11.57±0.03 | 4.47±0.03 | 0.18±0.001 | 0.386±0.003 | 0.435±0.002 | 0.722±0.002 | 0.459±0.004 |
| | | 50 | 16.05±0 | 11.35±0.05 | 4.7±0.05 | 0.182±0.004 | 0.391±0.003 | 0.427±0.005 | 0.707±0.003 | 0.435±0.007 |
| | | 60 | 16.07±0 | 11.15±0.03 | 4.91±0.03 | 0.192±0.003 | 0.391±0.002 | 0.418±0.003 | 0.694±0.002 | 0.406±0.004 |
| | | 80 | 16.08±0 | 10.83±0.02 | 5.25±0.02 | 0.199±0.002 | 0.392±0.003 | 0.409±0.004 | 0.674±0.001 | 0.369±0.003 |
| | | 100 | 16.09±0 | 10.6±0.08 | 5.49±0.08 | 0.209±0.005 | 0.395±0.004 | 0.395±0.008 | 0.659±0.005 | 0.338±0.01 |
| | | 200 | 16.1±0 | 9.77±0.03 | 6.33±0.03 | 0.234±0.002 | 0.394±0.002 | 0.373±0.002 | 0.607±0.002 | 0.244±0.003 |
| | chr18 | 2.5 | 16.57±0 | 14.05±0.02 | 2.52±0.02 | 0.135±0.002 | 0.299±0.002 | 0.567±0.004 | 0.848±0.001 | 0.712±0.003 |
| | | 5 | 17.04±0 | 13.95±0.01 | 3.08±0.01 | 0.152±0.001 | 0.33±0.001 | 0.518±0.001 | 0.819±0.001 | 0.652±0.002 |
| | | 10 | 17.29±0 | 13.42±0.02 | 3.86±0.02 | 0.178±0.003 | 0.353±0.002 | 0.469±0.003 | 0.777±0.001 | 0.561±0.004 |
| | | 20 | 17.42±0 | 12.57±0.07 | 4.85±0.07 | 0.205±0.006 | 0.364±0.002 | 0.431±0.006 | 0.721±0.004 | 0.449±0.011 |
| | | 40 | 17.49±0 | 11.72±0.05 | 5.77±0.05 | 0.228±0.005 | 0.371±0.003 | 0.4±0.005 | 0.67±0.003 | 0.351±0.008 |
| | | 50 | 17.5±0 | 11.37±0.04 | 6.13±0.04 | 0.237±0.004 | 0.373±0.003 | 0.391±0.007 | 0.65±0.002 | 0.315±0.006 |
| | | 60 | 17.51±0 | 11.16±0.08 | 6.35±0.08 | 0.238±0.004 | 0.374±0.002 | 0.388±0.004 | 0.637±0.005 | 0.295±0.01 |
| | | 80 | 17.52±0 | 10.76±0.07 | 6.76±0.07 | 0.252±0.003 | 0.375±0.002 | 0.373±0.004 | 0.614±0.004 | 0.253±0.006 |
| | | 100 | 17.53±0 | 10.5±0.08 | 7.03±0.08 | 0.251±0.006 | 0.376±0.001 | 0.372±0.006 | 0.599±0.005 | 0.233±0.01 |
| | | 200 | 17.54±0 | 9.86±0.08 | 7.69±0.08 | 0.275±0.006 | 0.381±0.002 | 0.344±0.007 | 0.562±0.005 | 0.171±0.009 |

**Table S5. The accuracies (mean ± standard error) of GEBVs of different traits in Xu et al study based on SNP datasets across ten cross-validation replicates.**

| Trait[1] | GBLUP_CHIP | GBLUP_IMP[b] | GBLUP_PRUNE[2] |
|---|---|---|---|
| BW | 0.827 ± 0.003 | **0.831 ± 0.003 (↑)** | **0.831 ± 0.003 (↑)** |
| BL | 0.778 ± 0.005 | **0.779 ± 0.005 (↑)** | **0.774 ± 0.005 (↓)** |
| BF | 0.780 ± 0.005 | **0.782 ± 0.006 (↑)** | **0.788 ± 0.005 (↑)** |
| CC | 0.793 ± 0.004 | 0.793 ± 0.004 (=) | **0.801 ± 0.004 (↑)** |
| BH | 0.844 ± 0.003 | 0.840 ± 0.003 (↓) | 0.840 ± 0.003 (↓) |
| CW | 0.844 ± 0.003 | **0.842 ± 0.003 (↓)** | **0.847 ± 0.003 (↑)** |
| HW | 0.848 ± 0.004 | **0.852 ± 0.004 (↑)** | **0.852 ± 0.005 (↑)** |

[1]BW = body weight; BL = body length; BF = backfat thickness; BH = body height; CC = chest circumference; CW = chest width, HW = hip width.

[2]The symbols in parentheses represent that the prediction accuracy of GBLUP_IMP or GBLUP_PRUNE is higher than (↑ and bolded), lower than (↓) or equal to (=) the accuracy of GBLUP_CHIP.

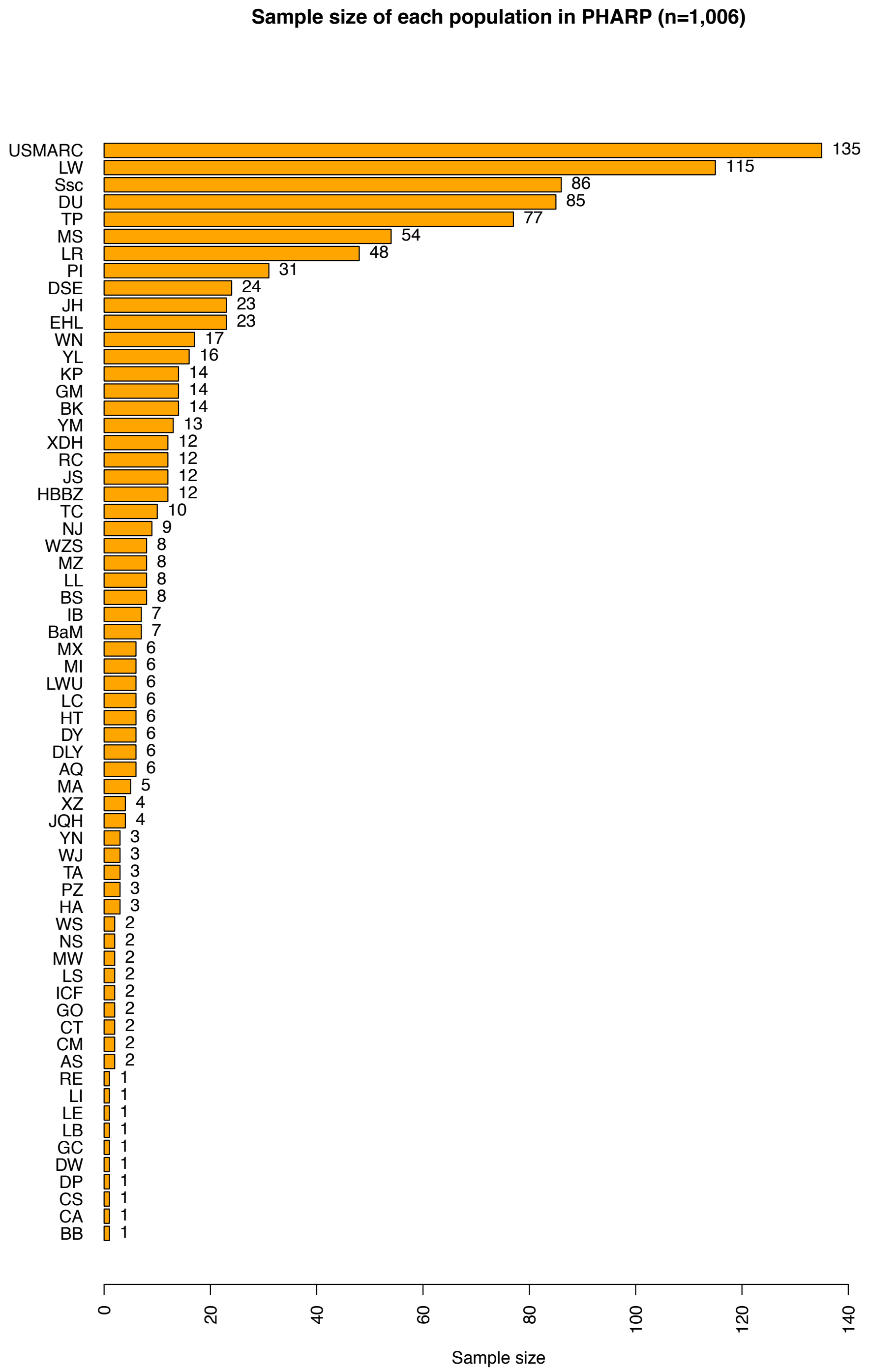

**Figure S1. Sample size of each population in PHARP.**

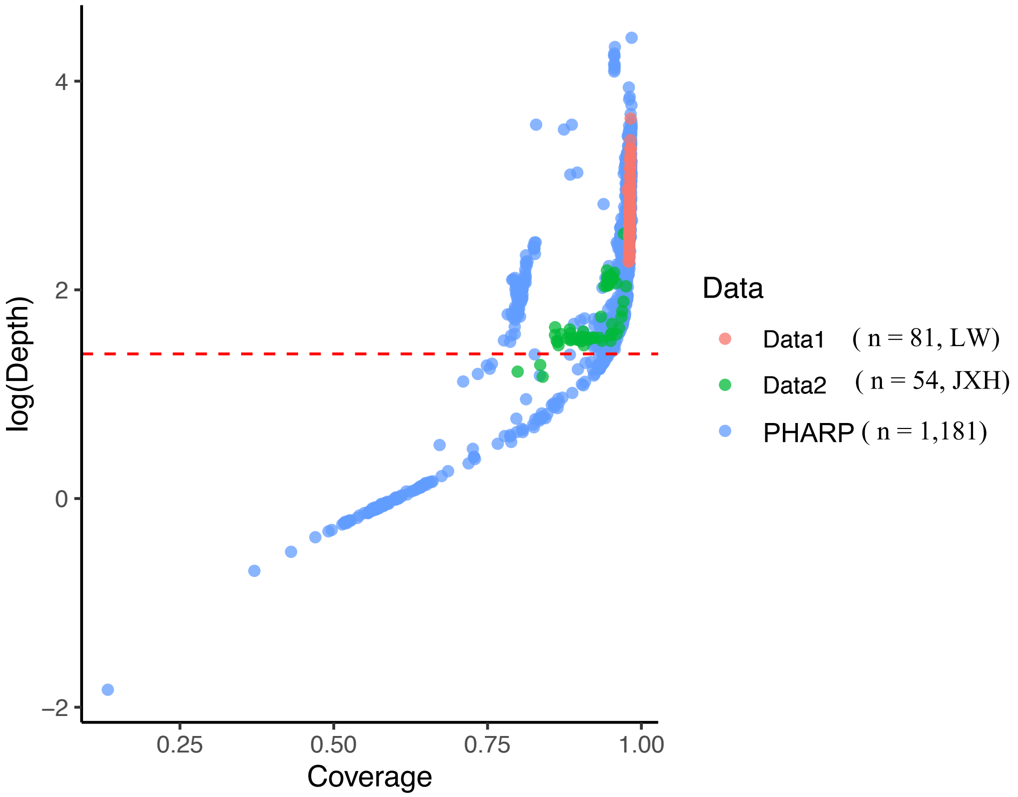

**Figure S2. The individual's coverage and depth distribution among the used datasets.** The red dashed line was a depth of 4X.

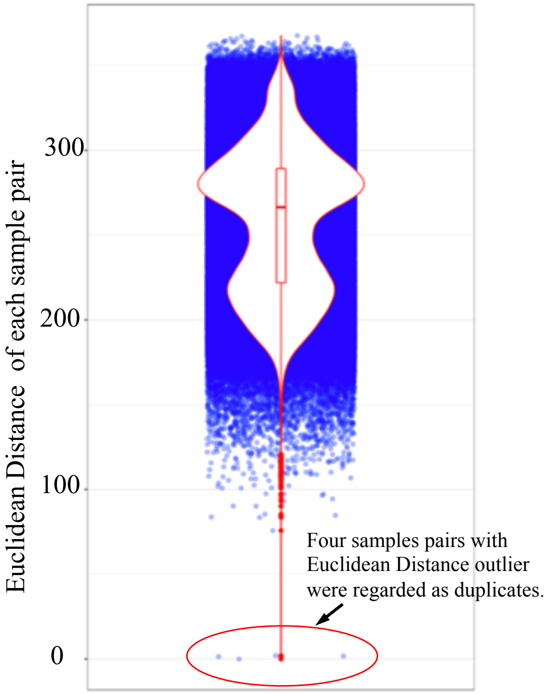

**Figure S3. The boxplot of the genetic distance measured by Euclidean distance for each sample pair.**

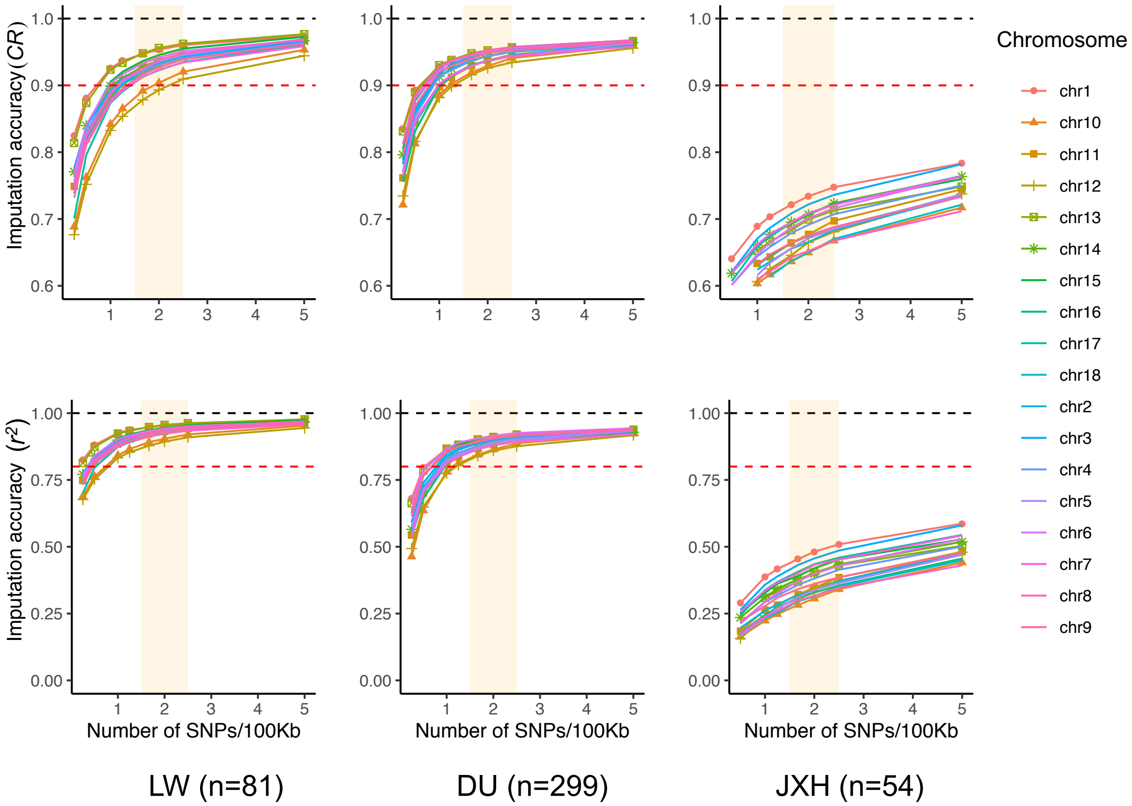

**Figure S4. Imputation accuracy estimated by mimicking the target panel with different densities (repeated 5 times) of SNPs on chromosomes using three test datasets (see more result detail in Table S4).**

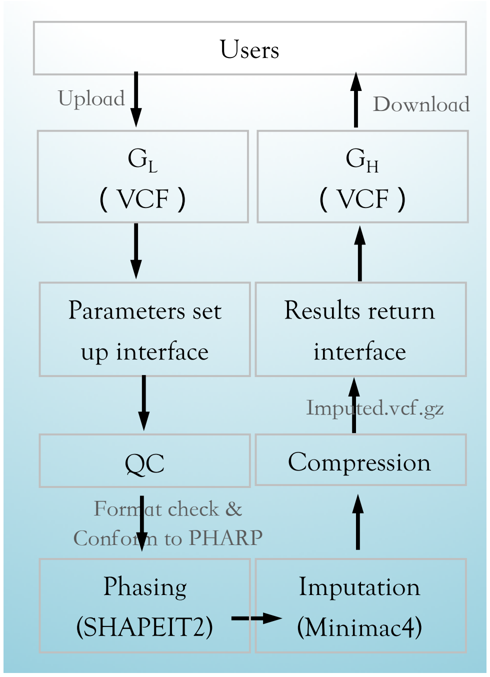

$\mathbf{G_L}$: Low-density genotypes ; $\mathbf{G_H}$: High-density genotypes

**Figure S5. The flow chart of the design consisting major steps for PHARP imputation server.**

**REFERENCE:**

1. Bosse, M., et al., *Regions of homozygosity in the porcine genome: consequence of demography and the recombination landscape*. PLoS Genet, 2012. **8**(11): p. e1003100.
2. Bosse, M., et al., *Genomic analysis reveals selection for Asian genes in European pigs following human-mediated introgression*. Nat Commun, 2014. **5**: p. 4392.
3. Rubin, C.J., et al., *Strong signatures of selection in the domestic pig genome*. Proceedings of the National Academy of Sciences of the United States of America, 2012. **109**(48): p. 19529-19536.
4. Groenen, M.A.M., et al., *Analyses of pig genomes provide insight into porcine demography and evolution*. Nature, 2012. **491**(7424): p. 393-398.
5. Fang, X., et al., *The sequence and analysis of a Chinese pig genome*. Gigascience, 2012. **1**(1): p. 16.
6. Vamathevan, J.J., et al., *Minipig and beagle animal model genomes aid species selection in pharmaceutical discovery and development*. Toxicology and Applied Pharmacology, 2013. **270**(2): p. 149-157.
7. Li, M.Z., et al., *Genomic analyses identify distinct patterns of selection in domesticated pigs and Tibetan wild boars*. Nature Genetics, 2013. **45**(12): p. 1431-U180.
8. Wang, C., et al., *Genome-wide analysis reveals artificial selection on coat colour and reproductive traits in Chinese domestic pigs*. Mol Ecol Resour, 2015. **15**(2): p. 414-24.
9. Choi, J.W., et al., *Whole-genome resequencing analyses of five pig breeds, including Korean wild and native, and three European origin breeds*. DNA Research, 2015. **22**(4): p. 259-267.
10. Moon, S., et al., *A genome-wide scan for signatures of directional selection in domesticated pigs*. BMC Genomics, 2015. **16**: p. 130.
11. Ai, H., et al., *Adaptation and possible ancient interspecies introgression in pigs identified by whole-genome sequencing*. Nat Genet, 2015. **47**(3): p. 217-25.
12. Li, M., et al., *Whole-genome sequencing of Berkshire (European native pig) provides insights into its origin and domestication*. Sci Rep, 2014. **4**: p. 4678.
13. Molnar, J., et al., *Genome sequencing and analysis of Mangalica, a fatty local pig of Hungary*. BMC Genomics, 2014. **15**: p. 761.
14. Ramirez, O., et al., *Genome data from a sixteenth century pig illuminate modern breed relationships*. Heredity (Edinb), 2015. **114**(2): p. 175-84.
15. Bosse, M., et al., *Using genome-wide measures of coancestry to maintain diversity and fitness in endangered and domestic pig populations*. Genome Res, 2015. **25**(7): p. 970-81.
16. Frantz, L.A., et al., *Evidence of long-term gene flow and selection during domestication from analyses of Eurasian wild and domestic pig genomes*. Nat Genet, 2015. **47**(10): p. 1141-8.
17. Jeong, H., et al., *Exploring evidence of positive selection reveals genetic basis of meat quality traits in Berkshire pigs through whole genome sequencing*. BMC Genet, 2015. **16**: p. 104.

18. Kim, J., et al., *Genome-wide detection and characterization of positive selection in Korean Native Black Pig from Jeju Island*. Bmc Genetics, 2015. **15**.
19. Revilla, M., et al., *A global analysis of CNVs in swine using whole genome sequence data and association analysis with fatty acid composition and growth traits*. Plos One, 2017. **12**(5).
20. Zhang, Y.B., et al., *Genome-wide identification of RNA editing in seven porcine tissues by matched DNA and RNA high-throughput sequencing*. Journal of Animal Science and Biotechnology, 2019. **10**.
21. Ma, Y.L., et al., *Genomic Analysis To Identify Signatures of Artificial Selection and Loci Associated with Important Economic Traits in Duroc Pigs*. G3-Genes Genomes Genetics, 2018. **8**(11): p. 3617-3625.
22. Reimer, C., et al., *Analysis of porcine body size variation using re-sequencing data of miniature and large pigs*. Bmc Genomics, 2018. **19**.
23. Keel, B.N., D.J. Nonneman, and G.A. Rohrer, *A survey of single nucleotide polymorphisms identified from whole-genome sequencing and their functional effect in the porcine genome*. Animal Genetics, 2017. **48**(4): p. 404-411.
24. Lu, M.D., et al., *Genetic variations associated with six-white-point coat pigmentation in Diannan small-ear pigs*. Scientific Reports, 2016. **6**.
25. Bianco, E., et al., *The chimerical genome of Isla del Coco feral pigs (Costa Rica), an isolated population since 1793 but with remarkable levels of diversity*. Molecular Ecology, 2015. **24**(10): p. 2364-2378.
26. Bianco, E., et al., *A Deep Catalog of Autosomal Single Nucleotide Variation in the Pig*. Plos One, 2015. **10**(3).
27. Funkhouser, S.A., et al., *Evidence for transcriptome-wide RNA editing among Sus scrofa PRE-1 SINE elements*. Bmc Genomics, 2017. **18**.
28. Zhao, P.J., et al., *Evidence of evolutionary history and selective sweeps in the genome of Meishan pig reveals its genetic and phenotypic characterization*. Gigascience, 2018. **7**(5).
29. Lin, Y., et al., *Genomic analyses provide insights into breed-of-origin effects from purebreds on three-way crossbred pigs*. Peerj, 2019. **7**.
30. Heckel, T., et al., *Functional analysis and transcriptional output of the Gottingen minipig genome*. Bmc Genomics, 2015. **16**.
31. Keel, B.N., et al., *A Survey of Copy Number Variation in the Porcine Genome Detected From Whole-Genome Sequence*. Frontiers in Genetics, 2019. **10**.
32. Li, M.Z., et al., *Comprehensive variation discovery and recovery of missing sequence in the pig genome using multiple de novo assemblies*. Genome Research, 2017. **27**(5): p. 865-874.
33. Falker-Gieske, C., et al., *GWAS for Meat and Carcass Traits Using Imputed Sequence Level Genotypes in Pooled F2-Designs in Pigs*. G3-Genes Genomes Genetics, 2019. **9**(9): p. 2823-2834.
34. Yan, G.R., et al., *Imputation-Based Whole-Genome Sequence Association Study Reveals Constant and Novel Loci for Hematological Traits in a Large-Scale Swine F-2 Resource Population*. Frontiers in Genetics, 2018. **9**.
35. Zhu, Y.L., et al., *Signatures of Selection and Interspecies Introgression in the Genome of Chinese Domestic Pigs*. Genome Biology and Evolution, 2017. **9**(10): p. 2592-2603.

36. Chen, H., et al., *Introgression of Eastern Chinese and Southern Chinese haplotypes contributes to the improvement of fertility and immunity in European modern pigs*. Gigascience, 2020. **9**(3).
37. Nosková, A., et al., *Infertility due to defective sperm flagella caused by an intronic deletion in DNAH17 that perturbs splicing*. Genetics, 2020. **217**(2).
38. Zhang, Z., et al., *Identifying the complex genetic architecture of growth and fatness traits in a Duroc pig population*. Journal of Integrative Agriculture, 2020. **19**(0).
39. Zhang, Z., et al., *Genome-Wide Association Study for Reproductive Traits in a Duroc Pig Population*. Animals (Basel), 2019. **9**(10).
40. Xu, P., et al., *Genome-wide association study for growth and fatness traits in Chinese Sujiang pigs*. Anim Genet, 2020. **51**(2): p. 314-318.
